# Genome-wide association studies identify 137 loci for DNA methylation biomarkers of ageing

**DOI:** 10.1101/2020.06.29.133702

**Authors:** Daniel L. McCartney, Josine L. Min, Rebecca C. Richmond, Ake T. Lu, Maria K. Sobczyk, Gail Davies, Linda Broer, Xiuqing Guo, Ayoung Jeong, Jeesun Jung, Silva Kasela, Seyma Katrinli, Pei-Lun Kuo, Pamela R. Matias-Garcia, Pashupati P. Mishra, Marianne Nygaard, Teemu Palviainen, Amit Patki, Laura M. Raffield, Scott M. Ratliff, Tom G. Richardson, Oliver Robinson, Mette Soerensen, Dianjianyi Sun, Pei-Chien Tsai, Matthijs D. van der Zee, Rosie M. Walker, Xiaochuan Wang, Yunzhang Wang, Rui Xia, Zongli Xu, Jie Yao, Wei Zhao, Adolfo Correa, Eric Boerwinkle, Pierre-Antoine Dugué, Peter Durda, Hannah R Elliott, Christian Gieger, The Genetics of DNA Methylation Consortium, Eco J.C. de Geus, Sarah E. Harris, Gibran Hemani, Medea Imboden, Mika Kähönen, Sharon L.R. Kardia, Jacob K. Kresovich, Shengxu Li, Kathryn L. Lunetta, Massimo Mangino, Dan Mason, Andrew M. McIntosh, Jonas Mengel-From, Ann Zenobia Moore, Joanne M. Murabito, NHLBI Trans-Omics for Precision Medicine (TOPMed) Consortium, Miina Ollikainen, James S. Pankow, Nancy L. Pedersen, Annette Peters, Silvia Polidoro, David J. Porteous, Olli Raitakari, Stephen S. Rich, Dale P. Sandler, Elina Sillanpää, Alicia K Smith, Melissa C. Southey, Konstantin Strauch, Hemant Tiwari, Toshiko Tanaka, Therese Tillin, Andre G Uitterlinden, David J. Van Den Berg, Jenny van Dongen, James G. Wilson, John Wright, Idil Yet, Donna Arnett, Stefania Bandinelli, Jordana T. Bell, Alexandra M Binder, Dorret I Boomsma, Wei Chen, Kaare Christensen, Karen N. Conneely, Paul Elliott, Luigi Ferrucci, Myriam Fornage, Sara Hägg, Caroline Hayward, Marguerite Irvin, Jaakko Kaprio, Deborah A. Lawlor, Terho Lehtimäki, Falk W. Lohoff, Lili Milani, Roger L. Milne, Nicole Probst-Hensch, Alex P. Reiner, Beate Ritz, Jerome I. Rotter, Jennifer A. Smith, Jack A. Taylor, Joyce B.J. van Meurs, Paolo Vineis, Melanie Waldenberger, Ian J. Deary, Caroline L. Relton, Steve Horvath, Riccardo E. Marioni

**Affiliations:** Centre for Genomic and Experimental Medicine, Institute of Genetics and Molecular Medicine, University of Edinburgh, Crewe Road South, Edinburgh, EH4 2XU, United Kingdom; MRC Integrative Epidemiology Unit at the University of Bristol, University of California Los Angeles, Los Angeles, CA 90095, USA; Population Health Science, Bristol Medical School, University of Bristol, University of California Los Angeles, Los Angeles, CA 90095, USA; Department of Human Genetics, David Geffen School of Medicine, University of California Los Angeles, Los Angeles, CA 90095, USA; Lothian Birth Cohorts, Department of Psychology, University of Edinburgh, Edinburgh, EH8 9JZ, UK; Department of Internal Medicine, Erasmus MC, Rotterdam, the Netherlands; The Institute for Translational Genomics and Population Sciences, Department of Pediatrics, The Lundquist Institute for Biomedical Innovation at Harbor-UCLA Medical Center, Torrance, CA USA; Swiss Tropical and Public Health Institute, Basel, Switzerland; University of Basel, Basel, Switzerland; National Institute on Alcohol Abuse and Alcoholism, National Institutes of Health, Bethesda, USA; Estonian Genome Center, Institute of Genomics, University of Tartu, Tartu, Estonia; Department of Gynecology and Obstetrics, Emory University School of Medicine, Atlanta, Georgia; Longitudinal Study Section, Translational Gerontology Branch, National Institute on Aging, Baltimore MD; Research Unit of Molecular Epidemiology, Helmholtz Zentrum München, German Research Center for Environmental Health, D-85764, Neuherberg, Bavaria, Germany; Institute of Epidemiology, Helmholtz Zentrum München, German Research Center for Environmental Health, D-85764, Neuherberg, Bavaria, Germany; TUM School of Medicine, Technical University of Munich, Munich, Germany; Department of Clinical Chemistry, Fimlab Laboratories, and Finnish Cardiovascular Research Center - Tampere, Faculty of Medicine and Health Technology, Tampere University, Tampere 33520, Finland; Epidemiology, Biostatistics and Biodemography, Department of Public Health, University of Southern Denmark, Odense, Denmark; Department of Clinical Genetics, Odense University Hospital, Odense, Denmark; Institute for Molecular Medicine Finland, FIMM, HiLife, University of Helsinki, Helsinki, Finland; Department of Biostatistics, University of Alabama at Birmingham; Department of Genetics, University of North Carolina at Chapel Hill, Chapel Hill, NC; Department of Epidemiology, School of Public Health, University of Michigan; MRC Centre for Environment and Health, School of Public Health, Imperial College London, UK; Department of Clinical Biochemistry and Pharmacology, Odense University Hospital, Odense, Denmark; Department of Epidemiology and Biostatistics, School of Public Health, Peking University Health Science Center,Beijing, China; Department of Twin Research and Genetic Epidemiology, King’s College London, UK; Department of Biomedical Sciences, Chang Gung University, Taoyuan, Taiwan; Genomic Medicine Core Laboratory, Chang Gung Memorial Hospital, Linkou, Taiwan; Department of Biological Psychology, Vrije Universiteit Amsterdam, Amsterdam, The Netherlands; Amsterdam Public Health Research Institute, Amsterdam, The Netherlands; Cancer Epidemiology Division, Cancer Council Victoria, 615 St Kilda Road, Melbourne, Victoria 3004, Australia; Department of Medical Epidemiology and Biostatistics, Karolinska Institutet; Brown Foundation Institute of Molecular Medicine, Mc Govern Medical School, University of Texas Health Science Center at Houston, Houston, TX; National Institute of Environmental Health Sciences, Research Triangle Park, NC 27709; Department of Medicine, University of Mississippi Medical Center, Jackson, MS; School of Public Health, University of Texas Health Science Center at Houston, Houston, TX; Precision Medicine, School of Clinical Sciences at Monash Health, Monash University, Clayton, Victoria 3168, Australia; Centre for Epidemiology and Biostatistics, Melbourne School of Population and Global Health, The University of Melbourne, 207 Bouverie Street, Melbourne, Victoria 3010, Australia; Department of Pathology & Laboratory Medicine, Larner College of Medicine, University of Vermont, Burlington, VT 05446, USA; Genetics of DNA methylation Consortium, Bristol; Department of Clinical Physiology, Tampere University Hospital, and Finnish Cardiovascular Research Center - Tampere, Faculty of Medicine and Health Technology, Tampere University, Tampere 33521, Finland; Children’s Minnesota Research Institute, Children’s Hospitals and Clinics of Minnesota, Minneapolis, MN 55404,USA; Department of Biostatistics, Boston University School of Public Health; NIHR Biomedical Research Centre at Guy’s and St Thomas’ Foundation Trust, London SE1 9RT, UK; Bradford Institute for Health Research, Bradford Teaching Hospitals NHS Foundation Trust, Bradford, UK; Division of Psychiatry, University of Edinburgh, Edinburgh, UK; Section of General Internal Medicine, Department of Medicine, Boston University School of Medicine, Boston, MA, USA; TOPMed Data Coordinating Center, Genetic Analysis Center, Department of Biostatistics, University of Washington; Division of Epidemiology and Community Health, University of Minnesota, Minneapolis, MN; German Center for Cardiovascular Research (DZHK), Partner Site Munich Heart Alliance, Munich, Germany; Centre for Population Health Research, University of Turku and Turku University Hospital, Turku, Finland; Research Centre of Applied and Preventive Cardiovascular Medicine, University of Turku, Turku, Finland; Department of Clinical Physiology and Nuclear Medicine, Turku University Hospital, Turku, Finland; Department of Public Health Sciences, Center for Public Health Genomics, University of Virginia, Charlottesville, VA 22908, USA; Gerontology Research Center, Faculty of Sport and Health Sciences, University of Jyväskylä, Jyväskylä, Finland; Department of Psychiatry and Behavioral Sciences, Emory University School of Medicine, Atlanta, Georgia; Institute of Genetic Epidemiology, Helmholtz Zentrum München, German Research Center for Environmental Health, D-85764, Neuherberg, Bavaria, Germany; Institute of Medical Biostatistics, Epidemiology and Informatics (IMBEI), University Medical Center, Johannes Gutenberg University, 55101 Mainz, Germany; Chair of Genetic Epidemiology, Institute for Medical Information Processing, Biometry, and Epidemiology, Faculty of Medicine, Ludwig-Maximilians-Universität München, Munich, Germany; MRC Unit for Lifelong Health and Ageing at UCL; Department of Epidemiology, Erasmus MC, Rotterdam, the Netherlands; Center for Genetic Epidemiology, Department of Preventive Medicine, Keck School of Medicine of USC, University of Southern California, Los Angeles, CA USA; Division of Cardiology, Beth Israel Deaconess Medical Center, Boston, MA; Department of Physiology and Biophysics, University of Mississippi Medical Center, Jackson, MS; Department of Bioinformatics, Institute of Health Sciences, Hacettepe University, 06100, Ankara, Turkey; Deans Office, College of Public Health, University of Kentucky; Geriatric Unit, Azienda Sanitaria Toscana Centro, Florence, Italy; Department of Epidemiology, Fielding School of Public Health, University of California, Los Angeles, CA, USA; Population Sciences in the Pacific Program (Cancer Epidemiology), University of Hawai□i Cancer Center, University of Hawai□i, Honolulu, HI, USA; Department of Epidemiology, Tulane University, New Orleans, LA 70112, USA; Department of Human Genetics, Emory University School of Medicine, Atlanta, Georgia; MRC Human Genetics Unit, Institute of Genetics and Molecular Medicine, University of Edinburgh, Crewe Rd. South, Edinburgh, EH4 2XU, United Kingdom; Dept of Epidemiology, University of Alabama at Birmingham; Department of Public Health, University of Helsinki, Helsinki, Finland; Bristol NIHR Biomedical Research Centre; Department of Epidemiology, University of Washington, Seattle, WA; Department of Biostatistics, Fielding School of Public Health, University of California Los Angeles, Los Angeles, California, 90095; USA

## Abstract

Biological ageing estimators derived from DNA methylation (DNAm) data are heritable and correlate with morbidity and mortality. Leveraging DNAm and SNP data from >41,000 individuals, we identify 137 genome-wide significant loci (113 novel) from meta-analyses of four epigenetic clocks and epigenetic surrogate markers for granulocyte proportions and plasminogen activator inhibitor 1 levels, respectively. We report strong genetic correlations with longevity and lifestyle factors such as smoking, education, and obesity. Significant associations are observed in polygenic risk score analysis and to a lesser extent in Mendelian randomization analyses. This study illuminates the genetic architecture underlying epigenetic ageing and its shared genetic contributions with lifestyle factors and longevity.

## Introduction

Ageing is associated with an increased risk of a multitude of physical, cognitive and degenerative disorders [1]. While the rate of chronological ageing is constant between individuals, there are inter-individual differences in the risk of age-associated morbidities. Biological ageing is influenced by both environmental and genetic factors [2]. Multiple measures of biological age exist, several of which have drawn information from DNA methylation (DNAm) across the genome. DNAm is a common epigenetic modification typically characterised by the addition of a methyl group to a cytosine-guanine dinucleotide (CpG). DNAm levels can be influenced by both genetic and environmental factors, and in recent years, DNAm signatures have become established correlates of multiple health-related outcomes [3, 4, 5]. Such signatures include “epigenetic clocks”, accurate markers of ageing which associate with several health outcomes [6, 7]. Epigenetic clocks use weighted linear combinations of CpGs to predict an individual’s chronological age and have common SNP-based heritability estimates ranging from 0.15 to 0.19 [8, 9]. Individuals with epigenetic clock estimates greater than their chronological age display “age acceleration”, and have been shown to be at a greater risk of all-cause mortality and multiple adverse health outcomes [10].

The first generation of epigenetic ageing clocks used penalized regression models to predict chronological age on the basis of DNA methylation data, e.g. the widely used clocks from Hannum (2013) and Horvath (2013) apply to blood and 51 human tissues/cell types, respectively [11, 12, 13]. A derivative of the Horvath clock, intrinsic epigenetic age acceleration (IEAA) has since been developed, conditioning out (i.e. removing) estimates of blood cell composition. An increasing literature supports the view that IEAA relates to properties of hematopoietic stem cells [8, 2, 14]. The second generation of epigenetic clocks move beyond estimating chronological age by incorporating information on morbidity and mortality risk (e.g., smoking, plasma protein levels, white blood cell counts), and chronological age. Two such predictors, termed PhenoAge (a DNAm predictor trained on a measure that itself was trained on mortality, using 42 clinical measures and age as input features) and GrimAge (trained on mortality, including a DNAm measure of smoking as a constituent part), outperform both Hannum and Horvath clocks in predicting mortality, and are associated with various measures of morbidity and lifestyle factors [15, 16]. DNAm GrimAge outperforms PhenoAge and the first generation of epigenetic clocks when it comes to predicting time to death [8, 17, 18].

While nothing is known about the genetics of the second generation of epigenetic clocks, 13 genetic loci have been associated with the first generation of epigenetic clocks. A study of nearly ten thousand individuals revealed a regulatory relationship between human telomerase (hTERT) and epigenetic age acceleration [8]. More recently, a larger GWAS (n=13,493) revealed that metabolic and immune pathways share genetic underpinnings with epigenetic clocks [9].

Here, we greatly expand on these studies across several dimensions. First, we analyse a large, multi-ethnic dataset comprised of over 41k individuals from 29 European ancestry studies, seven African American studies, and one Hispanic ancestry study. Second, we characterize for the first time the genetic architecture of the second generation epigenetic clocks, GrimAge and PhenoAge. Third, we also conduct GWAS of two important DNAm based surrogate markers: DNAm plasminogen activator inhibitor-1 (PAI1) levels and granulocyte proportion, respectively. DNAmPAI1 was chosen because it exhibited stronger associations with cardiometabolic disease than the epigenetic clocks [15]. The DNAm based estimate of granulocyte proportions was chosen because a) it exhibited significant associations with several epigenetic clocks (including GrimAge and PhenoAge) and with health outcomes such as Parkinson’s disease [15, 16, 19]. The unprecedented sample size of the current study allowed us to develop polygenic risk scores for these six epigenetic biomarkers.

We report 137 independent loci, including 113 novel loci (i.e. not previously identified in previous GWAS meta-analyses of epigenetic age estimators [8, 9]), and examine the genetic and causal relationships between epigenetic ageing, lifestyle behaviours, health outcomes, and longevity.

## Results

To identify genetic variants associated with six methylation-based biomarkers, genome wide association studies (GWAS) of 34,962 European ancestry and 6,482 African American individuals were performed (**Appendix 1** and **Tables S1-S2**). A fixed-effects meta-analysis was performed to combine the summary statistics within each ancestry group. Genomic inflation factors ranged between 1.01 and 1.06 (**Figure S1-S6**; https://datashare.is.ed.ac.uk/handle/10283/3645) for the European-only meta-analyses and from 1.11 and 1.21 for the meta-analyses comprising African American participants (**Figures S7-S12; Table 1**). Inflation was present and consistent across all allele frequencies in the African American analyses; there was much greater variability in the effect sizes in the African American analyses (**Appendix 2**). We examined the relationship between means and standard deviations of predicted age versus means and standard deviations of chronological age for each cohort, separated by ancestry group, observing weaker mean correlations in the African American cohorts. There was little difference in the relationship between the standard deviations of age acceleration and chronological age by ancestry group (**Figure S13**). Heterogeneity between studies may decrease power to detect genetic associations. We found little evidence of systematic between-study heterogeneity in both the European ancestry and African American meta-analyses, as determined by M-statistic outlier analysis, meta-regressions against cohort characteristics, and analysis of heterogeneity I^2^ statistics [21] (**Appendix 2**).

**Table 1:**
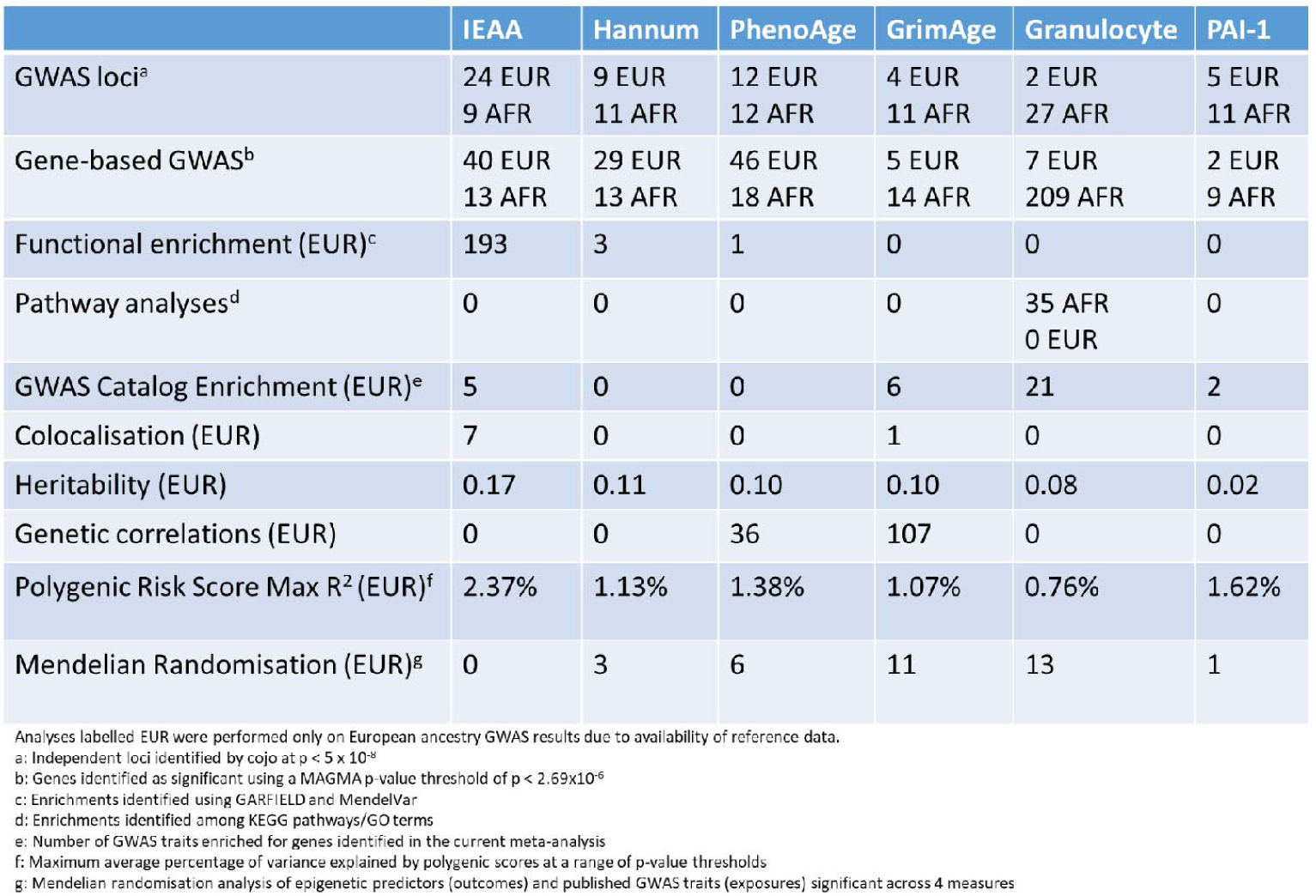
Summary of key findings of post GWA analyses. African American meta-analyses based on 6,148 participants from 7 cohorts; European ancestry meta-analyses based on 34,449 participants from 28 cohorts.

|  | IEAA | Hannum | PhenoAge | GrimAge | Granulocyte | PAI-1 |
| --- | --- | --- | --- | --- | --- | --- |
| GWAS loci<sup>a</sup> | 24 EUR<br>9 AFR | 9 EUR<br>11 AFR | 12 EUR<br>12 AFR | 4 EUR<br>11 AFR | 2 EUR<br>27 AFR | 5 EUR<br>11 AFR |
| Gene-based GWAS<sup>b</sup> | 40 EUR<br>13 AFR | 29 EUR<br>13 AFR | 46 EUR<br>18 AFR | 5 EUR<br>14 AFR | 7 EUR<br>209 AFR | 2 EUR<br>9 AFR |
| Functional enrichment (EUR)<sup>c</sup> | 193 | 3 | 1 | 0 | 0 | 0 |
| Pathway analyses<sup>d</sup> | 0 | 0 | 0 | 0 | 35 AFR<br>0 EUR | 0 |
| GWAS Catalog Enrichment (EUR)<sup>e</sup> | 5 | 0 | 0 | 6 | 21 | 2 |
| Colocalisation (EUR) | 7 | 0 | 0 | 1 | 0 | 0 |
| Heritability (EUR) | 0.17 | 0.11 | 0.10 | 0.10 | 0.08 | 0.02 |
| Genetic correlations (EUR) | 0 | 0 | 36 | 107 | 0 | 0 |
| Polygenic Risk Score Max R<sup>2</sup> (EUR)<sup>f</sup> | 2.37% | 1.13% | 1.38% | 1.07% | 0.76% | 1.62% |
| Mendelian Randomisation (EUR)<sup>g</sup> | 0 | 3 | 6 | 11 | 13 | 1 |

Analyses labelled EUR were performed only on European ancestry GWAS results due to availability of reference data.
a: Independent loci identified by cojo at $p < 5 \times 10^{-8}$
b: Genes identified as significant using a MAGMA p-value threshold of $p < 2.69 \times 10^{-6}$
c: Enrichments identified using GARFIELD and MendelVar
d: Enrichments identified among KEGG pathways/GO terms
e: Number of GWAS traits enriched for genes identified in the current meta-analysis
f: Maximum average percentage of variance explained by polygenic scores at a range of p-value thresholds
g: Mendelian randomisation analysis of epigenetic predictors (outcomes) and published GWAS traits (exposures) significant across 4 measures

The key findings of the post GWA analyses are summarised in **Table 1**.

### European ancestry GWAS meta-analysis: 56 independently associated loci

We identified 56 conditionally independent associations (P <5×10^−8^) across the six epigenetic biomarkers in European ancestry populations using a stepwise model (**Figures S14-S19; Table 1**; **Table S3**; [22, 23]. We replicated 10/10 loci associated with IEAA and 1/1 locus associated with a cell-adjusted Hannum based measure of epigenetic age acceleration identified in an earlier GWAS (P < 0.05/11 = 0.0045) [9]. All but three loci (associated with IEAA) were replicated at the genome-wide significant level. To validate the associations with DNAm derived granulocyte counts, we compared our results to a previous GWAS of FACS granulocyte counts which identified 155 independent loci [20]. In the current meta-analysis, we replicated 13/129 present loci (P < 0.05/129 = 3.88×10^−4^; **Table S4; Figures S20-S21**), two of which replicated at the genome-wide significant level. Effect sizes at the 129 loci were strongly correlated between studies (r=0.85).

To examine whether genetic variation across the six epigenetic biomarkers was shared, we performed genetic colocalization analyses of the 56 loci [24]. There was evidence for colocalization at 29 loci between epigenetic biomarkers (posterior probability > 0.8; **Table S5**). IEAA was associated with the greatest number of independent associations (n=24), whereas granulocyte proportion was associated with the fewest (n=2). The most significant independent association was observed between IEAA and rs10949481 (β_T_allele_ = -0.20; P=4.5×10^−54^), mapping to the 3’UTR of *NHLRC1* on chromosome 6p22.3, within one of the top loci identified in previous meta-analyses of IEAA [8, 9].

### African American GWAS meta-analysis: 81 independently associated loci

We identified 81 conditionally independent associations for the six epigenetic biomarkers in the African American analyses (**Figures S22-S27; Table 1; Table S3**). The number of associated loci per epigenetic biomarker ranged from 9 (IEAA) to 27 (granulocyte proportion). The most significant association was observed between granulocyte proportions and rs2814778 (β_T_allele_ = -1.24; P =1.93×10^−83^), in the 5’ UTR of *ACKR1* on chromosome 1.

### Trans-ethnic meta-analyses identify 69 loci

To determine if any loci were shared across the European and African American populations, a trans-ethnic meta-analysis was carried out for each of the six epigenetic biomarkers using MRMEGA [25]. There were between 6 and 23 risk loci common to all ancestries (69 loci in total). Fifty-three of these loci contained independent lead SNPs from the ancestry-specific meta-analyses (38 and 15 lead SNPs from the European ancestry and African American meta-analyses were contained within these loci, respectively); 21 of the trans-ethnic loci did not contain a SNP identified in the ancestry-specific meta-analysis models (**Table S6**). We found 21 ancestry-specific lead SNPs (i.e. those identified in either European ancestry or African American analyses and not within the trans-ethnic loci; **Table S7)**. Allele frequencies for 11/21 ancestry-specific lead SNP allele frequencies differed by >10%.

We compared effect sizes of the lead SNPs from these loci in a Hispanic-American ancestry subset of the MESA cohort (n=287). Correlations between the respective effect sizes ranged from 0.16 (granulocyte proportions) to 0.92 (Hannum age acceleration; **Table S8** and **Figure S28**).

### Gene Based GWAS identifies 364 significant genes

Gene-based GWASs (carried out using MAGMA) identified between two and 46 genes (111 unique genes in total) associated with the six epigenetic biomarkers (**Table S9**) in the European ancestry data [26]. These included associations between IEAA and *TERT* (P=9.13×10^−12^), *TRIM59* (P=4.06×10^−12^), and *LRRC16A* (P=2.83×10^−8^) as previously reported [8, 9]. *TERT* and *TRIM59* were also associated with both Hannum age acceleration (P=5.44×10^−8^ and 9.92×10^−8^) and PhenoAge acceleration (P=2.07×10^−7^ and 2.66×10^−6^), respectively. In the African American data, between nine and 209 genes (264 unique genes in total) were associated with the epigenetic biomarkers, including *TMPT* (associated with PhenoAge acceleration and IEAA; P<2.39×10^−9^), with the most significant association observed between *CREB3L3* and PhenoAge acceleration (P=2.56×10^−53^; **Table S9**). Across all epigenetic biomarkers and ancestries there were 364 unique gene-based associations.

### Independently-associated loci are associated with DNA methylation levels

To explore whether any of the 56 loci from the European GWAS shared genetic variation influencing epigenetic clock DNAm sites, colocalization analyses using GoDMC summary statistics were conducted (**Methods**). We found strong evidence (posterior probability > 0.8) that 1/4 loci (25%) for GrimAge acceleration, 3/12 loci (25%) for PhenoAge acceleration, 11/24 loci (46%) for IEAA and 5/9 loci (56%) for Hannum age acceleration had shared genetic variation influencing epigenetic clock DNAm sites (**Tables S10-S11)**. Next, we used genetic colocalization to evaluate whether the DNAm derived phenotypes capturing genetic variation influencing DNAm sites associated with BMI [27] or smoking [28]; 29/56 loci (52%) colocalized with genetic variation influencing smoking associated DNAm sites and 1/56 loci (1.8%) was colocalized with genetic variation influencing a BMI-associated DNAm site. Specifically, GrimAge acceleration (78%) and Hannum age acceleration (75%) loci showed a large overlap of genetic variation influencing smoking DNAm sites.

Utilising results from a published GWAS of IEAA in brain tissue [29], we tested whether genetic variation influencing IEAA in blood and brain was shared for the 24 blood-related IEAA loci (cross-tissue plot for lead SNPs shown in **Figure S29**). Colocalization analysis showed that there was no strong evidence (PP > 0.80) for a single SNP being associated with both traits. However, the true extent of sharing is difficult to estimate because the sample size of the brain study (n=1796) is much smaller than our blood-based study, limiting power to detect shared loci. Previous simulations using a sample size of 2000 individuals have indicated that the shared variant must explain close to 2% of the variance of a biomarker to attain a posterior probability >0.8 for shared genetic effects [24]. Nevertheless, we observed suggestive evidence for colocalization (PP=0.53; LocusZoom plot in **Figure S30**) for a locus mapping to *DSCR6* on chromosome 21. This locus also overlaps with a mQTL for an IEAA clock CpG, cg13450409 (PP = 0.99; **Table S11**).

### Functional enrichment analysis

To gain an understanding of the functional and regulatory properties of the variants that underlie the six epigenetic biomarkers, we performed functional enrichment analyses across various gene annotations and regulatory and cell-type specific elements on the summary statistics for each of the European ancestry GWAS results (see **Methods, Table S12**) [30]. At an epigenetic biomarker-specific adjusted p-value calculated from the effective number of annotations, significant enrichments were present for IEAA (n=191) and Hannum age acceleration (n=3). Associations with IEAA were enriched in DHS hotspots in several tissues (which might reflect that Horvath’s pan tissue clock applies to all tissues) but the strongest enrichment of associatios with IEAA could be observed for mobilized CD34 primary cells (OR=6.06, P=6.1×10^−12^), which supports the view that IEAA reflects properties of hematopoietic stem cells.

### Pathway enrichment analysis

In the African American analysis, genes associated with granulocyte proportions were enriched amongst 35 Gene Ontology (GO) terms (Bonferroni P<0.05), the majority of which were immune-related (e.g. adaptive immune response, lymphocyte activation, regulation of immune system process or skin development; **Table S13**). By contrast, there was no significant enrichment amongst KEGG pathways or GO terms for the significantly-associated genes in the European ancestry analysis.

### Overlap with Mendelian disease genes

In order to characterise the potential overlap of Mendelian disease genes and associated pathways with our findings, a series of enrichment analyses were conducted (**Methods**) [31]. In the European ancestry analysis, enrichment of Mendelian disease genes was observed for IEAA and two gene-sets (‘disorders of platelet function’ and ‘vascular skin abnormality’: bootstrapped P=0.027 and 0.049, respectively). Enrichment analysis of PhenoAge loci revealed overrepresentation of methylation-related Mendelian disease genes (**Table S14;** bootstrapped *p*-value = 0.04) with *MTR* (methyltetrahydrofolate-homocysteine S-methyltransferase) present in addition to *TPMT*. No significant over-representations of Mendelian disease gene sets were observed for any of the genes identified in the African American analysis.

### SNP- and gene-based enrichment within published GWAS

To determine whether any of the 56 lead SNPs in the European ancestry meta-analyses for the six epigenetic biomarkers showed evidence for pleiotropic associations, a look-up of published GWAS significant associations (P <5×10^−8^) was carried out (**Table S15**). Four of the SNPs (rs2736100 in *TERT*, rs2275558 in *PBX1*, rs144317085 in *TET2* and rs2492286 in *RPN1*) were associated with 16 unique traits including multiple cancers (e.g. lung cancer, glioma) [32, 33] and blood cell counts (e.g. platelet count, eosinophil count, red blood cell count) [20, 32] (**Table S15**). All four SNPs were associated with IEAA in the current study. A gene-based test of enrichment among traits within the GWAS catalog output (**Table S16**) showed genes associated with IEAA, GrimAge and granulocyte proportion were enriched among those associated with white blood cell counts [20, 34]. Several genes associated with granulocyte proportions were also enriched among those associated with inflammatory traits (e.g. inflammatory bowel disease, rheumatoid arthritis, asthma) [35, 36, 37, 38, 39, 40]. IEAA-associated genes were also significantly enriched among those identified in a previous GWAS of IEAA [8, 9].

### Colocalization to identify GWAS SNPs that might regulate expression levels

Colocalization analyses were conducted to investigate whether any of the 56 loci from the European ancestry GWASs showed evidence of regulating gene expression levels (**Methods**). There was strong evidence (posterior probability > 0.8) that eight loci had shared genetic effects with eQTLs (**Table 1**; **Table S17**). Of these, one was associated with GrimAge acceleration and seven were associated with IEAA. The locus associated with GrimAge acceleration was linked to the expression of *C6orf183* whereas IEAA-associated loci were linked to the expression of 11 transcripts including genes related to lipid transport and immune function (e.g. *ATP8B4, CD46, TRIM59*).

### Heritability and LD Score Regression

We quantified the proportion of variance in the six epigenetic biomarkers from the European ancestry meta-analyses that can be explained by our SNP sets using LD Score regression [41]. The GWAS summary statistic SNP-based heritability ranged from 0.02 (SE = 0.02) for PAI1 levels, to 0.17 (SE = 0.02) for IEAA (**Table 1; Figure S31**; **Table S18**). We omitted DNAm PAI1 from the genetic correlation analysis due to its low heritability estimate. Several of the remaining five epigenetic biomarkers exhibited significant (p<0.05) pairwise genetic correlation coefficients ranging from r=0.27 (GrimAge acceleration and IEAA) to r=0.66 (GrimAge acceleration and PhenoAge acceleration; **Figure 1; Table S19**). A selection of significant genetic correlations is presented in **Figure 2**.

**Figure 1.**
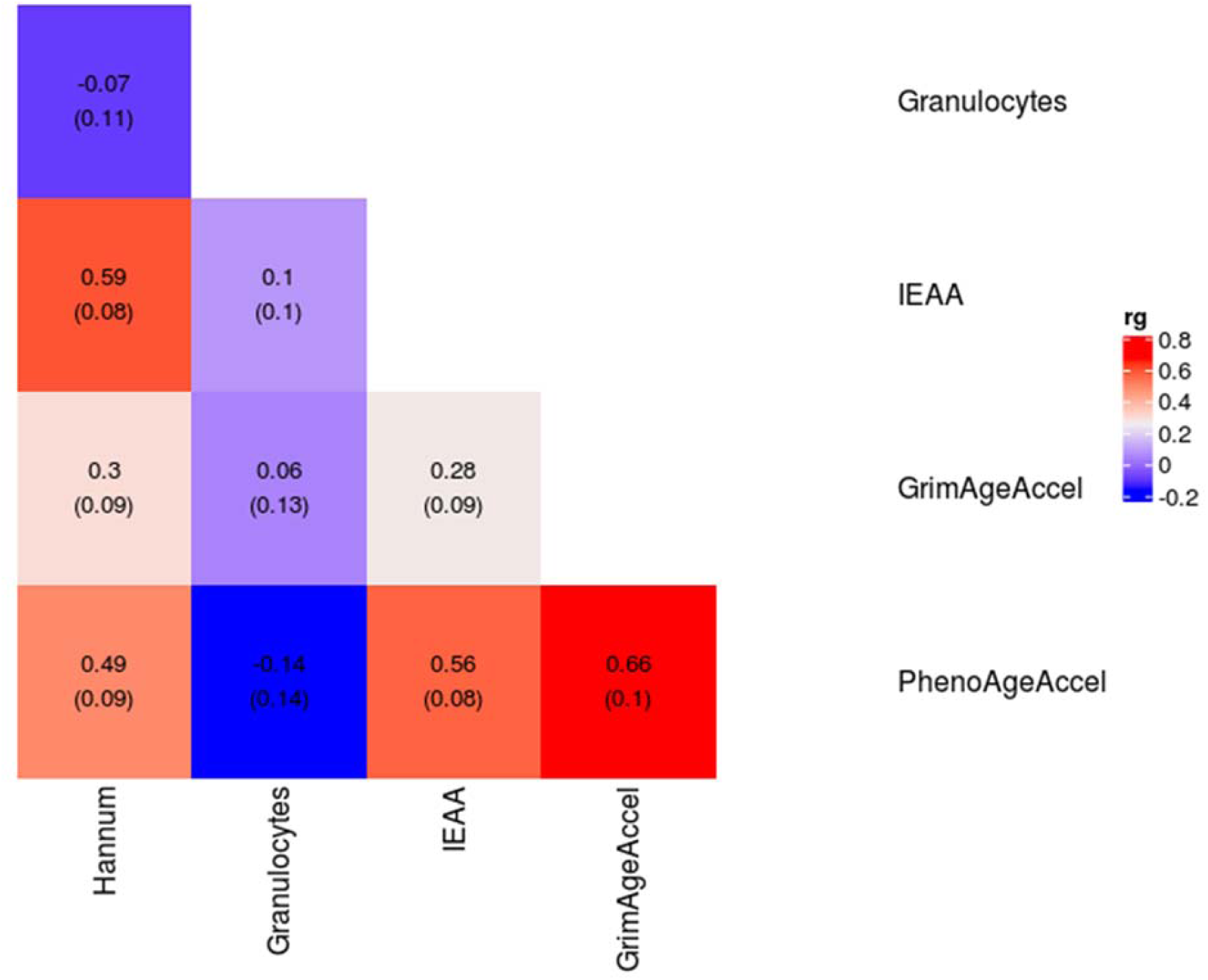
Genetic correlations and standard errors for the six epigenetic biomarkers. IEAA (Intrinsic Epigenetic Age Acceleration)

**Figure 2:**
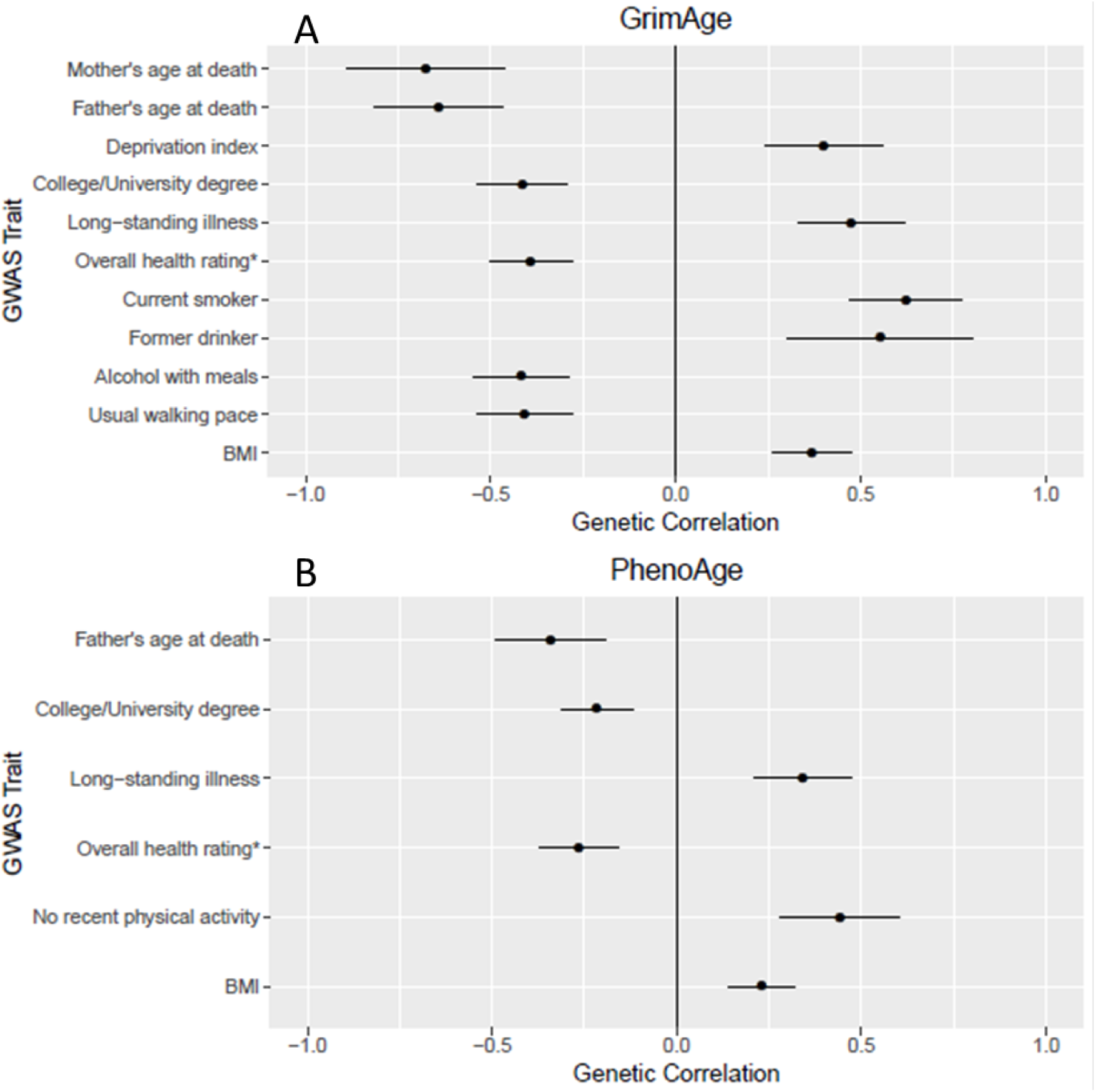
Significant genetic correlations betweenGrimAge acceleration (Panel A) and PhenoAge acceleration (Panel B) and a selection of GWAS traits. *This variable was originally coded with a high score representing lower health rating. We have multiplied the genetic correlation by -1 for interpretability.

PhenoAge acceleration had significant genetic correlations with 36 (out of 693) health-related traits including educational and cognitive traits (e.g. years of schooling, intelligence; r_g_ = -0.26 and -0.30; P≤ 3.29×10^−5^) [42, 43, 44], anthropometric traits (e.g. waist circumference, obesity, extreme BMI, hip circumference; r_g_ = 0.22-0.31; P ≤ 3.61×10^−5^)[45, 46, 47], adiposity (e.g. leg, arm, and trunk fat mass; r_g_ = 0.22-0.23; P ≤ 3.46×10^−6^) [47], and longevity (e.g. father’s age at death: r_g_ = -0.34; P = 9.66×10^−6^) [47]. GrimAge acceleration was genetically correlated with similar traits to PhenoAge (e.g. father’s age at death: r_g_ = -0.64; P = 6.2×10^−13^), along with smoking-related traits (e.g. current tobacco smoking: r_g_ = 0.62; P = 1.5 x10^−15^) [47] and cancer-related traits (e.g. lung cancer: r_g_ = 0.48; P = 8.3×10^−6^) [48]. The shared genetic contributions to GrimAge and smoking/mortality are expected given that GrimAge uses a DNAm based estimator of smoking pack-years in its definition. There were no significant genetic correlations between Hannum age acceleration, IEAA, or granulocyte proportions and any of the traits tested after correction for multiple testing (P < 0.05/693 = 7.22×10^−5^; **Table S20**).

### Polygenic risk score (PRS) profiling

To determine how well SNP-based genetic scores can approximate the six epigenetic biomarkers, and to see if these genetic scores associate with health outcomes, a polygenic risk score analysis was conducted on the European ancestry data. Re-running the meta-analysis with an iterative leave-one-cohort-out process (and on the full summary statistics in a completely independent cohort – the Young Finns Study), the mean polygenic predictions explained between 0.21 and 2.37% of the epigenetic biomarkers (**Table 1**; **Figure 3; Table S21**). The maximum prediction for a single cohort was 4.21% for PAI1 levels in ARIES. Parsimonious predictors (built using SNPs with P <5×10^−8^) performed well for IEAA, PAI1 levels and PhenoAge acceleration, whereas predictors including more SNPs (P <0.01 – P<1) tended to explain the most variance in GrimAge acceleration and granulocyte proportions.

**Figure 3.**
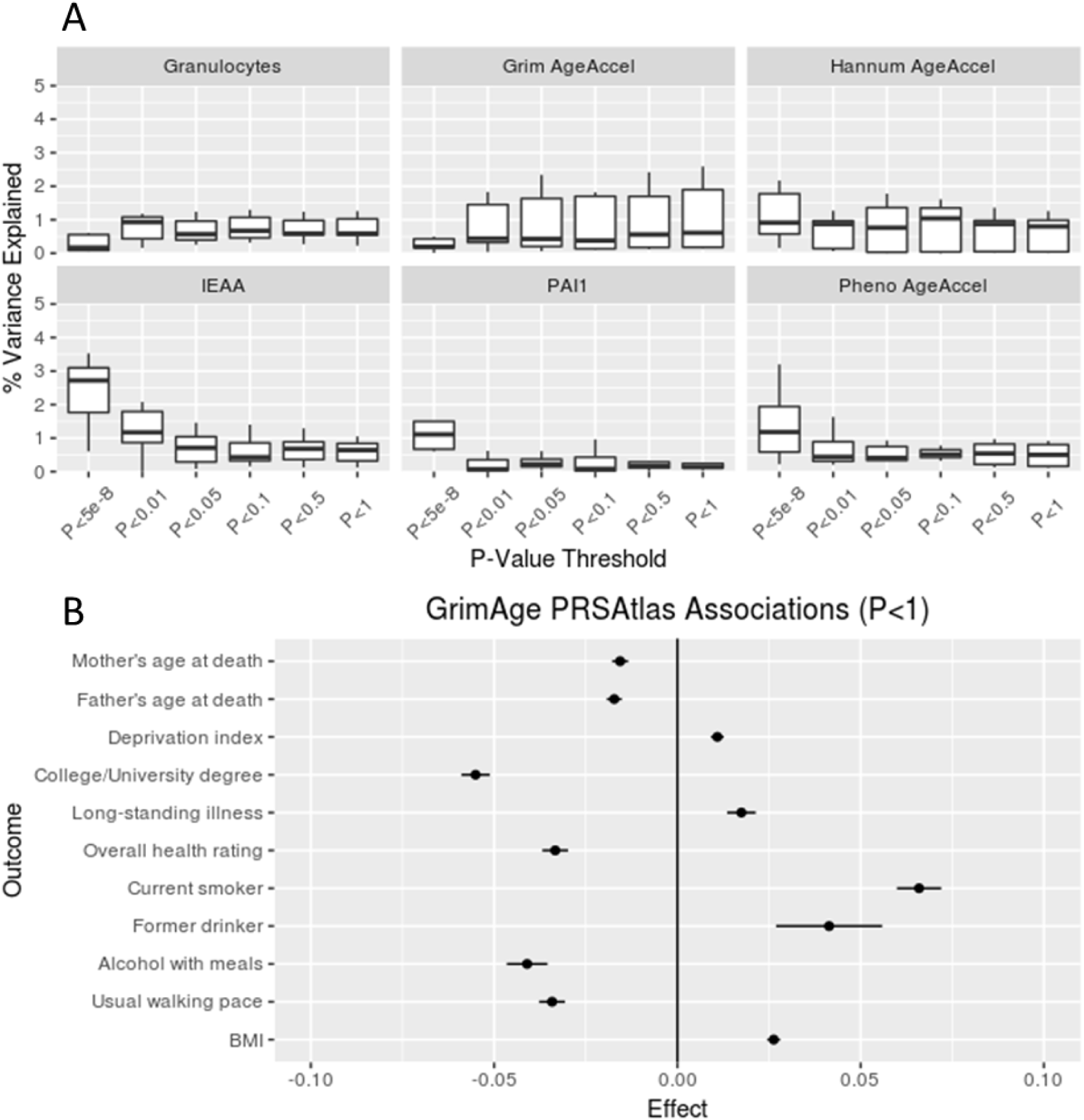
Panel A: Polygenic predictions for the six epigenetic biomarkers in LBC1921, LBC1936, SABRE, Born in Bradford, ARIES, FHS, and the Young Finns Study. IEAA (Intrinsic Epigenetic Age Acceleration), PAI1 (Plasminogen activator inhibitor-1). Panel B: Associations between GrimAge polygenic risk score (P<1) and UK Biobank GWAS traits.

In order to investigate the association between the polygenic risk scores and health outcomes, we utilised a PRS Atlas to model associations with 581 heritable traits (n_range_=10,299 to 334,915) from the UK Biobank study [49]. The PRS inputs included the independent SNPs with P<1 for GrimAge acceleration and granulocyte proportion and P<5×10^−8^ for the other four epigenetic biomarkers, with thresholds based on the results from the leave-one-out predictions (**Figure 3**). Using an FDR-corrected p-value for each of the six epigenetic biomarkers, we found between 7 and 250 significant associations for GrimAge acceleration, Granulocyte proportions and Hannum age acceleration (P_FDR_ < 0.05; **Table S22**). The strongest associations were between the GrimAge acceleration PRS and the following traits: adiposity-related traits (e.g. body fat percentage: β = 0.02; P_FDR_ = 7.3×10^−39^); education (e.g. college or university degree: β = -0.06; P_FDR_ = 2.6×10^−43^); and parental longevity (e.g. father’s age at death: β = -0.02; P_FDR_ = 5.7×10^−16^; mother’s age at death: β = -0.02; P_FDR_ = 1.6×10^−11^). Higher C-reactive protein was associated with a higher PRS for both granulocyte proportions and GrimAge acceleration (granulocyte proportions: β = 0.01; P_FDR_ = 8.2×10^−4^; GrimAge acceleration: β = 0.02; P_FDR_ = 2.1×10^−29^), and a lower score for Hannum age acceleration (β = -0.006; P_FDR_ = 0.02). A higher Hannum age acceleration PRS was also associated with an increased likelihood of taking insulin medication and lower total protein levels.

### Mendelian Randomisation between age acceleration phenotypes and health and life style outcomes

To investigate if the epigenetic measures were causally influenced by lifestyle factors and had a causal effect on ageing and disease outcomes, we performed Mendelian randomization (MR) analyses on 150 traits for the European ancestry data (**Table S23)**. We found 12 inverse-variance weighted MR effects between the main exposures and epigenetic outcomes (GrimAge acceleration, PhenoAge acceleration and PAI1 levels), after adjustment for multiple testing (**Table 1; Table S24**). Of these, three remained significant (P <0.05) across the other three MR methods. All of these consistent effects were with GrimAge as the outcome. Greater adiposity was associated with greater GrimAge acceleration: BMI (Beta_IVW_ =0.76 years per SD increase in BMI, P =3.7×10^−16^); hip circumference (Beta_IVW_ =0.42 years per SD increase in hip circumference, P =2.5×10^−5^); waist circumference (Beta_IVW_ =0.59 years per SD increase in waist circumference, P =5.9×10^−6^; **Figure 4**). Current tobacco smoking showed evidence for a causal effect on increased GrimAge in two of the MR methods (Beta_IVW_ =3.42 years for smokers, P =9.0×10^−6^; **Figure 4**), as anticipated given that it incorporates a DNAm-based estimator of smoking pack-years [8]. Past tobacco smoking showed evidence for an inverse causal effect (Beta_IVW_ =-1.09 years, P =6.6×10^−9^), indicating that GrimAge acceleration is reduced upon smoking cessation. As a DNAm-proxy for leptin was also included in the derivation of GrimAge, the smoking and adiposity findings may act as positive controls. There was also evidence from three of the four MR methods to support a link between higher educational attainment (both years of schooling and college/university degree) and lower GrimAge acceleration (**Figure 4)**. For the secondary exposures, there was evidence across all methods for a causal effect of a greater body size at age 10 on higher GrimAge acceleration (Beta_IVW_ =0.70, P =1.6×10^−4^; **Table S23**; **Table S25**). Consistent findings across all four MR methods provided evidence to support a causal effect of 13 cell count traits and DNA methylation-estimated granulocyte proportions, and between lower lymphocyte proportions and higher Hannum and GrimAge acceleration (**Table S26**). There was evidence for heterogeneity in the causal effects for most of the cell types on epigenetic age measures, as well as years of schooling on GrimAge acceleration, although weaker evidence for directional pleiotropy was detected based on the Egger intercept (**Table S27**).

**Figure 4:**
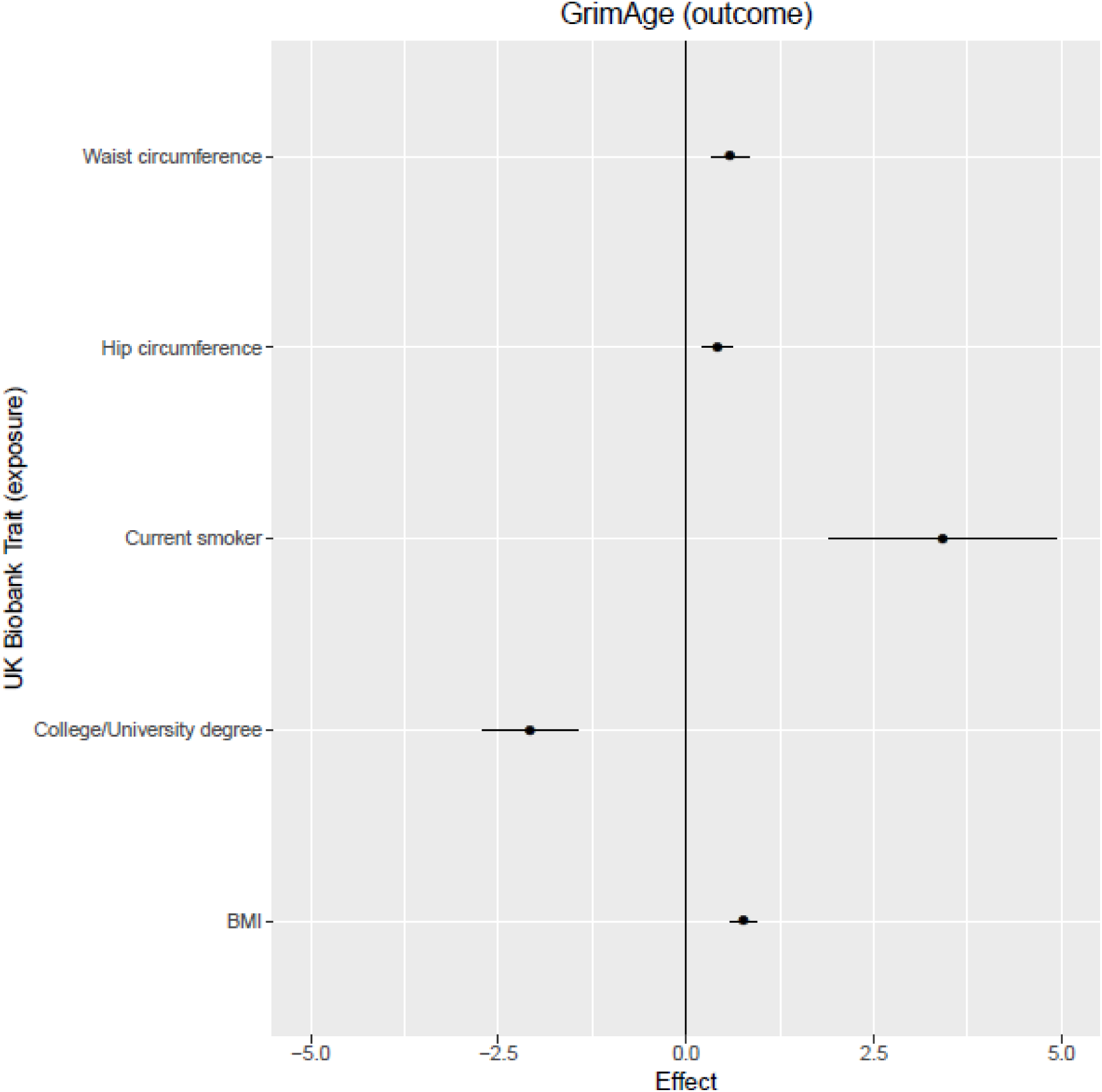
Causal effects of UK Biobank GWAS traits on GrimAge acceleration. Effects correspond to increase/decrease in GrimAge acceleration per SD increase in waist circumference, hip circumference and BMI; or per log odds increase for university/college education and current smoker status.

The biomarker analyses with epigenetic measures as outcomes identified no consistent effects across all four MR methods (**Table S26**). We found limited evidence to support any causal effects of the epigenetic measures (as exposures) on key disease and health outcomes, including longevity (**Tables S29-S30)**.

## Discussion

Epigenetic biomarkers of ageing and mortality have been extensively studied in relation to a plethora of health and disease outcomes. Here, we conducted a comprehensive suite of analyses in a meta-analysis sample of over 40,000 individuals, including the first GWA studies of two DNAm based estimators of mortality risk (PhenoAge and GrimAge), as well as for DNAm-based proxies for granulocyte proportion and plasminogen activation inhibitor 1 protein levels. We identified 137 loci, of which 113 were novel, related to the six epigenetic biomarkers. Although our comparison of genetic architectures across different ancestries was limited by sample size, our trans-ethnic meta-analyses implicated many shared genetic loci. However, heterogeneity of effect sizes between European and African American ancestries was found for genome wide significant loci in African Americans which may reflect differential tagging of underlying causal variants. Alternatively, gene-environment interactions or poor prediction of epigenetic biomarkers in African Americans may explain heterogeneity in these effect sizes.

GWAS results that are shared between IEAA (which adjusts for differences in white blood cell proportions) and the other epigenetic clocks (Hannum clock, PhenoAge, GrimAge) are less likely to be influenced by differences in cell composition IEAA and PhenoAge acceleration share the following genome-wide significant gene-based associations: *TPMT, TERT, NHLRC1, KDM1B, EDARADD*. IEAA and Hannum age acceleration share the following associations: *TERT, TRIM59, KPNA4, RP11-432B6*.*3, IFT80*, and *TET2. TET2* is particularly interesting in light of its mechanistic role (catalysing the conversion of methylcytosine to 5-hydroxymethylcytosine) and its established role in several ageing/regenerative phenotypes [51, 52].

Several of the GWAS overlapping genes in the European ancestry analysis (e.g. *TRIM59, KPNA4*) are also implicated by our eQTL colocalization analysis which identified regulatory relationships between SNPs associated with IEAA and expression of neighbouring genes. The IEAA-associated locus on chromosome 3 (lead SNP rs1047210) colocalized with an eQTL for *TRIM59*. DNA methylation levels at *TRIM59* have been robustly associated with chronological age and its expression has been noted in multiple cancers [53, 54, 55, 56, 57]. Apart from *TRIM59*, the eQTL colocalization also implicated other genes associated with cancer progression (e.g. *KPNA4, SMC4*, and *CXXC4*) [58, 59, 60]. *SMC4* is known to inhibit cellular senescence in replicating cells [61, 62]. Additional eQTL colocalization results for IEAA include *CD46*, a regulator of the complement system and T-cell function [63, 64], and the lipid transporter gene *ATP8B4*, which contains variants that have been reported in relation to centenarian status in Italians and Alzheimer’s disease [65, 66].

Using the findings from the European ancestry meta-analyses, we observed shared genetic contributions between PhenoAge and GrimAge acceleration with education, cognitive ability, adiposity, and smoking; there were also significant epigenetic biomarker-specific genetic correlations with numerous other health and modifiable lifestyle factors. The best epigenetic biomarker of mortality risk, GrimAge acceleration, exhibited strong genetic correlations with parental longevity and lung cancer. As GrimAge uses a DNA methylation based estimator of smoking pack-years in its definition, it is possible that the significant genetic correlation with GrimAge acceleration and lung cancer is mediated by smoking.

Four different Mendelian randomisation methods provide directional evidence of a causal influence of adiposity-related traits on GrimAge acceleration, while smoking cessation was inversely related to GrimAge acceleration. Several MR analyses indicate that increased educational attainment is associated with lower GrimAge acceleration. However, there was no causal evidence for associations with other lifestyle traits, such as alcohol consumption. There was limited evidence from MR to implicate any of the DNAm predictors as playing an important causative role in longevity or disease risk.

This is the first study to present polygenic risk scores for six epigenetic biomarkers of ageing. A phenome-wide scan of the six polygenic risk scores (PRS) yielded highly significant associations between the PRS of GrimAge acceleration and adiposity-related traits, education, and parental longevity. Both the PRS analysis and the MR analyses suffer from two limitations i) the geographical structure in the UK Biobank cohort might confound these analyses [67] and ii) low heritability estimates for some of the phenotypes (e.g. longevity or epigenetic biomarkers such as PAI1). Limited statistical power due to low heritability or low sample size may help explain the disconnect between the genetic correlation analysis (which revealed a plethora of significant genetic correlations for GrimAge and PhenoAge) and the MR analysis (which led to a dearth of significant findings). In general, careful interpretation of the MR findings is required. In particular, causal MR analyses that modelled epigenetic biomarkers as exposures and disease states as outcomes suffered from weak genetic instruments (e.g., for GrimAge acceleration and granulocyte proportions, where the variance explained was <1%) or inadequate power in two-sample analysis. Future multivariate MR analyses will be required to test whether the protective causal effect of education on GrimAge is mediated by smoking, obesity, or other factors.

Overall, this study highlights the shared genetic architecture between epigenetic ageing, lifestyle factors (smoking, obesity), and parental longevity, which shows that DNAm-based biomarkers are valuable endophenotypes of biological ageing.

## Supporting information

Appendix 1: Cohort Descriptions

Appendix 2: Heterogeneity Analyses

## Acknowledgements

REM, SH, and AL are supported by a National Institute of Health U01 grant, U01AG060908–01. REM and DLM are supported by Alzheimer’s Research UK major project grant, ARUK-PG2017B-10. RCR is a de Pass Vice Chancellor’s Research Fellow at the University of Bristol. CR and RCR receive support from a Cancer Research UK Programme Grant (C18281/A191169). CLR, JLM, and RCR are members of the UK Medical Research Council Integrative Epidemiology Unit at the University of Bristol (MC_UU_00011/5). IJD is supported by Age UK (Disconnected Mind programme), UKRI Medical Research Council grant, MR/R0245065/1, and by National Institute of Health R01 grant, 1R01AG054628-01A1. GD is supported by the University of Edinburgh School of Philosophy, Psychology and Language Sciences. Molecular data for the Trans-Omics in Precision Medicine (TOPMed) program was supported by the National Heart, Lung and Blood Institute (NHLBI). Core support including centralized genomic read mapping and genotype calling, along with variant quality metrics and filtering were provided by the TOPMed Informatics Research Center (3R01HL-117626-02S1; contract HHSN268201800002I). Core support including phenotype harmonization, data management, sample-identity QC, and general program coordination were provided by the TOPMed Data Coordinating Center (R01HL-120393; U01HL-120393; contract HHSN268201800001I). We gratefully acknowledge the studies and participants who provided biological samples and data for TOPMed. Cohort specific acknowledgements are presented in **Appendix 1**.

## Methods

### Study cohort information

The meta-analysis sample comprised 37 datasets from 31 cohorts encompassing 41,731 participants (controls/healthy volunteers). Of these, 28 included individuals of European ancestry comprising 34,449 participants, seven were of individuals of African American comprising 6,148 participants, and one was of Hispanic ancestry comprising 287 participants. The total meta-analysis sample age range was 27.2-79.1 years (mean 54.0 years overall; 54.8 years across European ancestry cohorts; 50.4 years across African American cohorts), and comprised 0-100% females (mean 58.26%; 57.3% across European ancestry cohorts; 58.3% across African American cohorts). Cohort-level descriptive data are presented in **Table S1** and described in **Appendix 1**. Each of the studies was approved by their local Ethical Committee. All subjects provided written informed consent.

### Data availability

Meta-analysis summary statistics for each epigenetic biomarker are publicly available at https://datashare.is.ed.ac.uk/handle/10283/3645. For cohort-specific details, please see **Appendix 1**, which contains information for each study.

### Data preparation

Age-adjusted DNA methylation-based estimates of Hannum age, Intrinsic Horvath age, PhenoAge, GrimAge, plasminogen activator inhibitor-1 levels, and unadjusted granulocyte proportion were calculated using the Horvath epigenetic age calculator software (https://dnamage.genetics.ucla.edu/ or standalone scripts provided by Steve Horvath and Ake Lu). The following outputs were assessed: Intrinsic Epigenetic Age Acceleration – “IEAA”, Hannum age acceleration – “AgeAccelerationResidualHannum”, PhenoAge acceleration – “AgeAccelPheno”, GrimAge acceleration – “AgeAccelGrim”, estimate levels of Plasminogen Activation Inhibitor 1, adjusted for age – “DNAmPAIadjAge”, and estimated proportion of granulocytes – “Gran”. For each cohort, an outlier threshold for methylation values of +/- 5 standard deviations was applied and outlier samples were excluded from the analysis.

### GWAS and meta-analysis

Quality control and imputation were done by each study separately (**Appendix 1**) Genotypes were imputed to either Haplotype Reference Consortium (HRC) or 1000 genomes phase 3 panels [68, 69]. In each cohort, association testing was conducted using imputed dosages using an additive model. Linear models were adjusted for sex and genetic principal components. GWAS summary statistics were obtained for between 1,097,816-15,221,271 genetic markers. This was the case for all cohorts with the exception of GOLDN (whole-genome sequence data) and the Sister Study (imputed data not available at the time of analysis). For each cohort, summary statistics were processed and harmonised using the R package EasyQC [70]. Multi-allelic variants were filtered to contain the variant with the highest minor allele count. At the individual cohort level, variants that were monomorphic, with a minor allele count ≤ 25, genotyped in <30 individuals, or with an imputation quality score <0.6 were removed. Allele codes and marker names were harmonized, duplicate variants were removed, and allele frequency checks were performed against the appropriate population reference data. Meta-analyses were performed with METAL using an inverse variance fixed effects scheme [71]. Meta-analyses were performed on European ancestry and African American studies separately (n=34,962 and 6,482, respectively). Variants were omitted from the meta-analysis if they were absent from >50% of the total meta-analysis sample size. Cohort specific genomic inflation factors ranged from 0.86 to 1.07. Genome-wide significance was defined as P < 5×10^−8^. To summarize the associations in terms of index SNP with the strongest association and other SNPs in linkage disequilibrium we used conditional and joint association analysis of GWAS summary data, including the HLA region, in the GCTA-COJO software. A stepwise selection model was used with default settings for SNP LD (R^2^<0.9), analysis window size (10Mb), and genome-wide significance (P<5×10^−8^) using HRC imputed genotype data from Generation Scotland, and 1000G imputed genotype data from ARIC as the reference panels for the European ancestry and African American analyses, respectively. Heterogeneity I^2^ statistics were obtained from the meta-analyses and plotted against both –log_10_ p-values and effect sizes to determine if SNPs with heterogeneous effects across cohorts were more statistically significant or had larger effect sizes. Systematic between-study heterogeneity was also investigated [21]. Meta-analyses were re-run after excluding cohorts identified as outliers and effect sizes were visually compared with the full meta-analysis output. Forest plots were prepared for all significant loci.

### Trans-ethnic meta-analysis

A trans-ethnic meta-analysis of all European ancestry and African American cohorts was conducted using default settings in the Meta-Regression of Multi-Ethnic Genetic Association (MR-MEGA) tool [25]. We considered summary output for the first principal component of the meta-regression.

MR-MEGA summary statistics were uploaded to the Functional Annotation of Meta-Analysis Summary Statistics (FUMA) (http://fuma.ctglab.nl) software for annotation of the top loci using default settings, selecting 1000 Genomes phase 3 (all populations) as the reference population [72]. Independent lead SNPs had P<5×10^−8^ and were independent of each other at r^2^<0.6; lead SNPs within this subset were required to have r^2^<0.1. A locus was defined by considering lead SNPs in a 250kb range and all SNPs in LD (r^2^≥0.6) with at least one independent SNP.

### Functional Annotation of Meta-Analysis Summary Statistics

The European ancestry and African American meta-analysis summary statistics were uploaded to FUMA (http://fuma.ctglab.nl) for further annotation and functional analysis [72]. Genes were annotated from SNP-level data using the “SNP2GENE” tool, permitting gene set and tissue expression analyses using MAGMA [26]. A Bonferroni-corrected significance threshold (adjusting for 18,606 tested genes) of P <2.69×10^−6^ was set for the gene-based GWAS. Genes annotated to significant GWAS loci were further investigated using the “GENE2FUNC” tool in FUMA for enrichment of GWAS catalog gene sets [73]. Bonferroni-corrected P-value thresholds were applied.

### Functional enrichment

To test if the GWAS meta-analysis findings were associated with regulatory and functional features of interest, enrichment analyses were conducted using GARFIELD [30]. SNPs were first pruned (r^2^>0.1) then annotated to categories (e.g., chromatin states, histone modifications, DNaseI hypersensitive sites, and transcription factor binding sites). Statistical enrichment was then carried out for SNPs at two p-value thresholds (P<1×10^−5^ and P<1×10^−8^) while accounting for MAF, distance to the nearest TSS, and number of LD proxies.

### Colocalization Analysis

We hypothesised that some of the lead loci from the meta-analyses will have shared variants 1) across the Epigenetic biomarkers, 2) with DNAm sites in blood, and 3) with gene expression levels in blood. We used GoDMC summary statistics on 190,102 DNAm sites [74] to examine the overlap between loci and epigenetic clock DNAm sites, BMI-associated DNAm sites and smoking-associated DNAm sites. We used eQTL Gen summary statistics on 19,942 transcripts [75] that were available in the MR-Base database [76]. We used the Rpackage gwasglue (https://mrcieu.github.io/gwasglue/) to extract SNPs that were +/- 1Mb of the lead SNP and to harmonise the datasets. For each Epigenetic biomarkers molecular trait pair (or pair of Epigenetic biomarkers) we then performed colocalization analysis using the coloc.abf function in the R/coloc package [24], using default parameters. We only kept colocalized pairs with more than 50 shared SNPs and a posterior probability above 0.8. We removed the *HLA* region from the eQTL colocalization analysis.

### Disease and Phenotype Ontology Enrichment

The potential role of Mendelian disease genes and associated pathways in influencing the epigenetic clock was investigated with MendelVar [31], independently for each marker phenotype. We did not limit our analysis to any particular phenotype class amongst the Mendelian disease genes but looked for enrichment of any disease processes found to be strongly linked to genes in the GWAS loci. MendelVar analysis was run using intervals based on ±0.5 Mbp window around the lead SNPs using the 1000 Genomes EUR population as LD reference [66]. Inside MendelVar, INRICH was run in “target” enrichment mode, with the target gene set filter set at minimum 5 (-i option) and maximum of 20000 (-j option), and minimum observed threshold of 2 (-z option) [77]. The nominal p-values were corrected for multiple testing with two rounds of permutation in INRICH.

### Heritability and Genetic Correlation Analysis

LD score regression, using LD scores and weights estimated from European ancestry populations (downloaded from https://data.broadinstitute.org/alkesgroup/LDSCORE/), was used to assess genetic correlations between the six epigenetic biomarkers. Genetic correlations were further assessed between the six epigenetic biomarkers and publicly-available GWAS summary statistics using the LDHub web interface (http://ldsc.broadinstitute.org/ldhub/) [41]. Meta-analysis results for each epigenetic biomarker were uploaded to the LDHub website, selecting all available traits for genetic correlation analysis. SNP heritability was estimated using univariate LD score regression. As the majority of large-scale GWA studies have been based on European ancestry populations, heritability and genetic correlation analyses were limited to this group to maximise statistical power. Filtering was performed to exclude traits where the LD Hub output came with the following warning messages: “Caution: using these data may yield less robust results due to minor departure of the LD structure” and “Caution: using this data may yield results outside bounds due to relative low Z score of the SNP heritability of the trait”. This left a total of 693 unique traits from 708-711 studies per epigenetic clock. A Bonferroni-corrected significance threshold of P < 0.05/693 = 7.21×10^−5^ was applied.

### Polygenic risk scores

To determine the proportion of variance in the six epigenetic biomarkers that can be explained by common genetic variants, we carried out a polygenic risk score analysis using results from the European ancestry meta-analyses. Weights for the additive genetic scores were created by re-running the meta-analyses excluding one cohort (test cohort) at a time. Six weighted-PGR-scores (one for each epigenetic biomarker) were generated using default settings of the PRSice software (clump-kb=250, clump-p=1; clump-r2=0.25) [78]. P-value thresholds were set at <5×10^−8^, <0.01, <0.05, <0.1, <0.5, and 1. Linear regression models were built to calculate the incremental R^2^ between the null model (epigenetic biomarker∼ sex) and full model (epigenetic biomarker ∼ sex + polygenic risk score) in the test cohort. The procedure was iterated after excluding different test cohorts one by one (Lothian Birth Cohort 1921, Lothian Birth Cohort 1936, Framingham Heart Study, Born in Bradford, ARIES, and SABRE, respectively) from the meta-analysis. Finally, these steps were repeated, using the full meta-analysis summary statistic output to generate polygenic risk scores in a completely independent cohort (Young Finns Study, n =1,320).

For Born in Bradford, ARIES and SABRE, best-guess genotypes files with a MAF cut-off of 1% and info score>0.8 were generated and the polygenic risk score analyses were corrected for 20 genetic PCs.

A phenome-wide association study of 581 heritable traits from the UK Biobank study was then carried out for parsimonious polygenic risk scores based on independent SNPs with P<5×10^−8^ or P<1 from each of the six GWAS meta-analyses (http://mrcieu.mrsoftware.org/PRS_atlas/) [49]. The P-value thresholds were based on the leave-one-out cohort PRS analyses described above (GrimAge acceleration and Granulocyte proportions: P<1; IEAA, Hannum Age acceleration, PhenoAge acceleration, IEAA and PAI1 levels: P<5×10^−8^). An FDR-corrected P-value (P_FDR_ <0.05) was applied separately to each set of PheWAS results.

### Mendelian Randomisation

To investigate if the epigenetic biomarkers were i) causally influenced by lifestyle factors and ii) had a causal effect on ageing and disease outcomes, Mendelian Randomisation (MR) was performed in MRBase [76]. The epigenetic measures were considered as both exposures (i.e., causally influencing the outcome) and outcomes (i.e., the epigenetic measure being causally influenced by a trait of interest). The analyses were further split into four sections: primary exposures/outcomes (common lifestyle risk factors and ageing/disease outcomes from the largest available GWAS in MR Base); secondary exposures/outcomes (traits identified as relevant via moderate genetic correlations from the LD regression analyses); 34 cell count exposures [20]; and 38 biomarker exposures [79]. SNPs instrumenting each exposure were clumped using a European LD reference panel and an r2 < 0.001. Harmonization of the SNP effects with the exposure and outcomes were performed so that palindromic SNPs were aligned when minor allele frequencies (MAFs) were <0.3 or were otherwise excluded.

Inverse-variance weighted (IVW) MR was carried out as the main analysis, with pleiotropy-robust sensitivity analyses featuring MR-Egger [80], weighted median [81], and weighted mode MR [82]. Significant associations were defined by a Bonferroni-corrected P-value < 0.05. Where there was evidence for a causal effect based of the IVW model, we also assessed the potential for horizontal pleiotropy by means of heterogeneity assessment (Cochran’s Q-statistic) of individual SNP effects in both IVW and MR-Egger analyses and the Egger intercept test for directional pleiotropy [80].

## Supplementary Tables

**Table S1:** Cohort summaries.

**Table S2:** Cohort-level genomic inflation.

**Table S3:** Independent SNP-level GWAS results.

**Table S4:** GWAS summary statistics for SNPs associated with IEAA and EEAA (Gibson et al.) and Granulocyte counts (Astle et al.)

**Table S5:** Colocalization of independent SNPs across all epigenetic biomarkers

**Table S6:** Independent loci identified in the trans-ancestry meta-analysis of six epigenetic biomarkers using FUMA.

**Table S7:** Summary statistics of ancestry-specific variants. Effects frequencies, p-values and standard errors for lead SNPs identified exclusively in European ancestry analysis (upper table) and African American analysis (lower table) are compared against African American summary statistics and European ancestry summary statistics, respectively.

**Table S8:** Summary statistics for lead SNPs identified in the trans-ancestry meta-analysis of six epigenetic biomarkers, queried against a Hispanic ancestry population (MESA)

**Table S9:** Gene-level GWAS results.

**Table S10:** mQTL colocalization summary for six epigenetic biomarkers, BMI and smoking.

**Table S11:** mQTL summary statistics for six epigenetic biomarkers

**Table S12:** Functional and regulatory enrichment results using GARFIELD.

**Table S13:** Pathway enrichment.

**Table S14:** Enrichment among Mendelian disease genes and associated ontology/pathway terms.

**Table S15:** SNP-level overlap with associations from the GWAS catalog.

**Table S16:** Gene enrichment among GWAS catalog traits.

**Table S17:** eQTL colocalization analysis.

**Table S18:** Heritability analysis.

**Table S19:** Genetic correlations between six epigenetic epigenetic biomarkers.

**Table S20:** Genetic correlations between six epigenetic epigenetic biomarkers and publicly-available GWAS traits.

**Table S21:** Polygenic risk score analysis.

**Table S22:** PRSAtlas output

**Table S23:** Study-specific information for primary and secondary MR traits

**Table S24:** Mendelian randomisation analysis of primary trait exposures

**Table S25:** Mendelian randomisation analysis of secondary trait exposures

**Table S26:** Mendelian randomisation analysis of cell count-related exposures

**Table S27:** Pleiotropy assessment for exposures with evidence of a causal effect on epigenetic measures.

**Table S28:** Mendelian randomisation analysis of biomarker-related exposures

**Table S29:** Mendelian randomisation analysis of clock exposures (primary traits)

**Table S30:** Mendelian randomisation analysis of clock exposures (secondary traits)

## Supplementary Figures

**Figure S1:** QQ Plot for granulocyte proportions in the European ancestry GWAS meta-analysis.

**Figure S2:** QQ Plot for GrimAge Acceleration in the European ancestry GWAS meta-analysis.

**Figure S3:** QQ Plot for Hannum Age Acceleration in the European ancestry GWAS meta-analysis.

**Figure S4:** QQ Plot for IEAA in the European ancestry GWAS meta-analysis

**Figure S5:** QQ Plot for PAI1 in the European ancestry GWAS meta-analysis.

**Figure S6:** QQ Plot for PhenoAge Acceleration proportions in the European ancestry GWAS meta-analysis.

**Figure S7:** QQ Plot for granulocyte proportions in the African American GWAS meta-analysis.

**Figure S8:** QQ Plot for GrimAge Acceleration in the African American GWAS meta-analysis.

**Figure S9:** QQ Plot for Hannum Age Acceleration in the African American GWAS meta-analysis.

**Figure S10:** QQ Plot for IEAA in the African American GWAS meta-analysis.

**Figure S11:** QQ Plot for PAI1 in the African American GWAS meta-analysis.

**Figure S12:** QQ Plot for PhenoAge Acceleration proportions in the African American GWAS meta-analysis.

**Figure S13:** Mean/SD epigenetic age plotted against mean/SD chronological age in European ancestry and African American cohorts.

**Figure S14:** Manhattan Plot for granulocyte proportions in the European ancestry GWAS meta-analysis.

**Figure S15:** Manhattan Plot for GrimAge Acceleration in the European ancestry GWAS meta-analysis.

**Figure S16:** Manhattan Plot for Hannum Age Acceleration in the European ancestry GWAS meta-analysis.

**Figure S17:** Manhattan Plot for IEAA in the European ancestry GWAS meta-analysis.

**Figure S18:** Manhattan Plot for PAI1 in the European ancestry GWAS meta-analysis.

**Figure S19:** Manhattan Plot for PhenoAge Acceleration proportions in the European ancestry GWAS meta-analysis.

**Figure S20:** Plot of effect sizes for genome-wide significant SNPs in Gibson et al. vs effect sizes in a lookup of the current meta-analysis results for Hannum Age Acceleration and IEAA.

**Figure S21:** Plot of effect sizes for genome-wide significant SNPs in Astle et al. vs effect sizes in a lookup of the current meta-analysis results for granulocyte proportion (and vice versa).

**Figure S22:** Manhattan Plot for granulocyte proportions in the African American GWAS meta-analysis.

**Figure S23:** Manhattan Plot for GrimAge Acceleration in the African American GWAS meta-analysis.

**Figure S24:** Manhattan Plot for Hannum Age Acceleration in the African American GWAS meta-analysis.

**Figure S25:** Manhattan Plot for IEAA in the African American GWAS meta-analysis.

**Figure S26:** Manhattan Plot for PAI1 in the African American GWAS meta-analysis.

**Figure S27:** Manhattan Plot for PhenoAge Acceleration proportions in the African American GWAS meta-analysis.

**Figure S28:** Plot of lead African- and European ancestry trans-ethnic meta-analysis SNP effect sizes against the same SNPs in the Hispanic subset of the MESA cohort.

**Figure S29:** Lookup of 24 blood-based independent genome-wide significant SNPs for IEAA in a brain-based GWAS of IEAA (overlap of 21 SNPs). The red line represents the linear regression line (r=0.74) for SNPs that are also mQTLs for IEAA clock CpG sites. The turquoise line represents the linear regression line (r=0.08) for SNPs that are not mQTLs for IEAA clock CpGs. Labelled points correspond to loci where there was strong evidence of SNPs sharing genetic effects with eQTLs.

**Figure S30:** LocusZoom plot for the region (*DSRC6*/*RIPPLY3*) with highest evidence of genetic colocalization for the blood- and brain-based GWASs. Note that the lead SNP from the blood-based GWAS also colocalizes with a mQTL for an IEAA clock CpG, cg13450409 (PP=0.99, **Table S11**).

**Figure S31:** LD regression SNP-based heritability estimates for the six epigenetic biomarkers.

## Notes

### Competing Interest Statement

The authors have declared no competing interest.

