## Supplementary Figures for "Genome-wide association studies identify 137 loci for DNA methylation biomarkers of ageing"

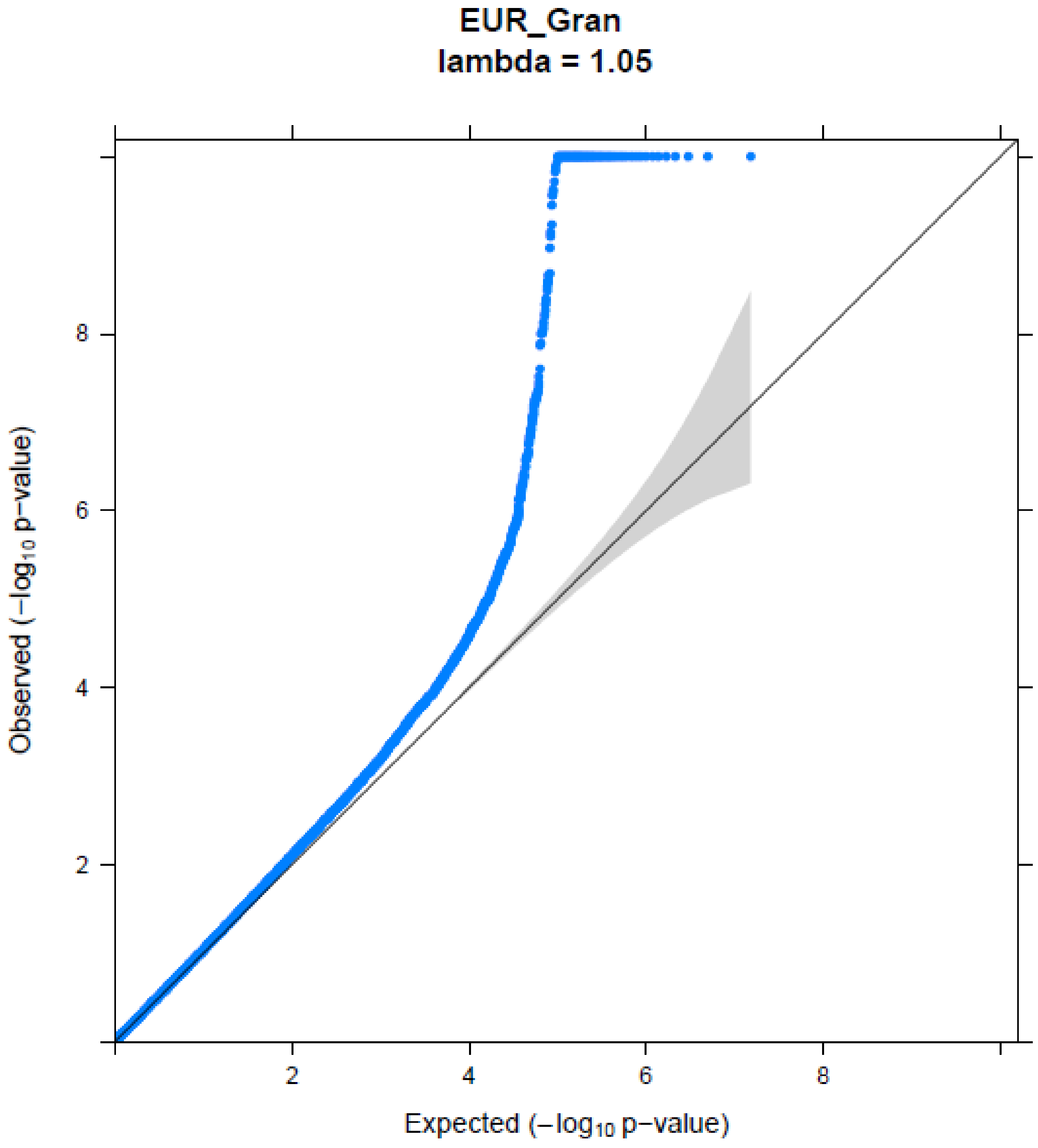


**Figure S1:** QQ Plot for granulocyte proportions in the European ancestry GWAS meta-analysis.

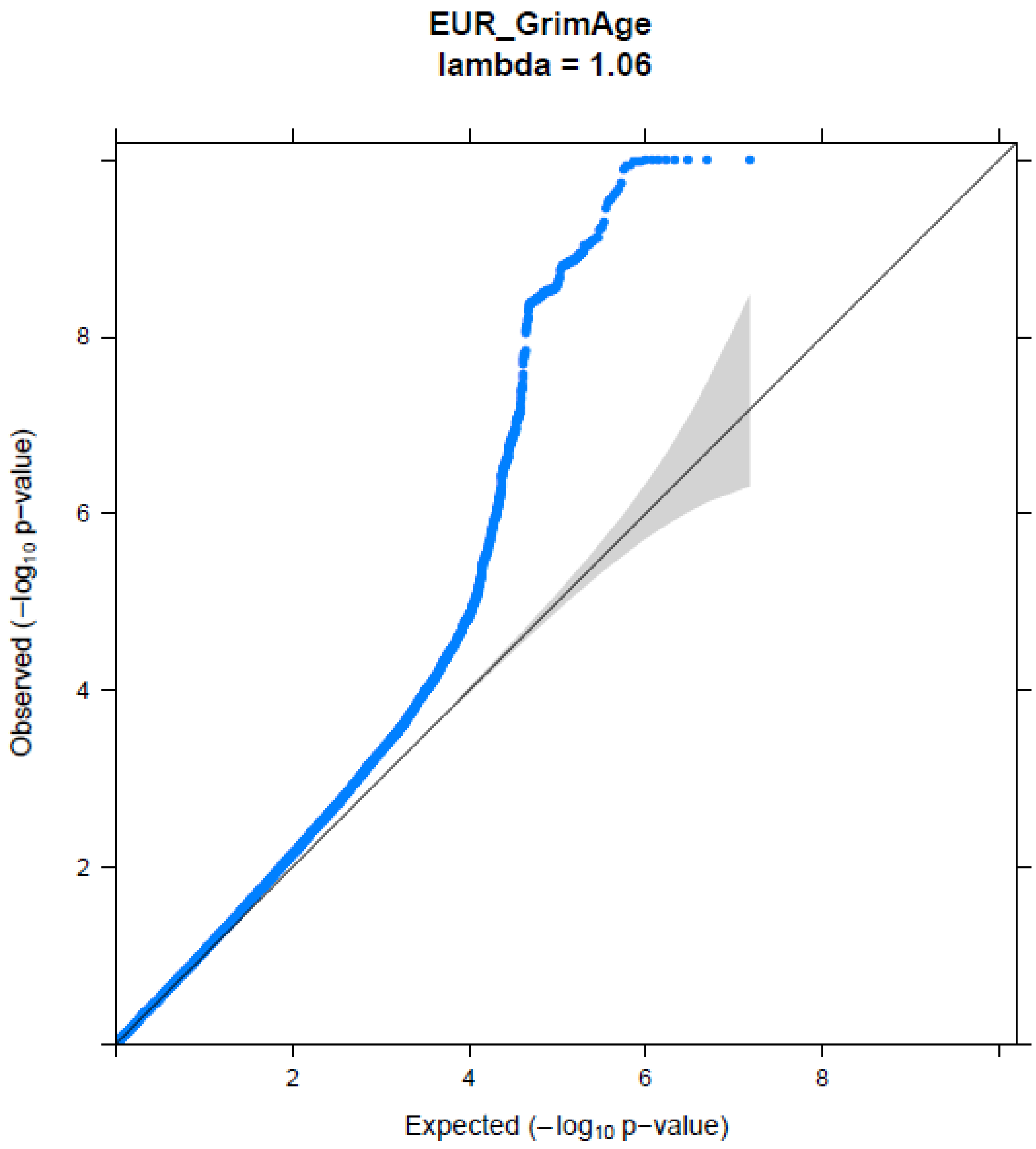


**Figure S2:** QQ Plot for GrimAge Acceleration in the European ancestry GWAS meta-analysis.

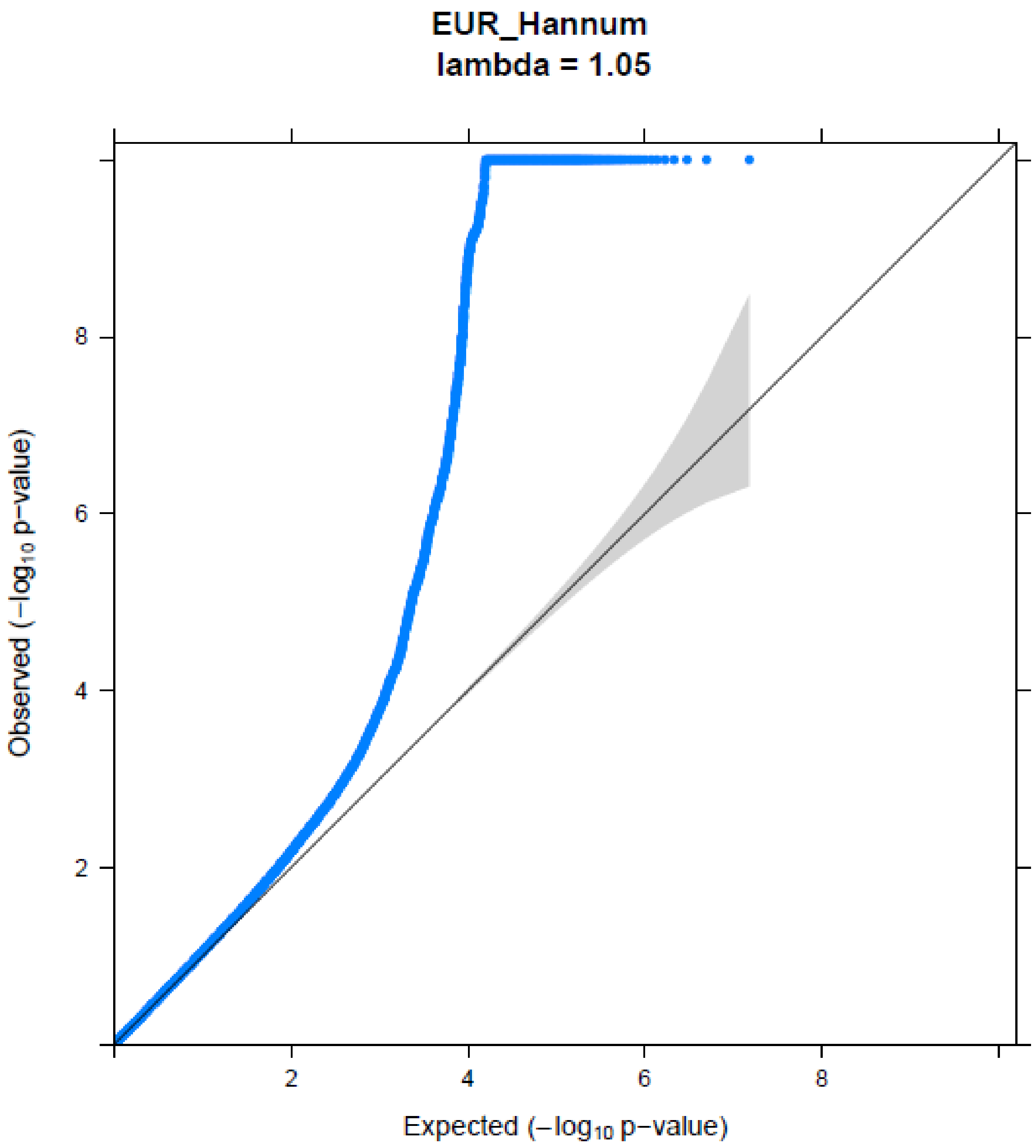


**Figure S3:** QQ Plot for Hannum Age Acceleration in the European ancestry GWAS meta-analysis.

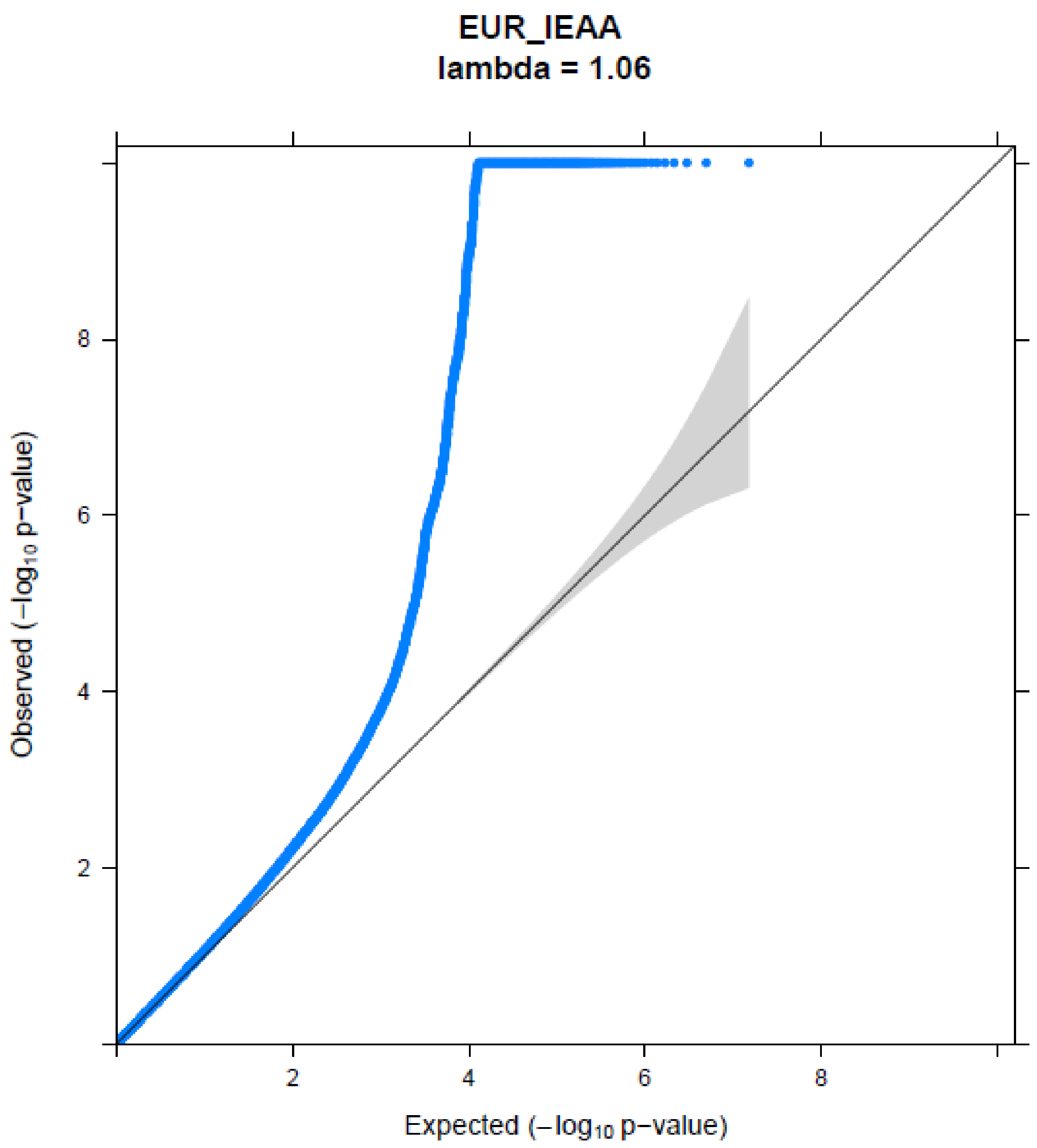


**Figure S4:** QQ Plot for IEAA in the European ancestry GWAS meta-analysis

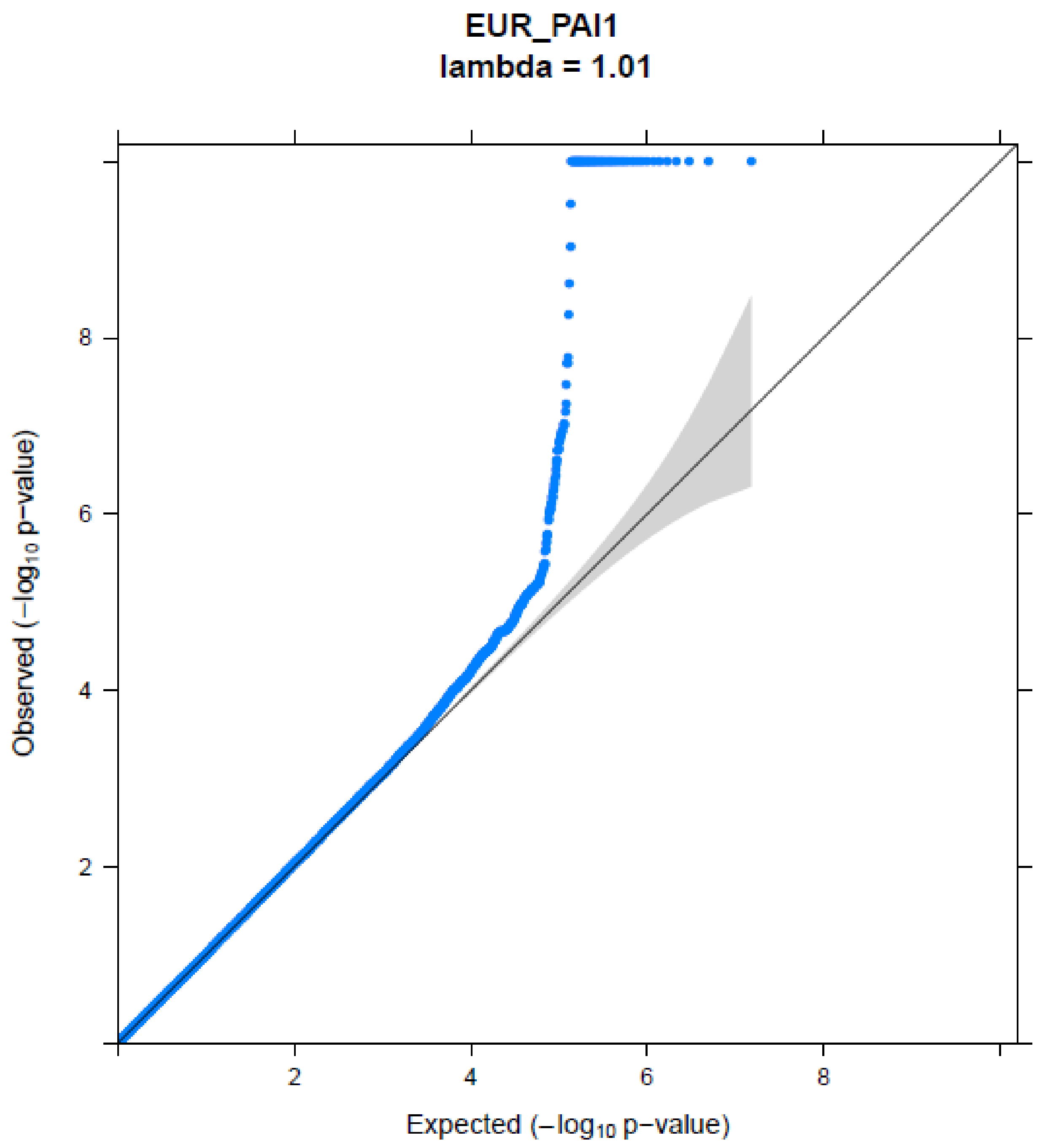


**Figure S5:** QQ Plot for PAI1 in the European ancestry GWAS meta-analysis.

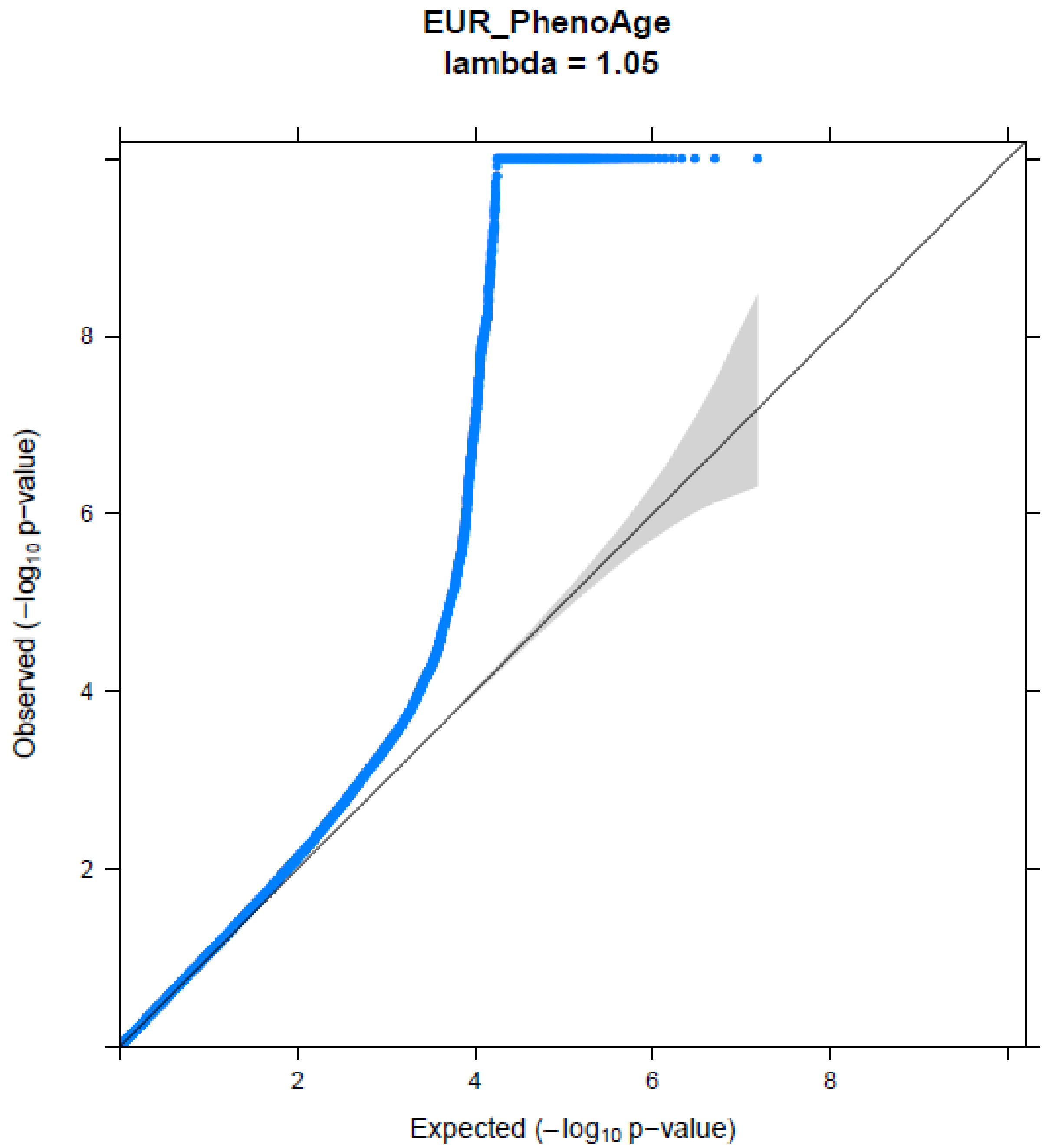


**Figure S6:** QQ Plot for PhenoAge Acceleration proportions in the European ancestry GWAS meta-analysis.

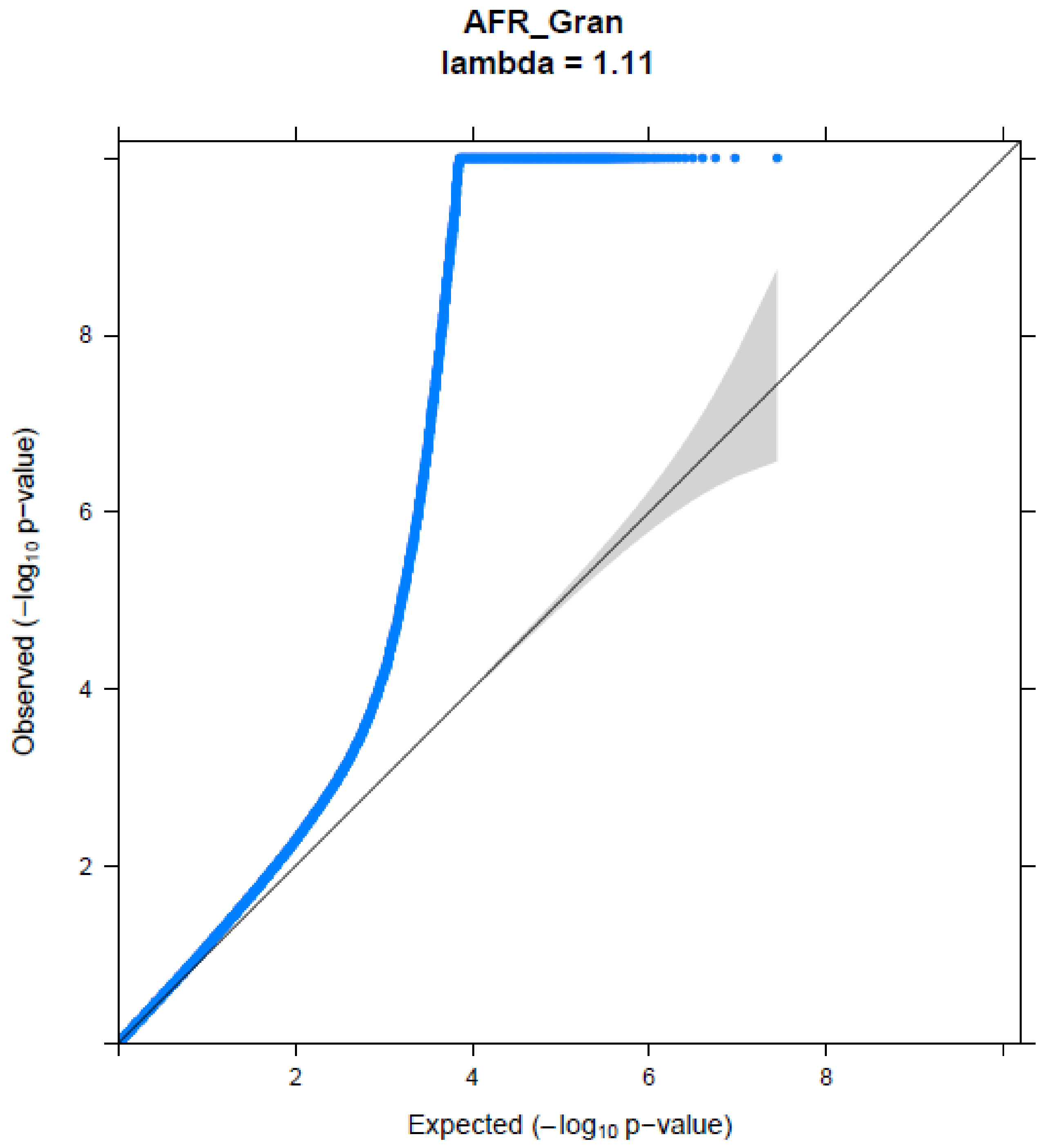


**Figure S7:** QQ Plot for granulocyte proportions in the African American GWAS meta-analysis.

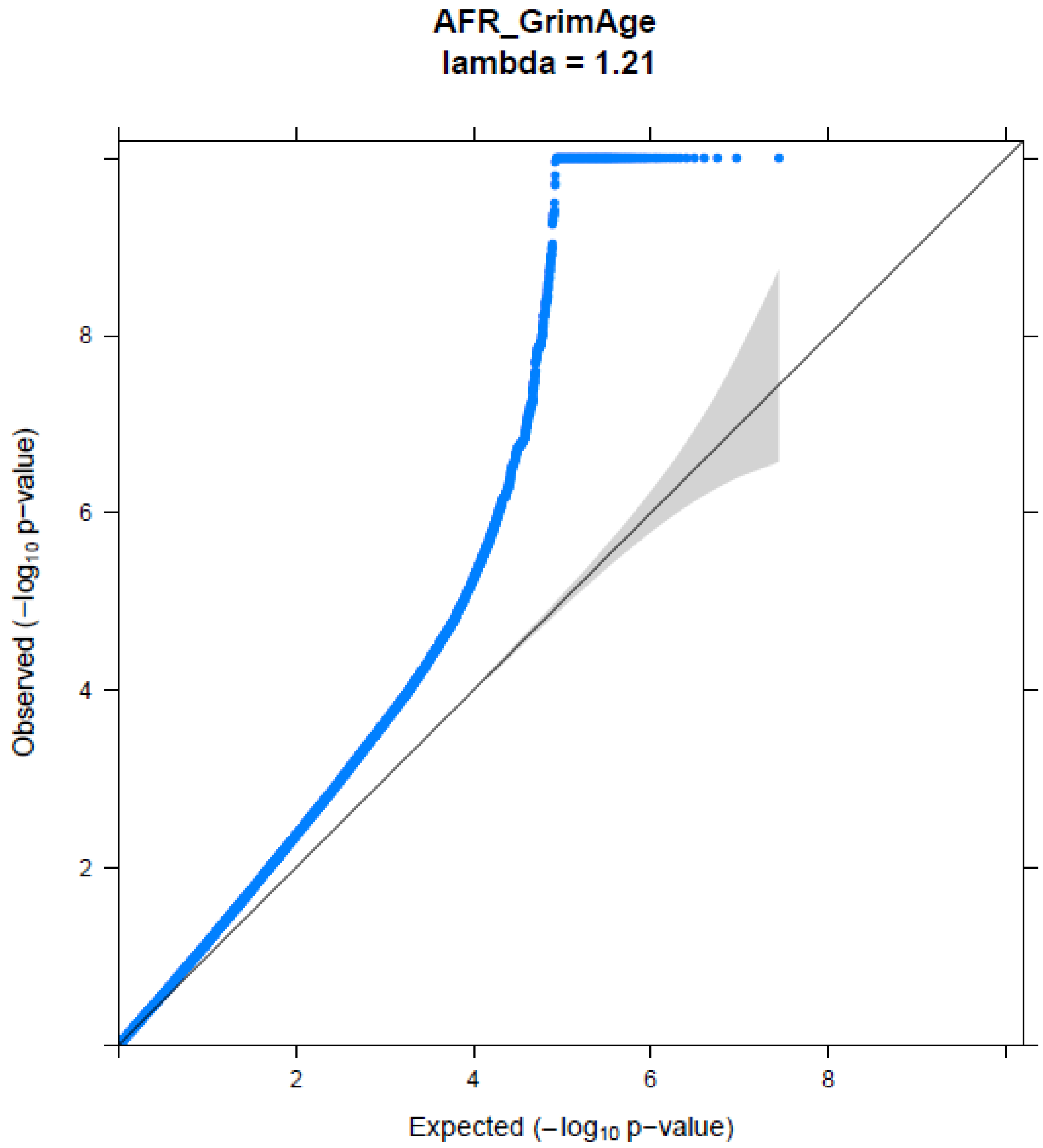


**Figure S8:** QQ Plot for GrimAge Acceleration in the African American GWAS meta-analysis.

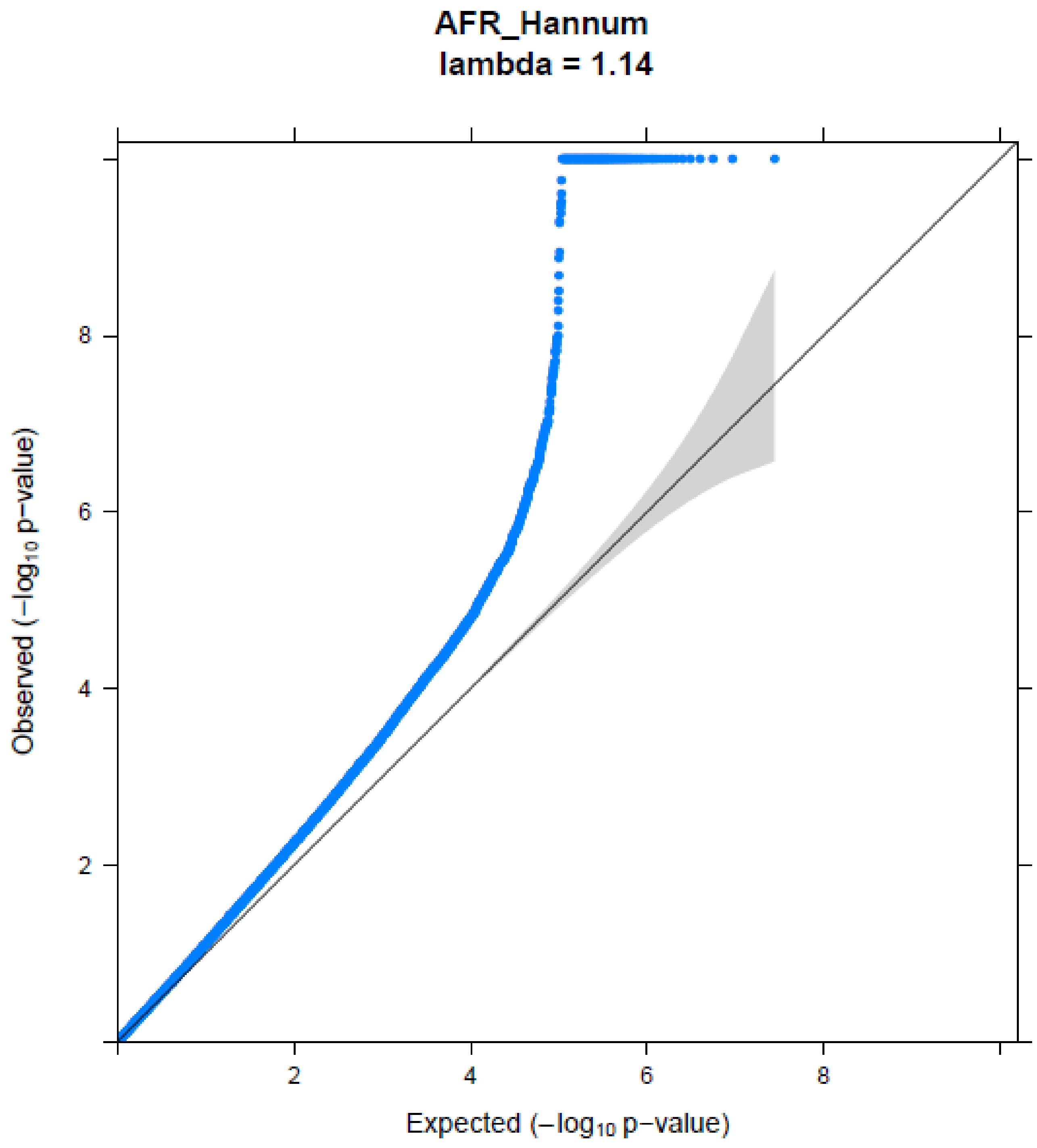


**Figure S9:** QQ Plot for Hannum Age Acceleration in the African American GWAS meta-analysis.

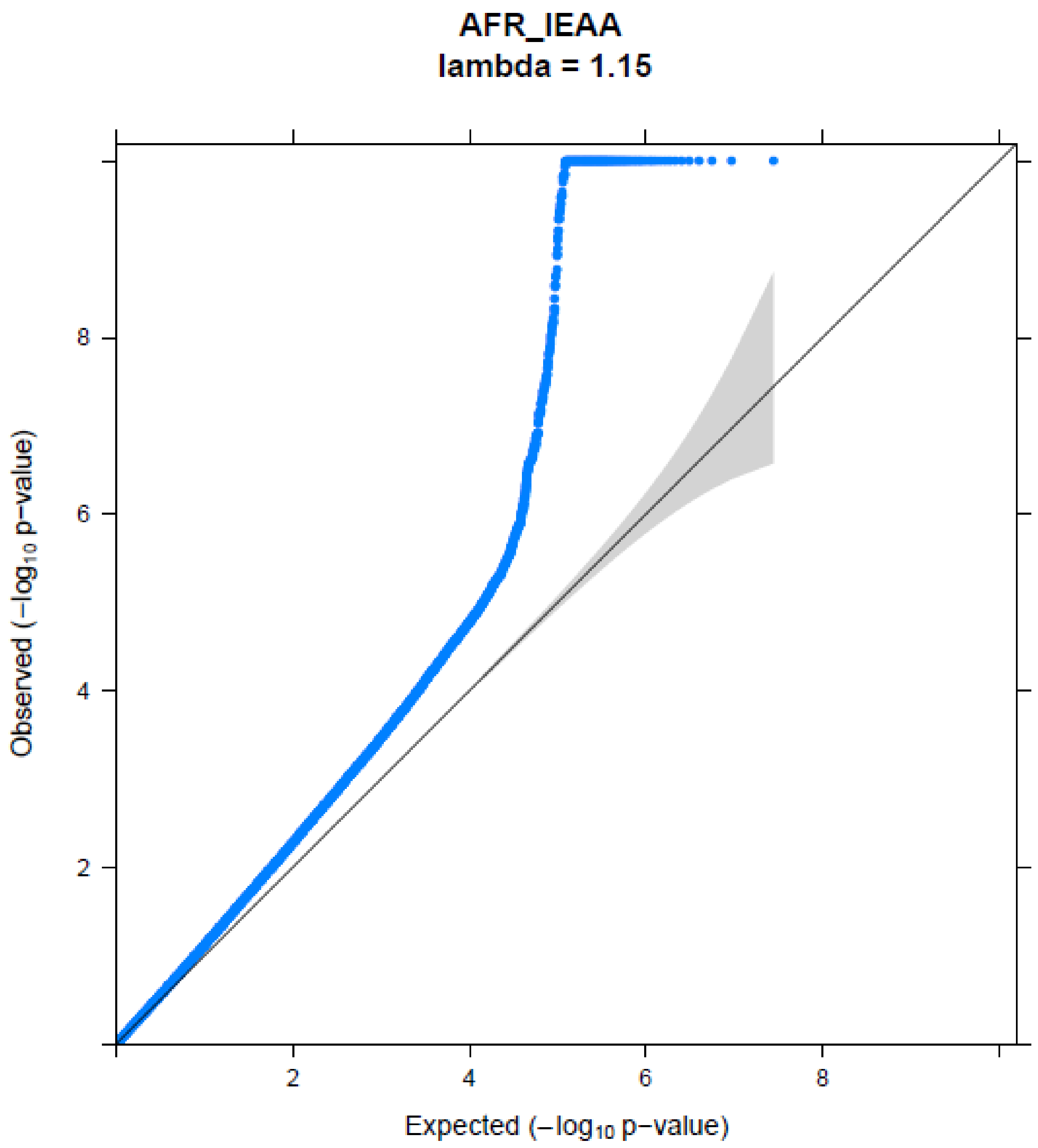


**Figure S10:** QQ Plot for IEAA in the African American GWAS meta-analysis.

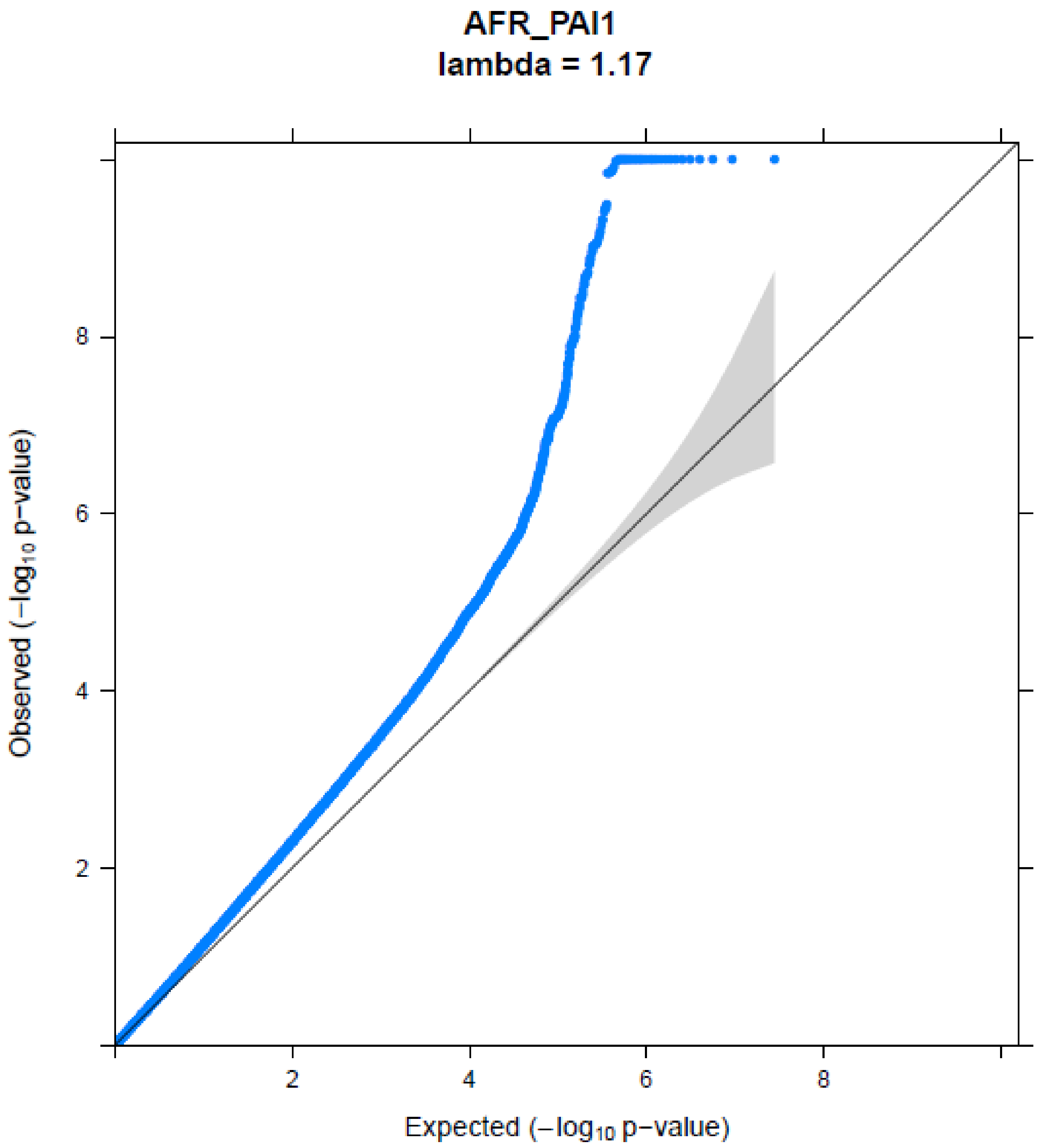


**Figure S11:** QQ Plot for PAI1 in the African American GWAS meta-analysis.

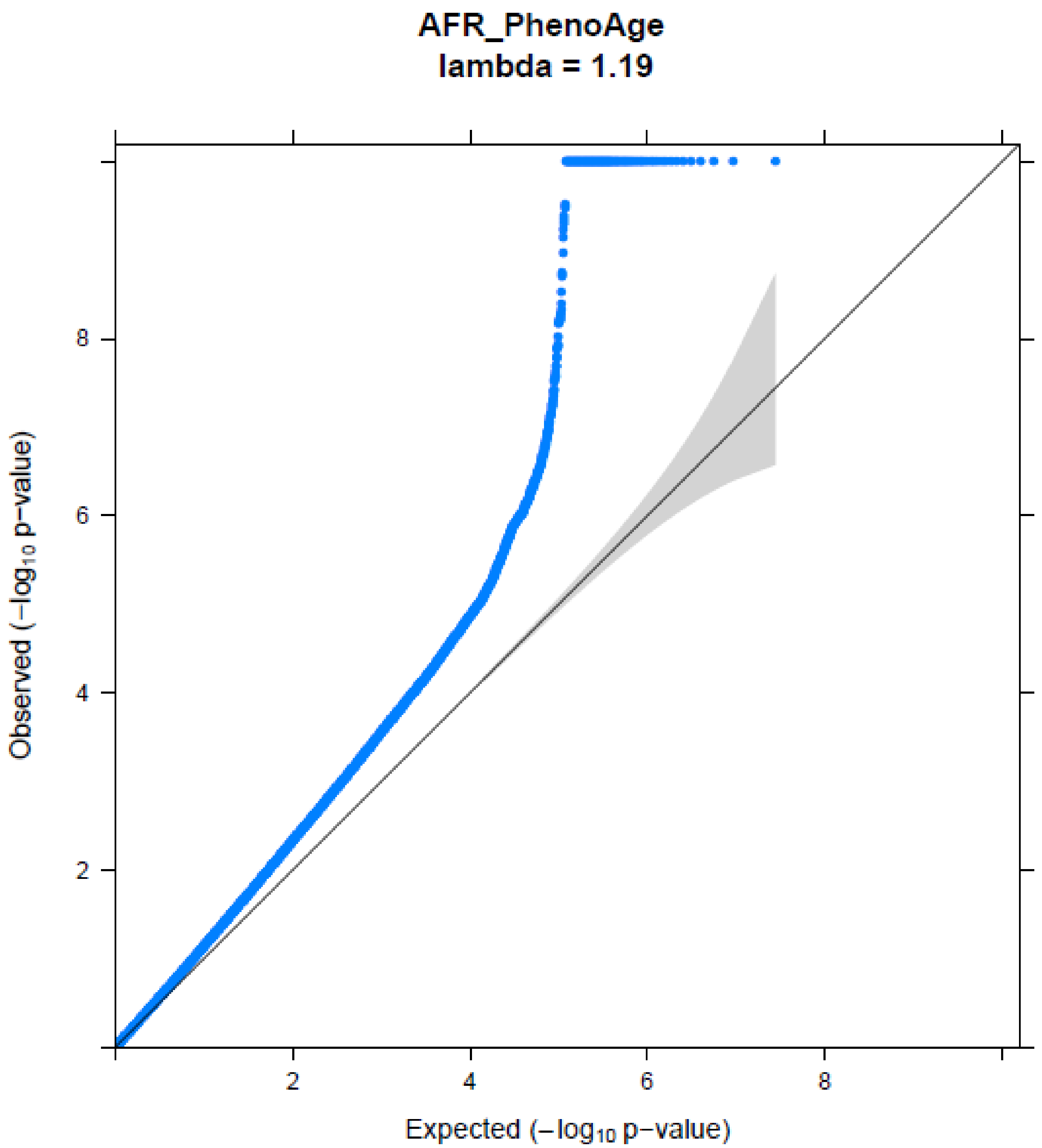


**Figure S12:** QQ Plot for PhenoAge Acceleration proportions in the African American GWAS meta-analysis.

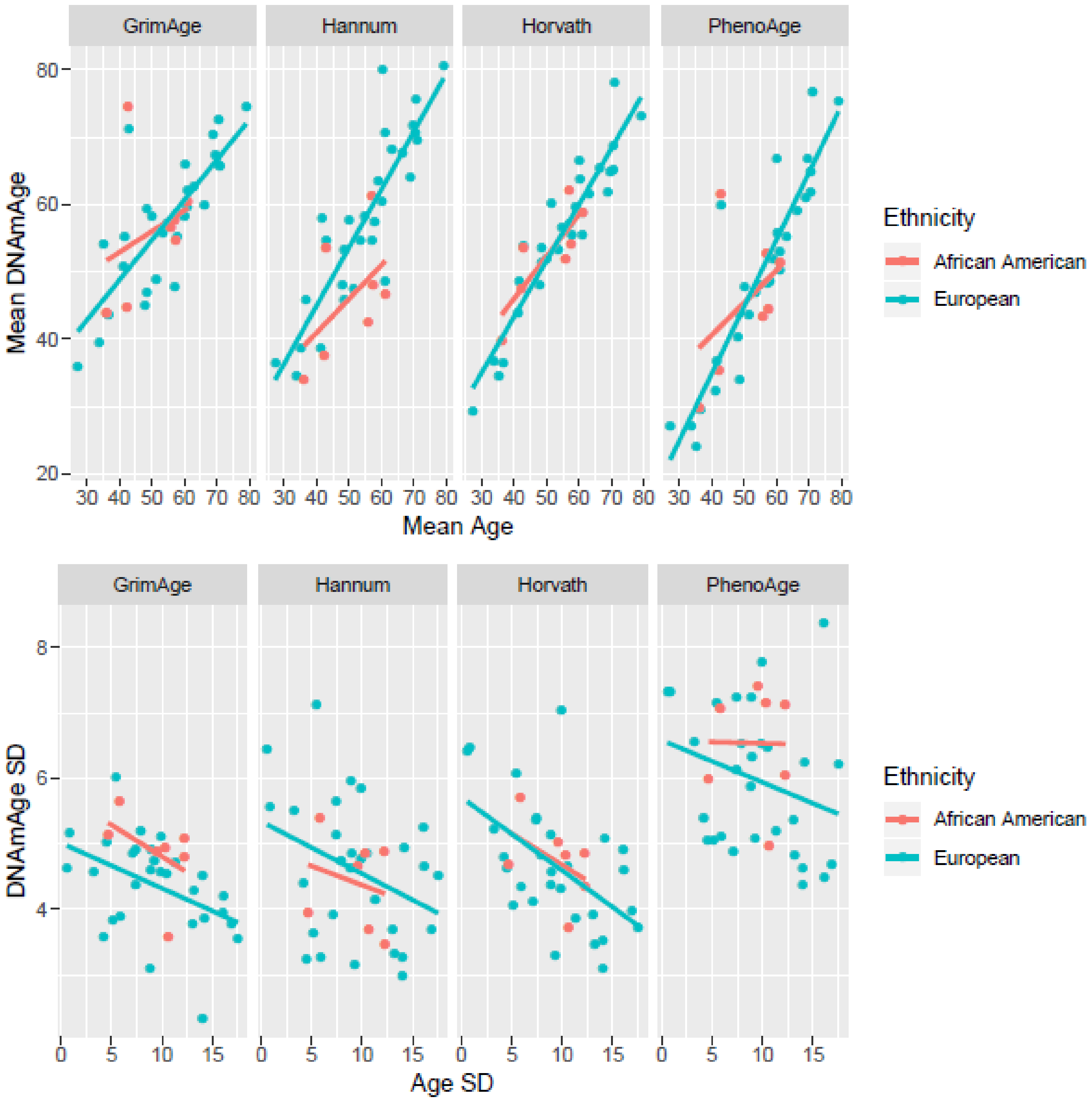


**Figure S13:** Mean/SD epigenetic age plotted against mean/SD chronological age in European ancestry and African American cohorts.

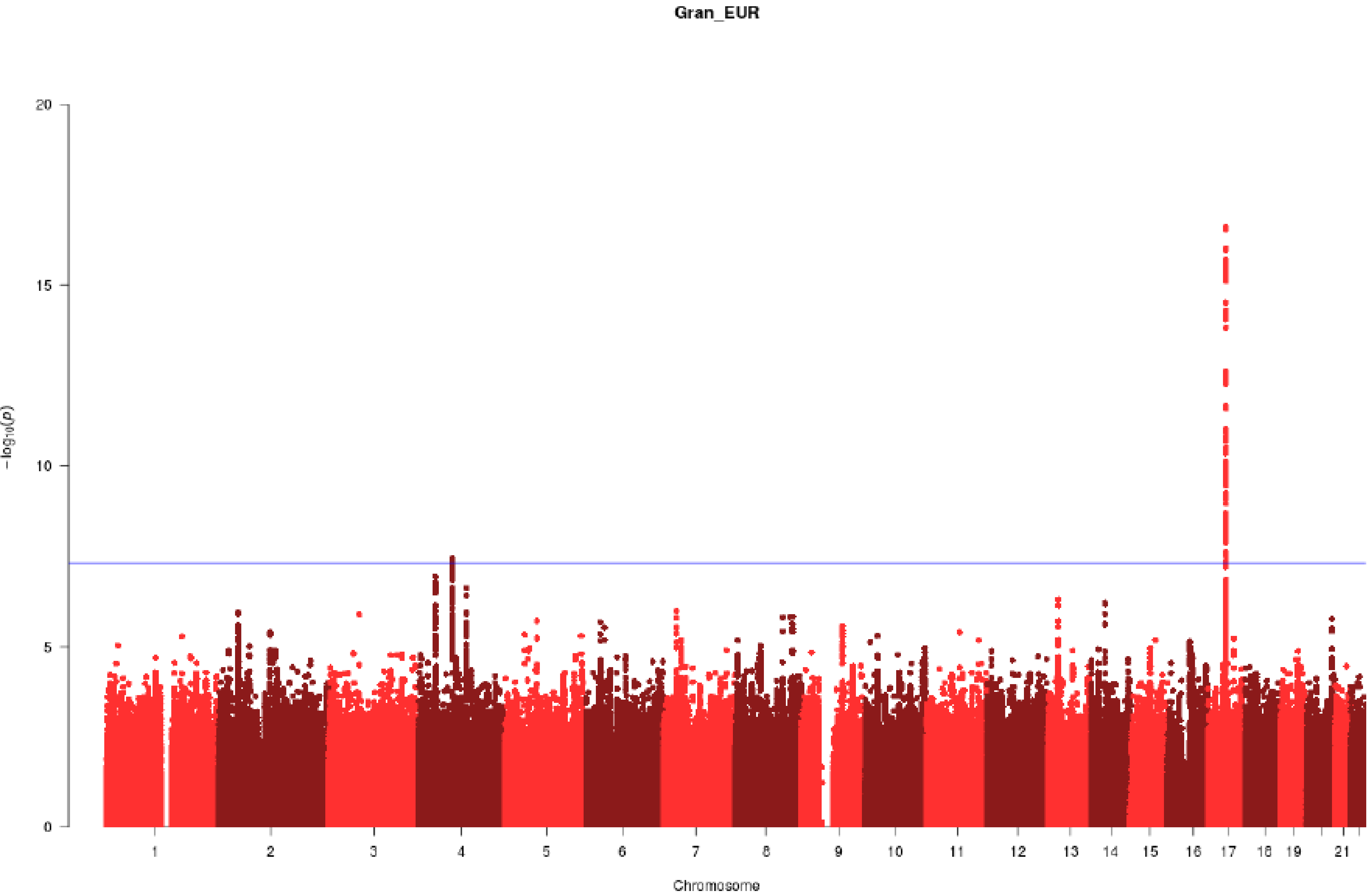


**Figure S14:** Manhattan Plot for granulocyte proportions in the European ancestry GWAS meta-analysis.

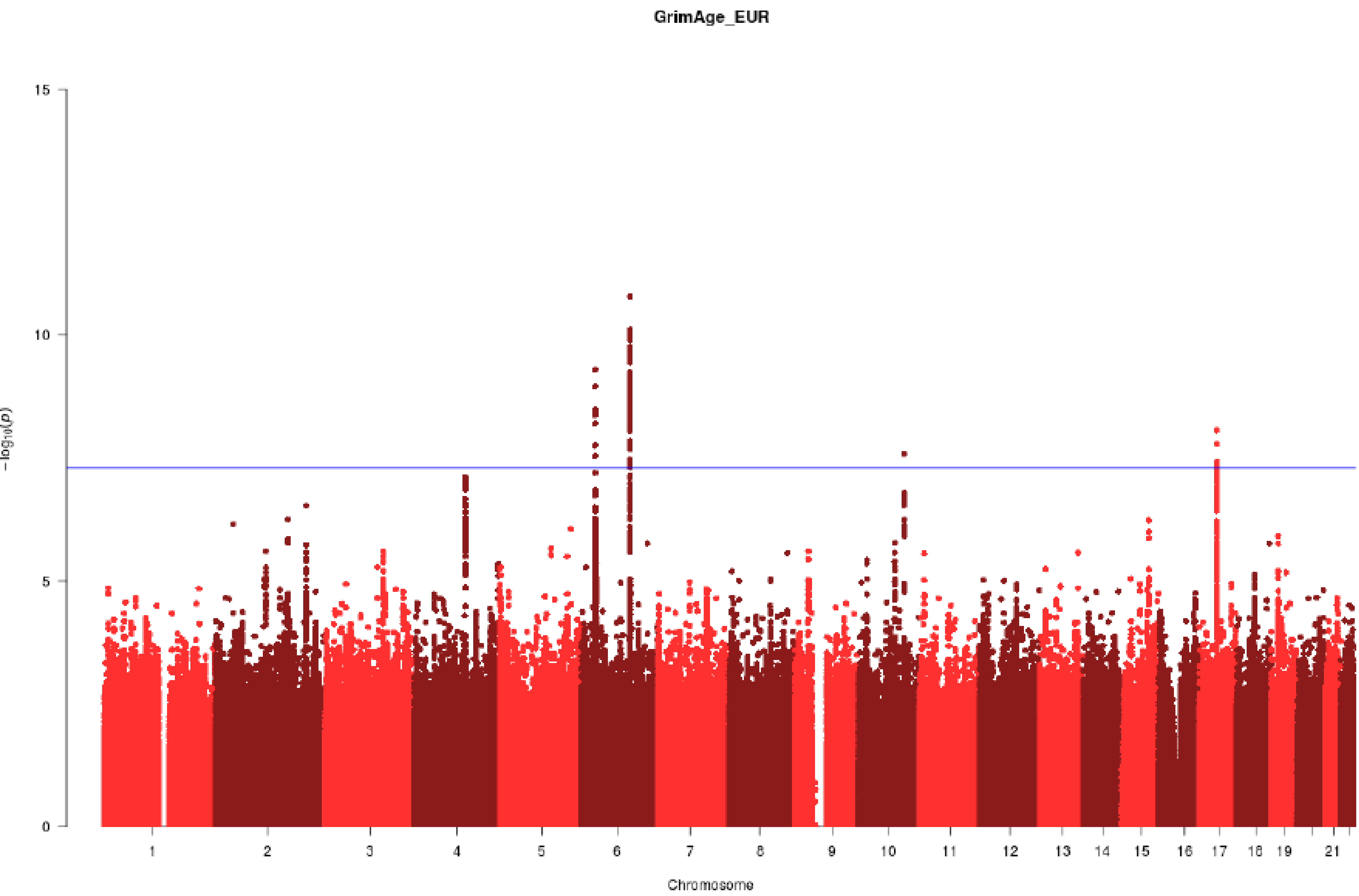


**Figure S15:** Manhattan Plot for GrimAge Acceleration in the European ancestry GWAS meta-analysis.

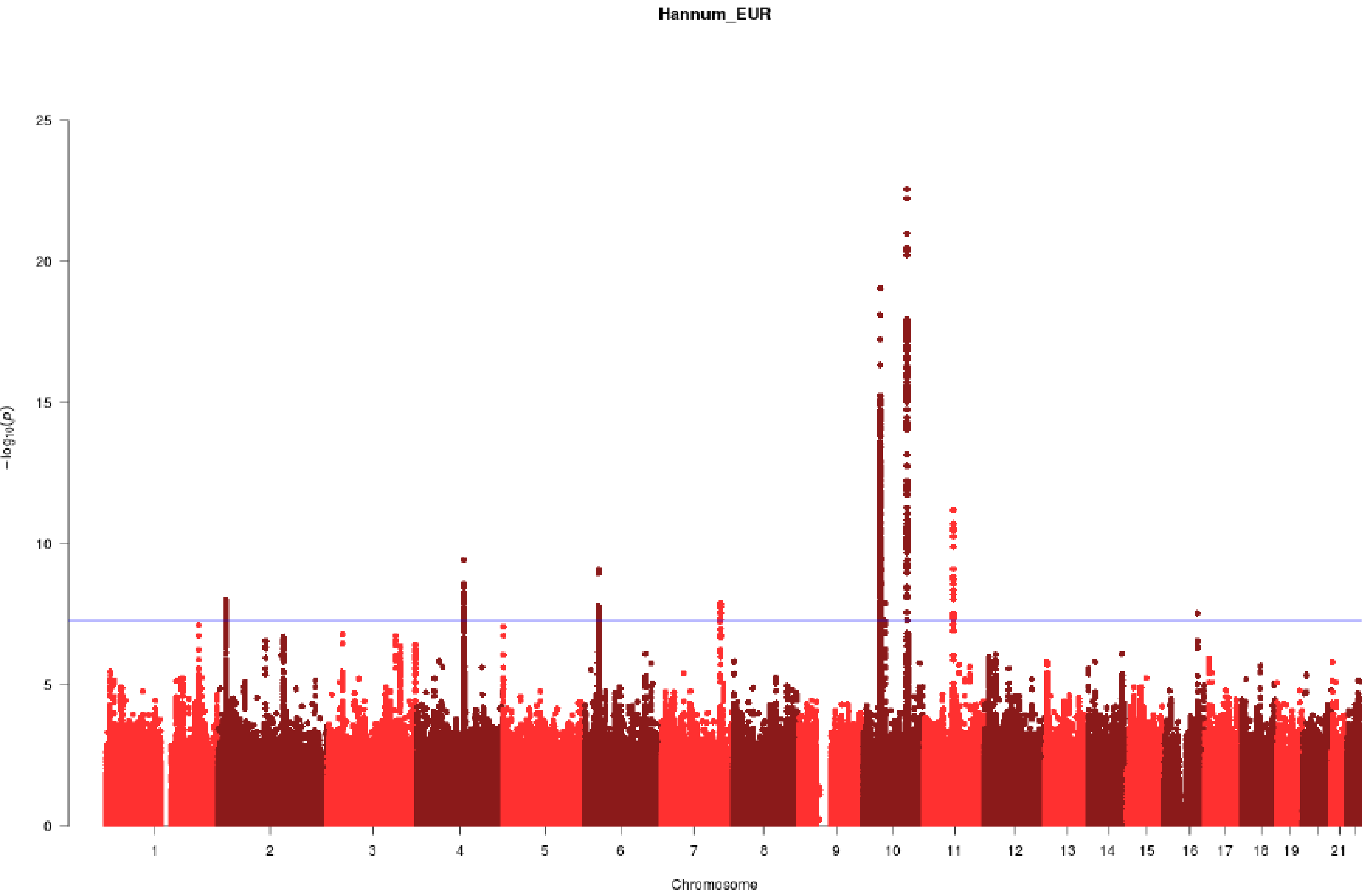


**Figure S16:** Manhattan Plot for Hannum Age Acceleration in the European ancestry GWAS meta-analysis.

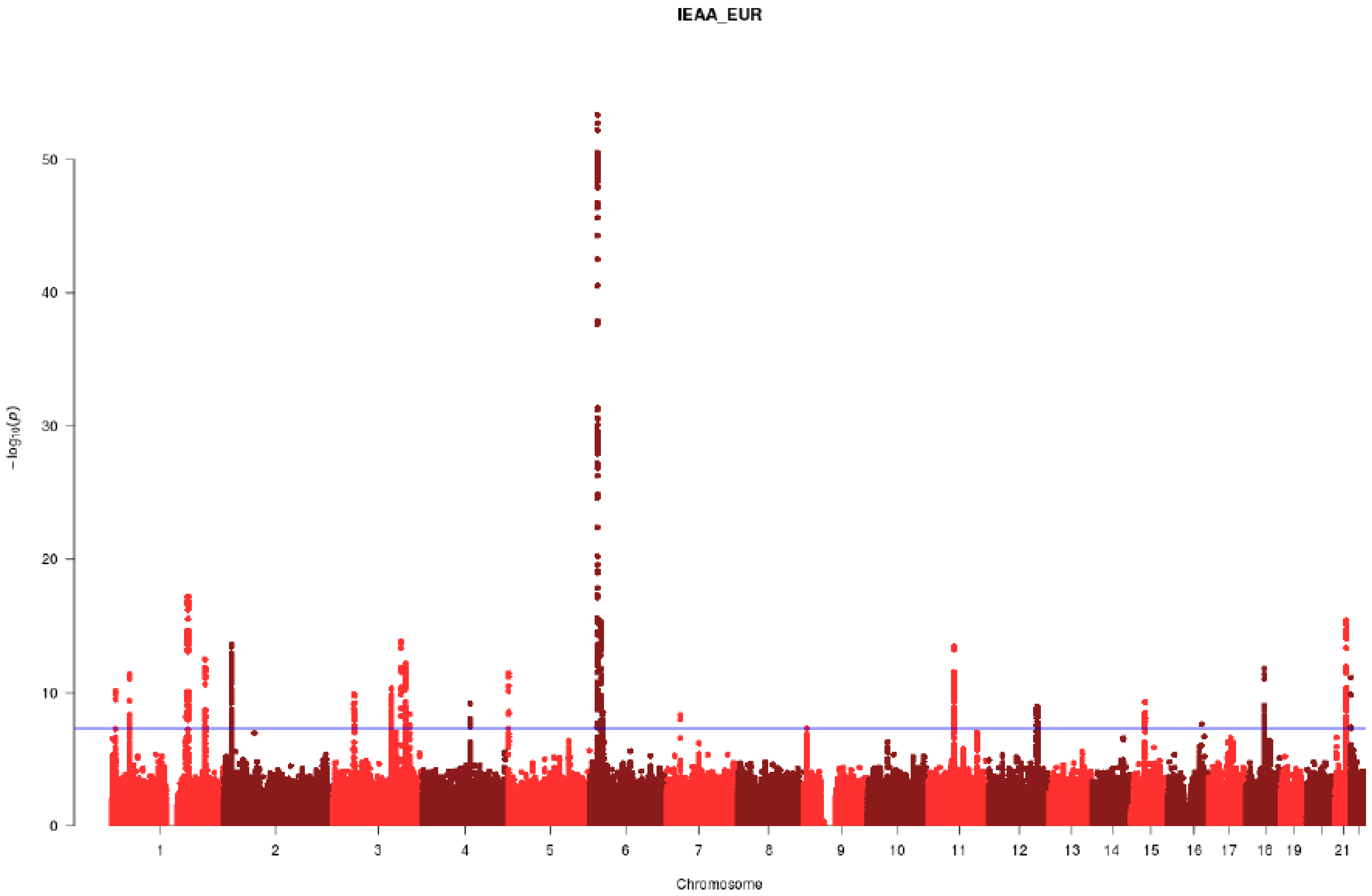


**Figure S17:** Manhattan Plot for IEAA in the European ancestry GWAS meta-analysis.

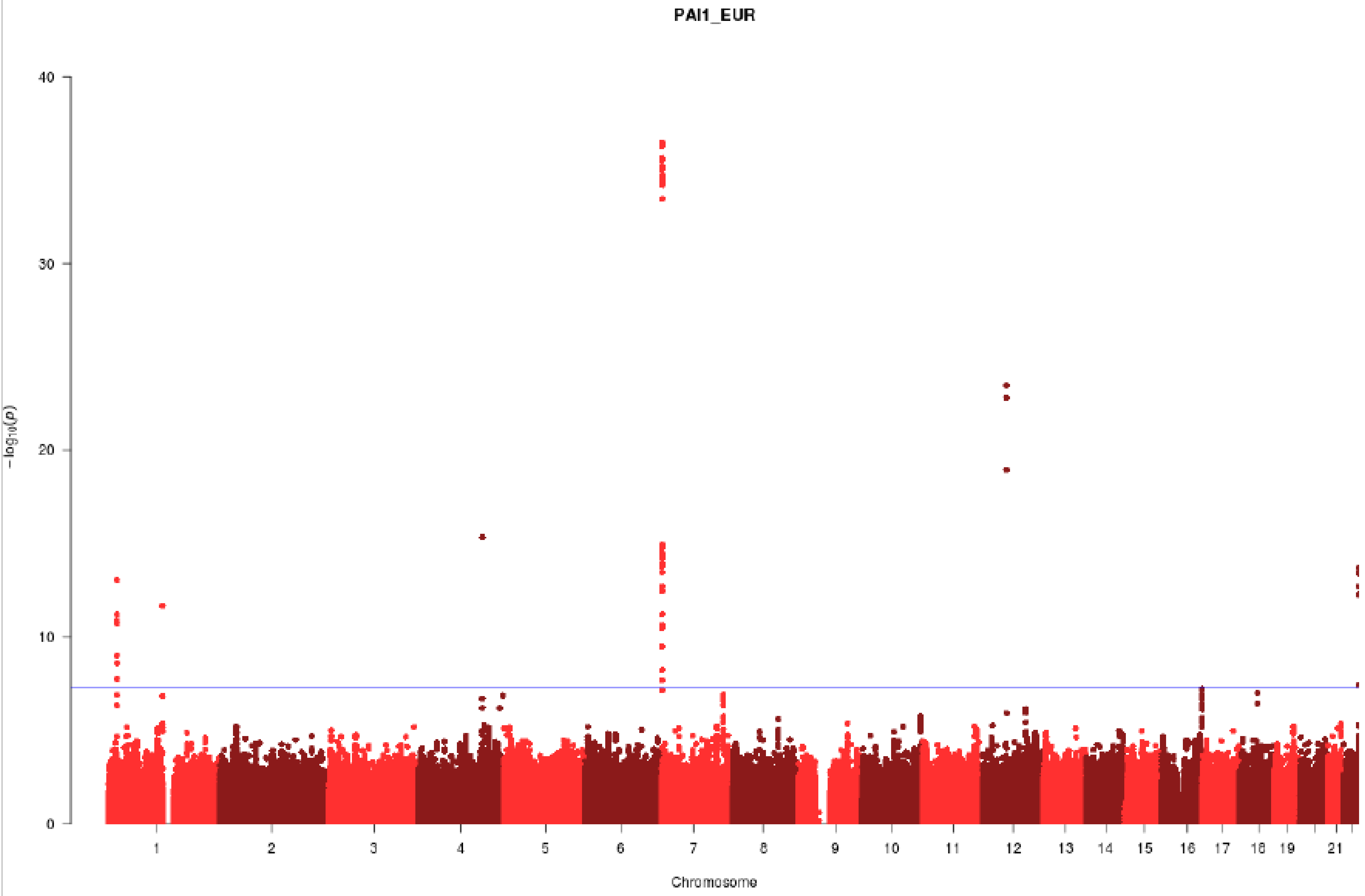


**Figure S18:** Manhattan Plot for PAI1 in the European ancestry GWAS meta-analysis.

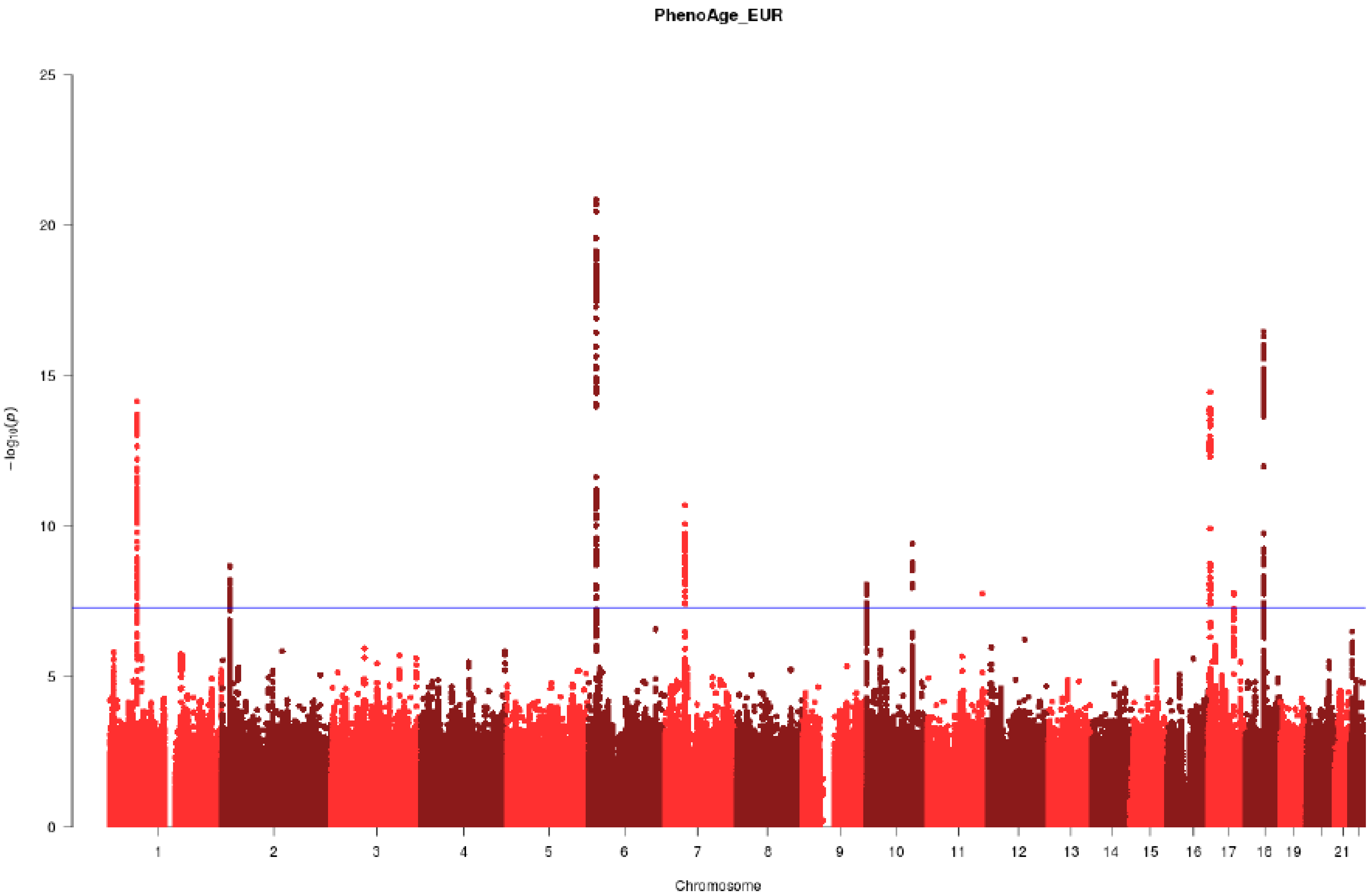


**Figure S19:** Manhattan Plot for PhenoAge Acceleration proportions in the European ancestry GWAS meta-analysis.

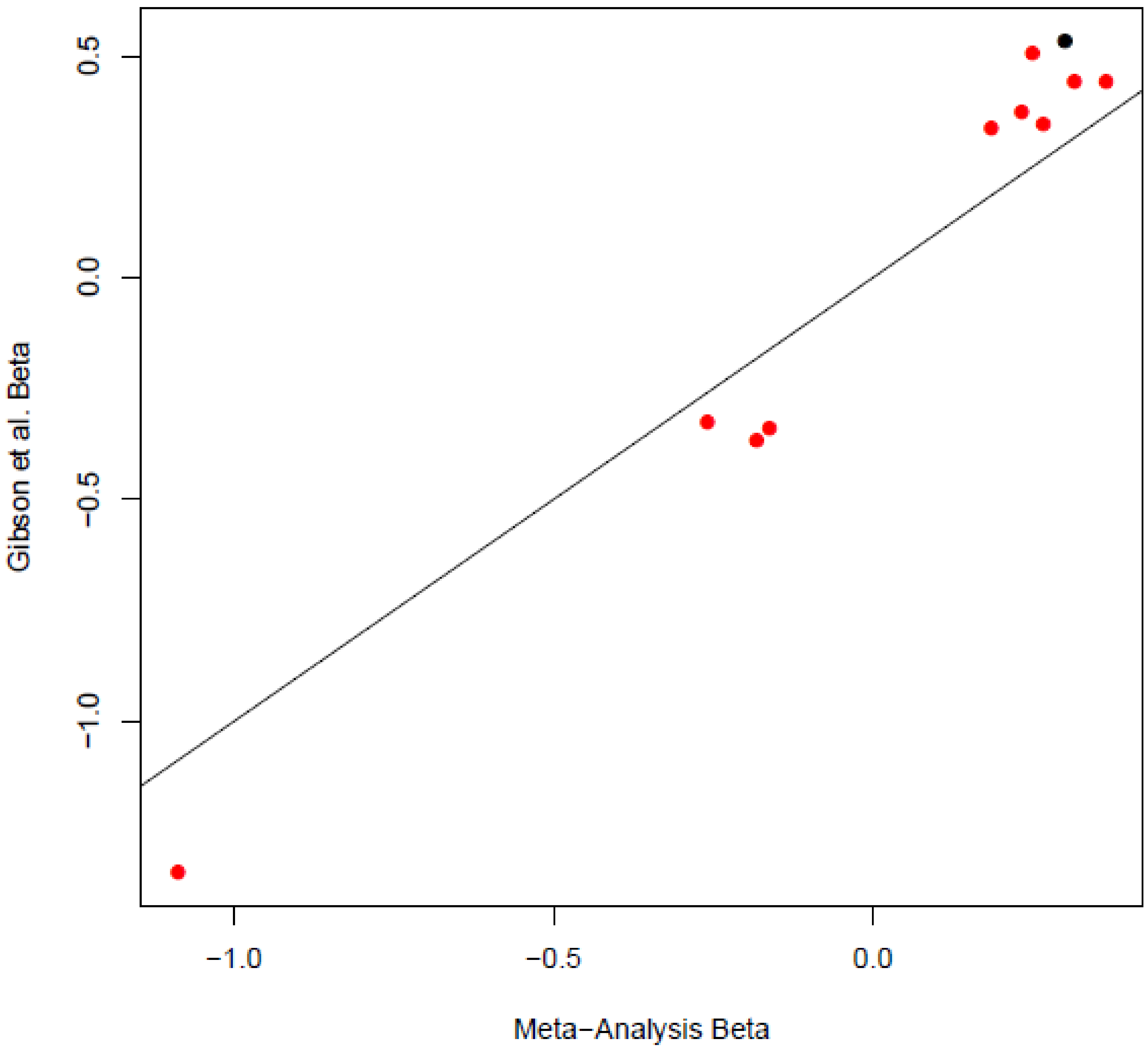


**Figure S20:** Plot of effect sizes for genome-wide significant SNPs in Gibson et al. vs effect sizes in a lookup of the current meta-analysis results for Hannum Age Acceleration and IEAA.

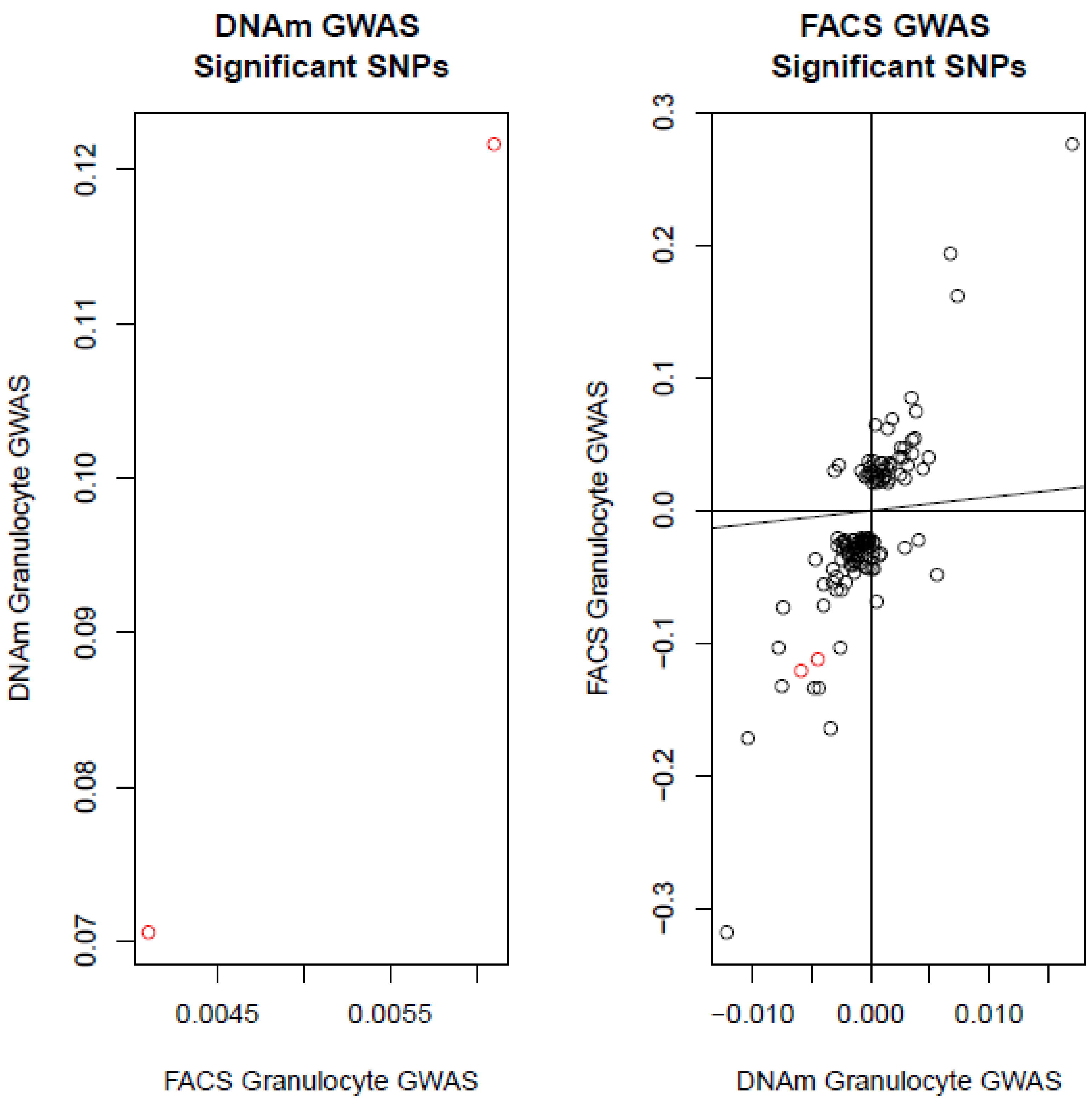


**Figure S21:** Plot of effect sizes for genome-wide significant SNPs in Astle et al. vs effect sizes in a lookup of the current meta-analysis results for granulocyte proportion (and vice versa).

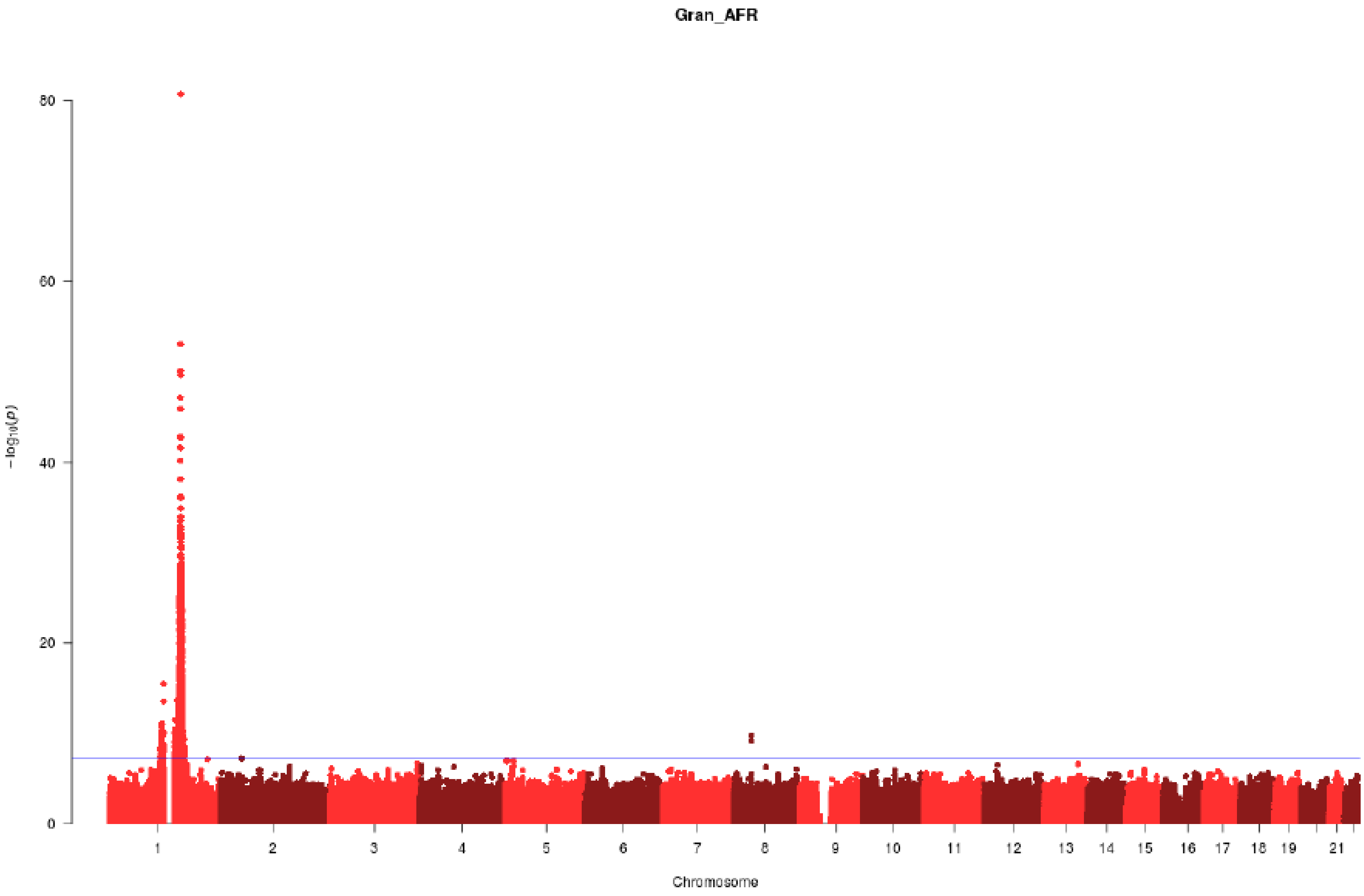


**Figure S22:** Manhattan Plot for granulocyte proportions in the African American GWAS meta-analysis.

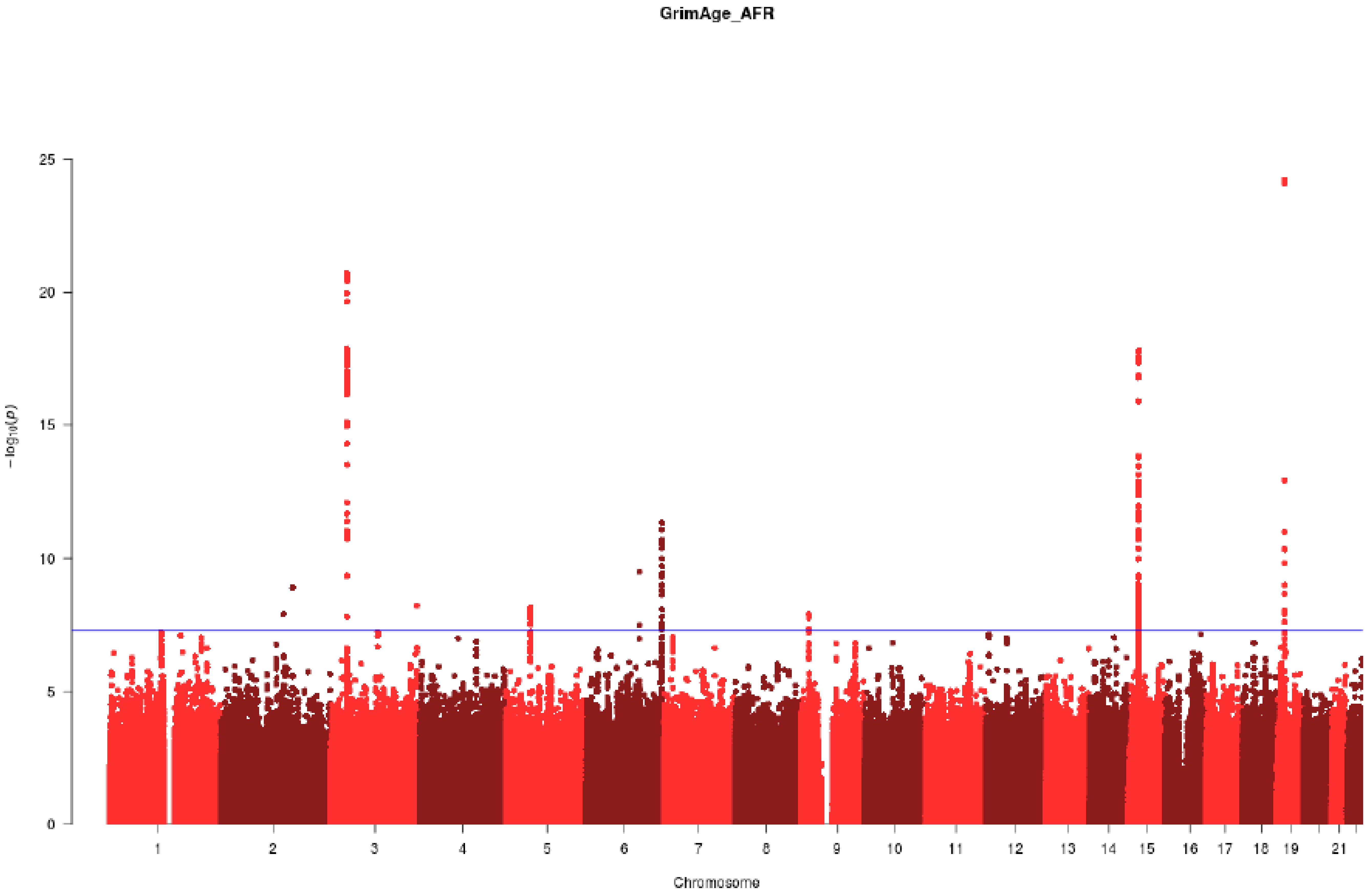


**Figure S23:** Manhattan Plot for GrimAge Acceleration in the African American GWAS meta-analysis.

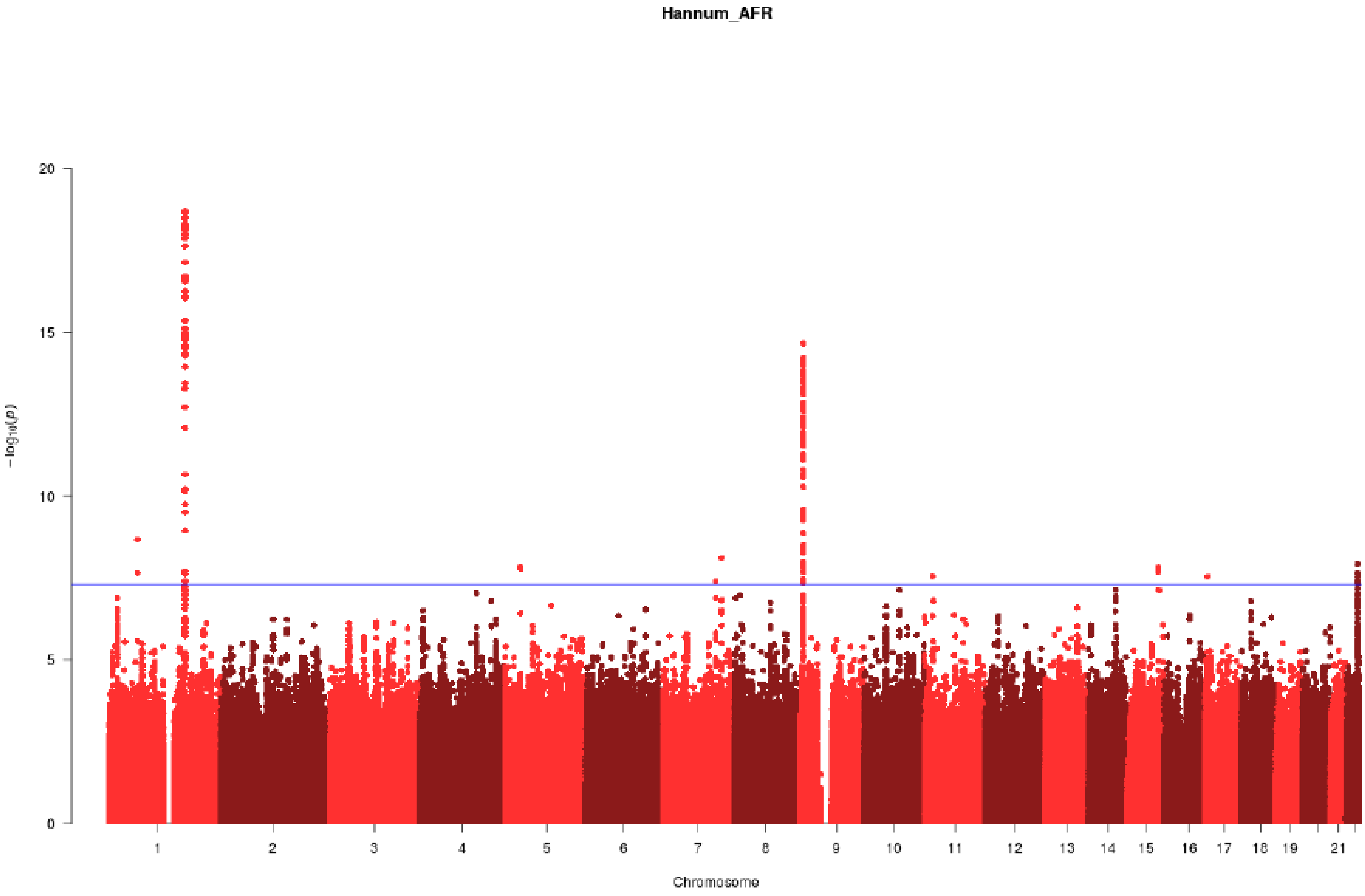


**Figure S24:** Manhattan Plot for Hannum Age Acceleration in the African American GWAS meta-analysis.

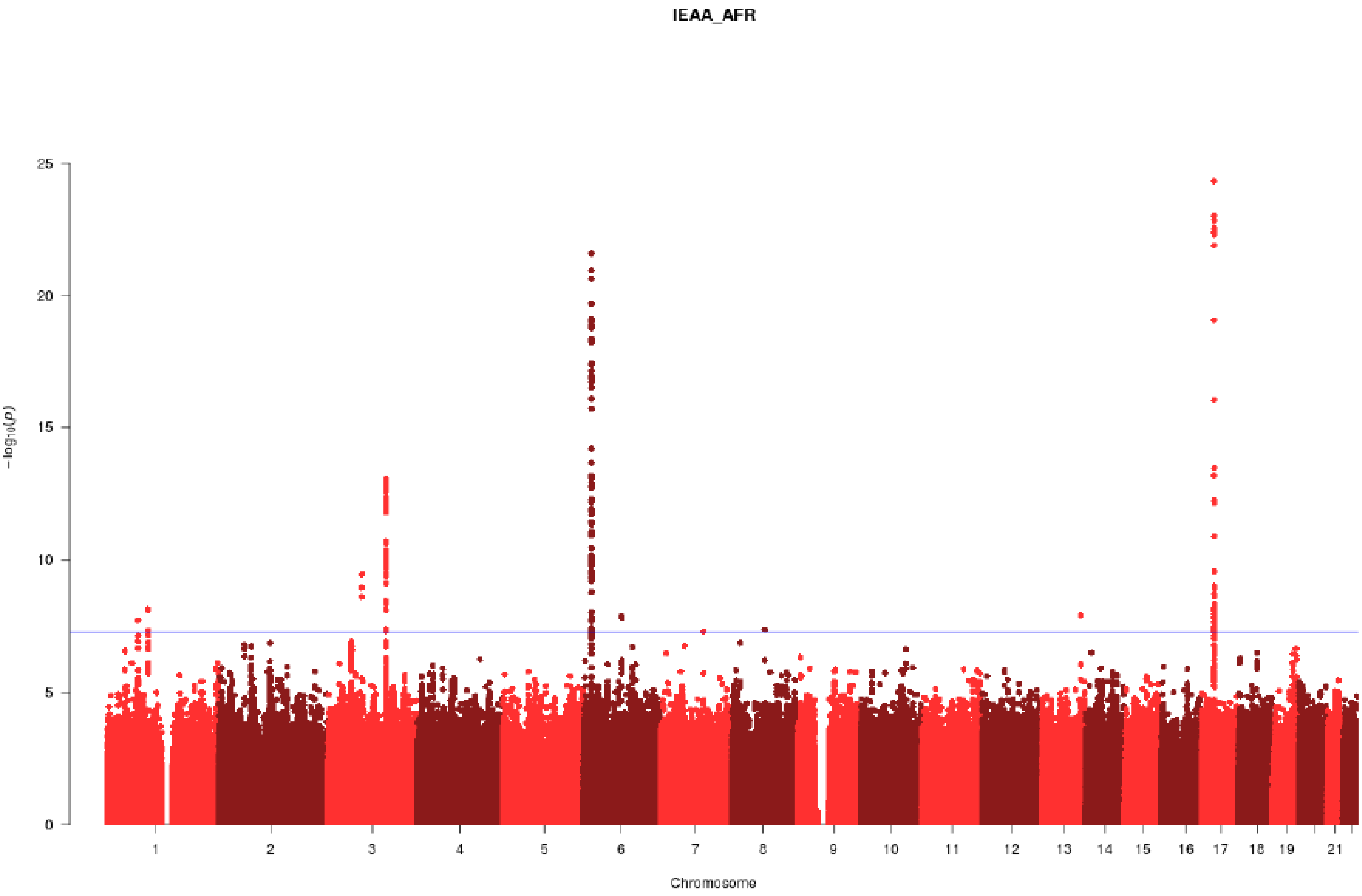


**Figure S25:** Manhattan Plot for IEAA in the African American GWAS meta-analysis.

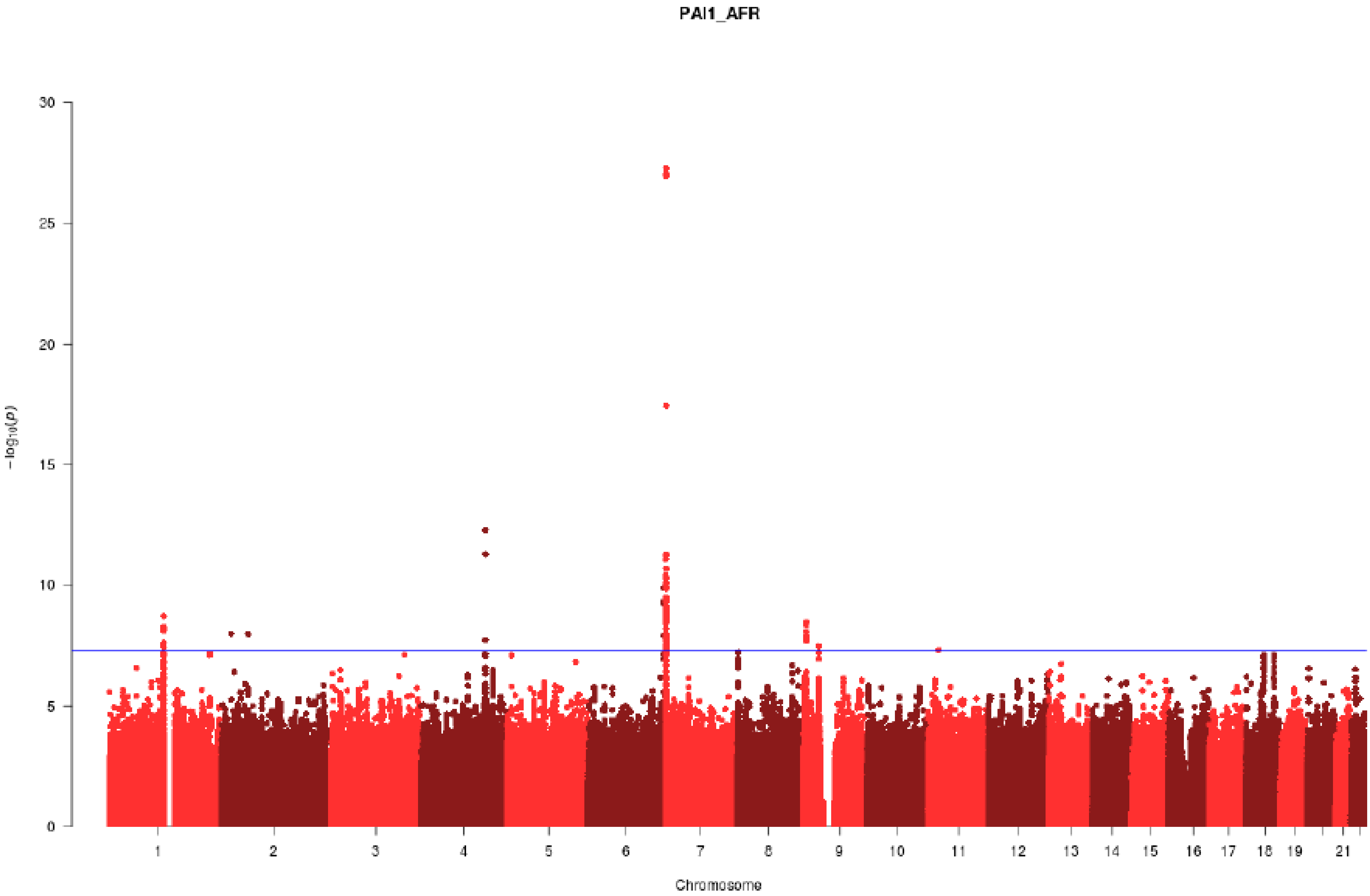


**Figure S26:** Manhattan Plot for PAI1 in the African American GWAS meta-analysis.

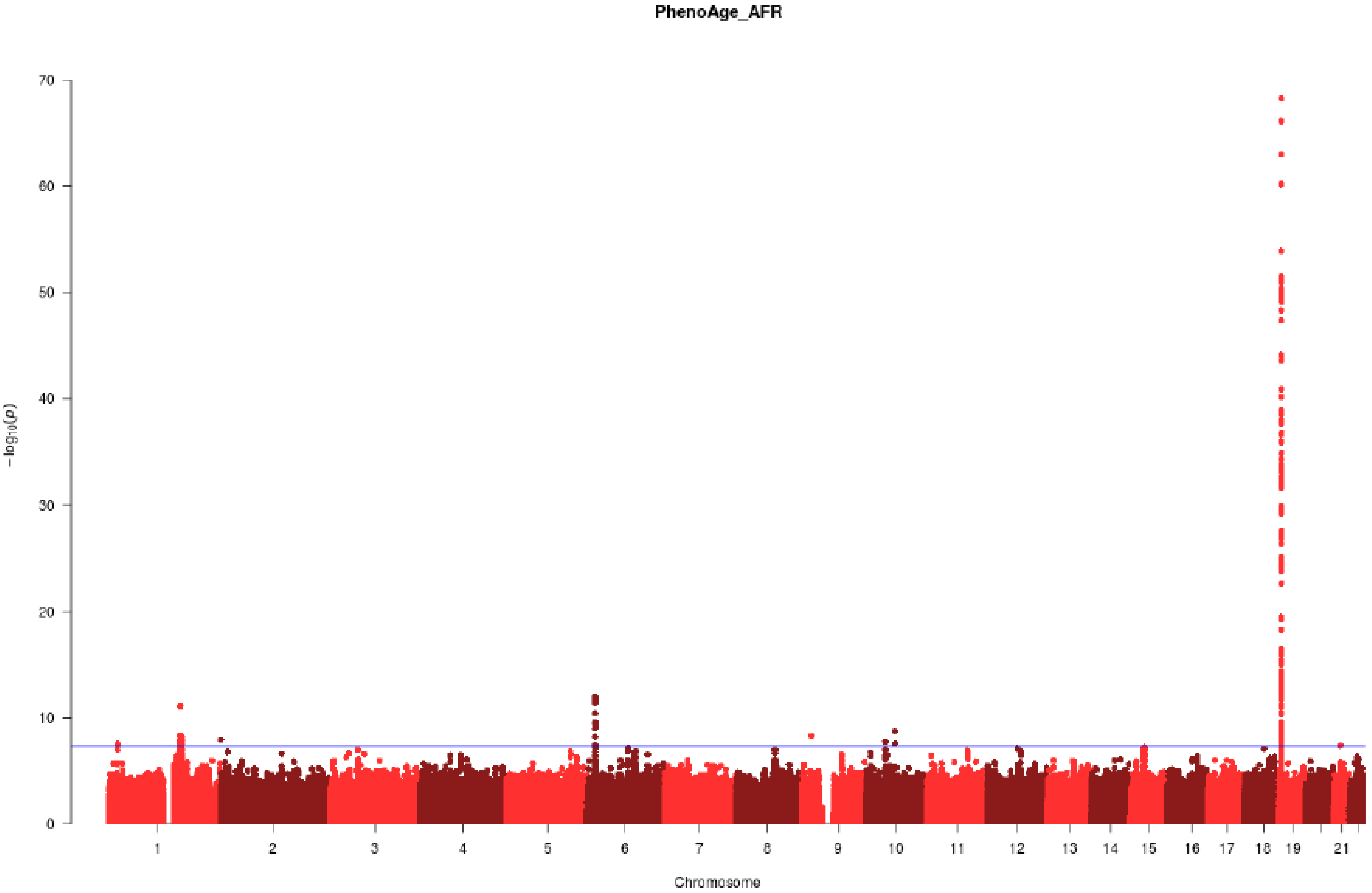


**Figure S27:** Manhattan Plot for PhenoAge Acceleration proportions in the African American GWAS meta-analysis.

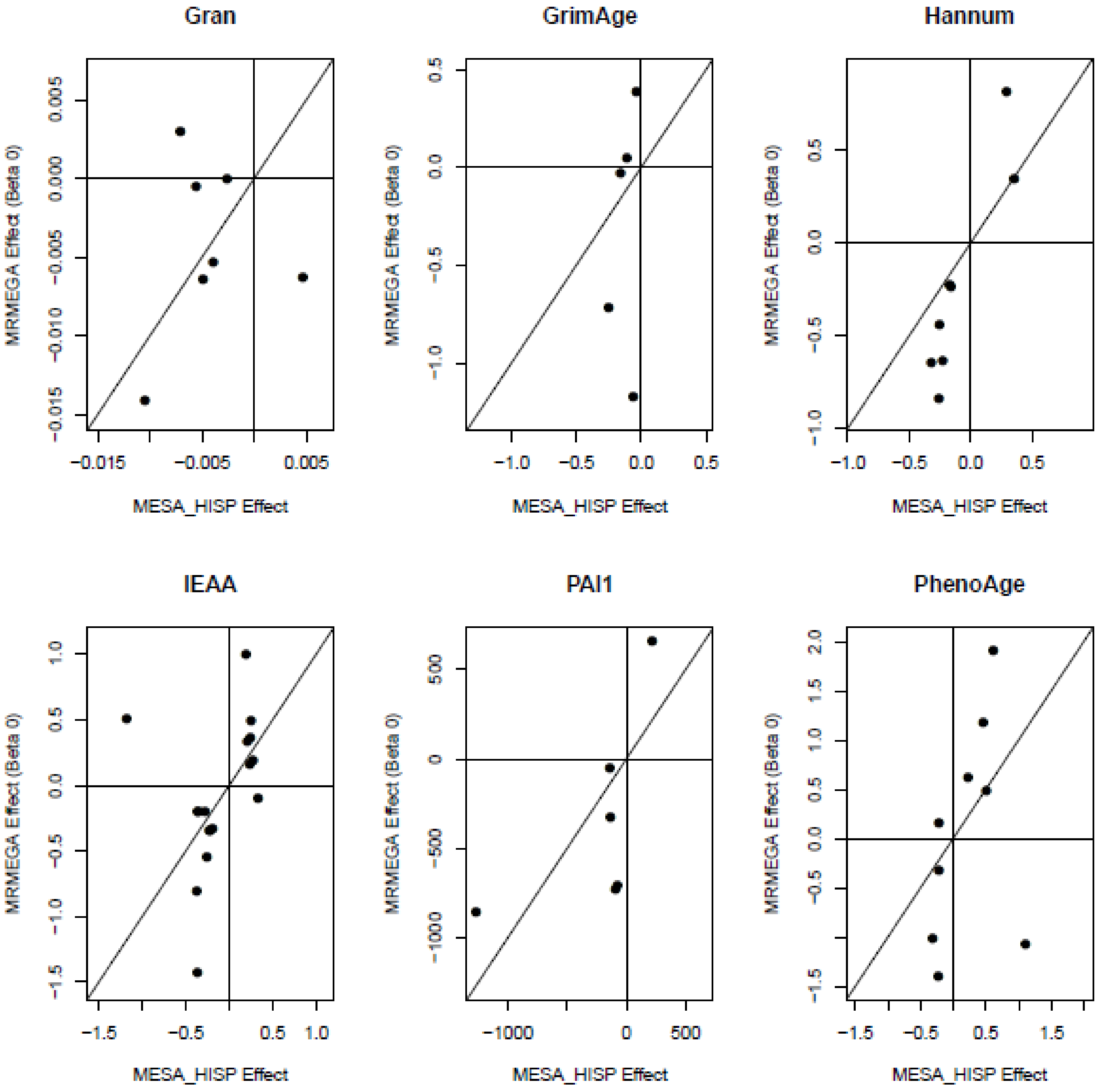


**Figure S28:** Plot of lead African- and European ancestry trans-ethnic meta-analysis SNP effect sizes against the same SNPs in the Hispanic subset of the MESA cohort.

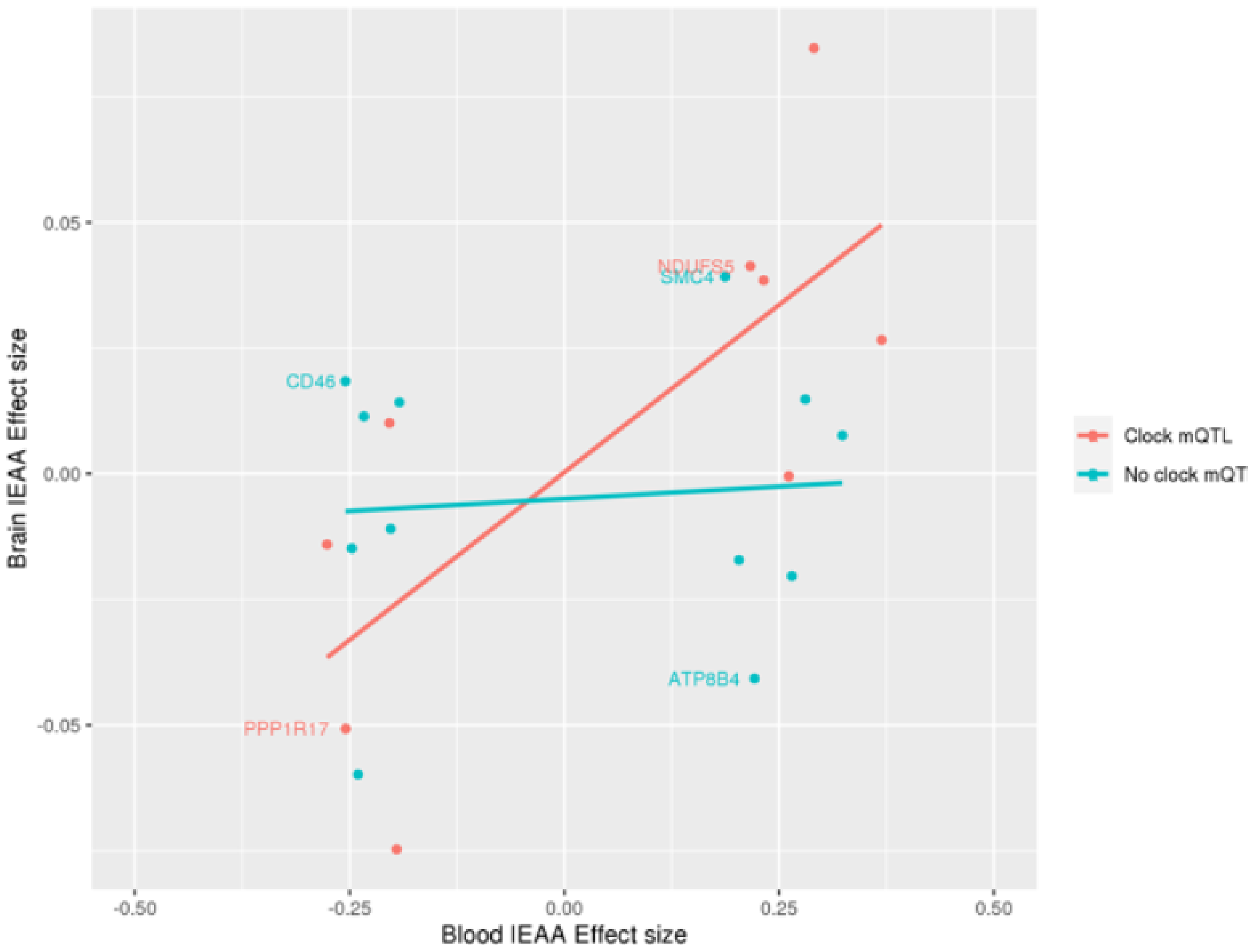


**Figure S29:** Lookup of 24 blood-based independent genome-wide significant SNPs for IEAA in a brain-based GWAS of IEAA (overlap of 21 SNPs). The red line represents the linear regression line (r=0.74) for SNPs that are also mQTLs for IEAA clock CpG sites. The turquoise line represents the linear regression line (r=0.08) for SNPs that are not mQTLs for IEAA clock CpGs. Labelled points correspond to loci where there was strong evidence of SNPs sharing genetic effects with eQTLs.

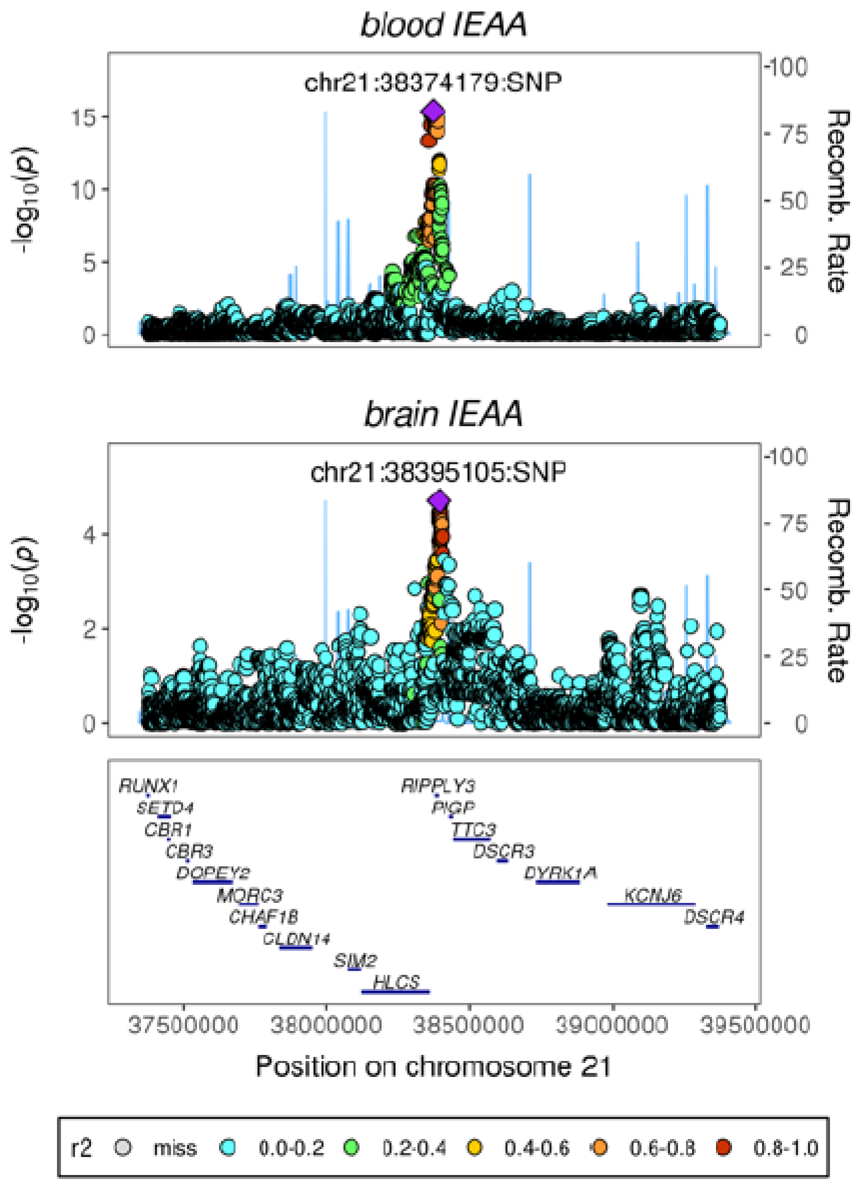


**Figure S30:** LocusZoom plot for the region (*DSRC6*/*RIPPLY3*) with highest evidence of genetic colocalization for the blood- and brain-based GWASs. Note that the lead SNP from the blood-based GWAS also colocalizes with a mQTL for an IEAA clock CpG, cg13450409 (PP=0.99, **Table S11**).

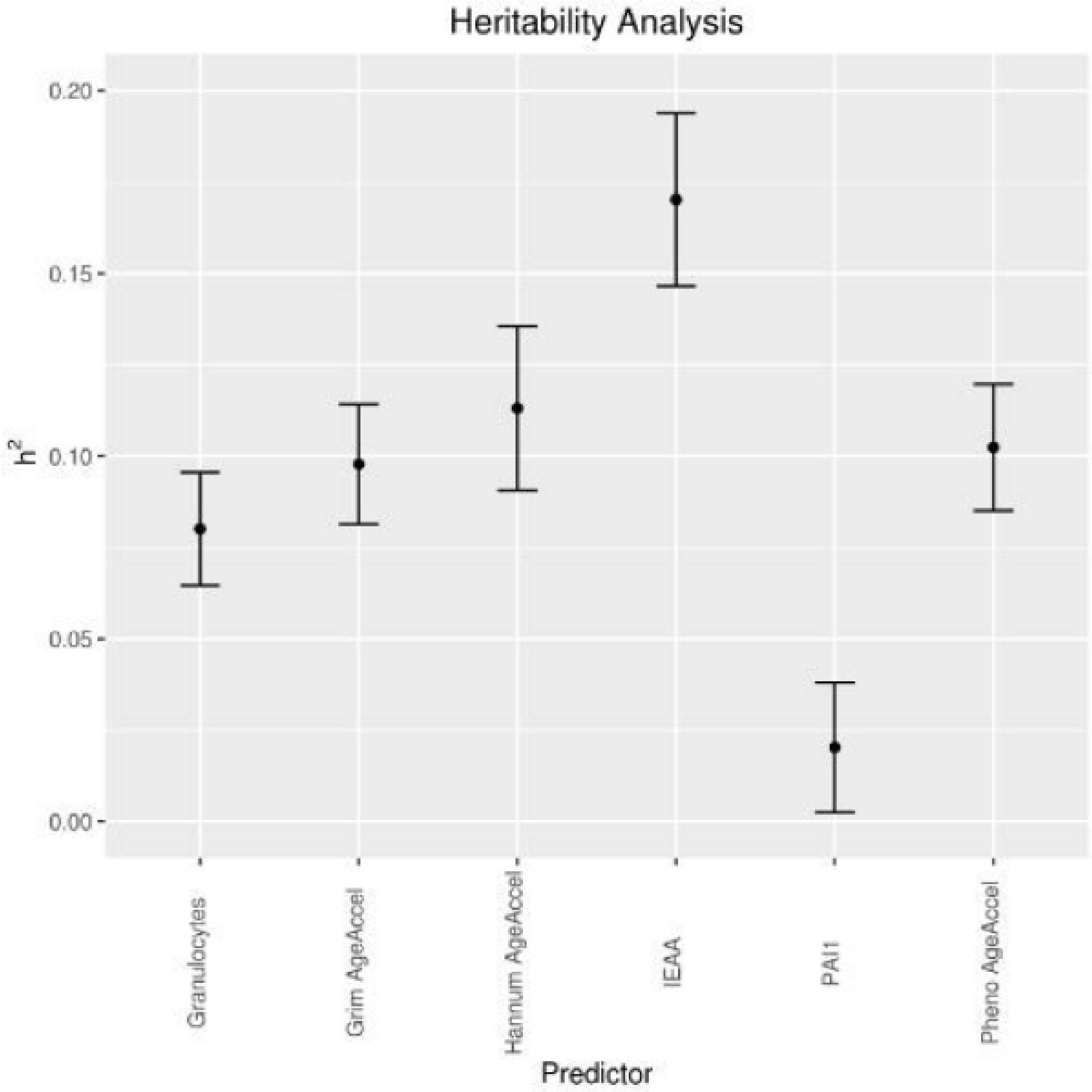


**Figure S31:** LD regression SNP-based heritability estimates for the six epigenetic biomarkers.
