## Appendix 1: Cohort Descriptions for "Genome-wide association studies identify 137 loci for DNA methylation biomarkers of ageing"

**Accessible Resource for Integrative Epigenomic Studies (ARIES)**

*Cohort description*

Samples were drawn from the Avon Longitudinal Study of Parents and Children [1, 2]. Pregnant women resident in Avon, UK with expected dates of delivery 1st April 1991 to 31st December 1992 were invited to take part in the study. The total number of pregnancies enrolled is 15,454 pregnancies, resulting in 15,589 foetuses. Of these 14,901 were alive at 1 year of age. Please note that the study website contains details of all the data that is available through a fully searchable data dictionary and variable search tool and reference the following webpage:

<http://www.bristol.ac.uk/alspac/researchers/our-data/>.

Blood from 1018 mother–child pairs (children at three time points and their mothers at two time points) were selected for analysis as part of the Accessible Resource for Integrative Epigenomic Studies (ARIES, <http://www.ariesepigenomics.org.uk/>) [3]. Following DNA extraction, samples were bisulphite converted using the Zymo EZ DNA Methylation™ kit (Zymo, Irvine, CA, USA).

*Data availability*

Data are available to researchers by request from the Avon Longitudinal Study of Parents and Children Executive Committee (<http://www.bristol.ac.uk/alspac/researchers/access/>) as outlined in the study's access policy <http://www.bristol.ac.uk/media-library/sites/alspac/documents/researchers/data-access/ALSPAC_Access_Policy.pdf>.

*Ethics*

Written informed consent has been obtained for all ALSPAC participants. Ethical approval for the study was obtained from the ALSPAC Ethics and Law Committee and the Local Research Ethics Committees.

*DNA Methylation*

Following conversion, genome-wide DNAm was measured using the Illumina Infinium HumanMethylation450 (HM450) BeadChip. The arrays were scanned using an Illumina iScan, with initial quality review using GenomeStudio. ARIES was preprocessed and normalised using the *meffil* R package [4]. ARIES consists of 5469 DNAm profiles obtained from 1022 mother-child pairs measured at five time points (three time points for children: birth, childhood and adolescence; and two for mothers: during pregnancy and at middle age). Low quality profiles were removed from further processing, and the remaining 4593 profiles were normalised using the Functional Normalization algorithm [5] with the top 10 control probe principal components (PCs). Full details of the preprocessing and normalization of ARIES has been described previously [4].

*Genotyping, imputation and quality control*

The ARIES participants were previously genotyped as part of the larger ALSPAC study, with QC, cleaning and imputation performed at the cohort level before extraction of the subset comprising ARIES. Mothers were genotyped using the Illumina Human660W-quad genome-wide SNP genotyping platform (Illumina Inc., San Diego, CA, USA) at the Centre National de Génotypage (CNG; Paris, France). Individuals were excluded based on non-European ancestry, missingness, relatedness, gender mismatches and heterozygosity. Additionally SNPs with a minor allele frequency of less than 1% were removed. PLINK (v1.07) [6] was used to carry out QC measures on an initial set of 10,015 subjects (including non-ARIES ALSPAC participants) and 557,124 directly genotyped SNPs. Following QC, the final directly genotyped dataset contained 526,688 SNP loci and 9,048 participants. Imputation was performed to increase the SNP density for all genotyped mothers and children combined. Genotypes were phased together using SHAPEIT (version 2, revision 727) [7] and then imputed against the HRC reference panel (pre-release 2015) [8] using IMPUTE v3. This gave 8,237 eligible children and 8,196 eligible mothers with available genotype data after exclusion of related subjects using cryptic relatedness measures. Association tests were adjusted for 20 genetic PCs using SNPTEST v2.5.2. We used genotype dosages and applied additive models.

*Acknowledgements*

We are extremely grateful to all the families who took part in the ALSPAC study, the midwives for their help in recruiting them, and the whole ALSPAC team, which includes interviewers, computer and laboratory technicians, clerical workers, research scientists, volunteers, managers, receptionists and nurses. 450K DNAm array data and part of the genotype data was generated in the Bristol Bioresource Laboratory Illumina Facility, University of Bristol.

*Funding*

This work was supported by the UK Medical Research Council; Wellcome ([www.wellcome.ac.uk](http://www.wellcome.ac.uk); [grant number 102215/2/13/2 to ALSPAC]); the University of Bristol to ALSPAC; the UK Economic and Social Research Council ([www.esrc.ac.uk](http://www.esrc.ac.uk); [ES/N000498/1] to CR) and the UK Medical Research Council ([www.mrc.ac.uk](http://www.mrc.ac.uk); grant numbers [MC_UU_00011/1, MC_UU_00011/5 to JLM, GH, GDS, CLR and MS]). DNAm data in the ALSPAC cohort were generated as part of the UK BBSRC funded (BB/I025751/1 and BB/I025263/1) Accessible Resource for Integrated Epigenomic Studies (ARIES, [http://www.ariesepigenomics.org.uk](http://www.ariesepigenomics.org.uk/)) and was funded by MC_UU_00011/5 to CLR. CLR, DAL, GH, JLM, HE and RCR are members of the UK Medical Research Council Integrative Epidemiology Unit at the University of Bristol (MC_UU_00011/5).

GWAS data was generated by Sample Logistics and Genotyping Facilities at Wellcome Sanger Institute and LabCorp (Laboratory Corporation of America) using support from 23andMe. GWAS data in the mothers was funded by Wellcome WT088806. This publication is the work of the authors and Josine Min will serve as guarantor for the contents of this paper.

**Airwave - The Airwave Health Monitoring Study**

*Cohort description*

Airwave - The Airwave Health Monitoring Study is an occupational cohort of employees of 28 police forces from across Great Britain. Full details of the cohort and methods are available in Elliott et al [1]. The study started recruitment in 2006 and now contains 53,280 participants. At the baseline health screening, participants underwent health examination, self-completed a computer questionnaire and blood samples were collected in EDTA tubes for DNA extraction.

*Ethics*

The study received ethical approval from the National Health Service Multi-Site Research Ethics Committee (MREC/13/NW/0588).

*DNA methylation*

For the microarray, bisulphite conversion of 500 ng of each DNA sample was performed using the EZ DNA Methylation-Lightning™ Kit according to the manufacturer’s protocol (Zymo Research, Orange, CA). Then, bisulfite-converted DNA was used for hybridization on the Infinium HumanMethylation EPIC BeadChip, following the Illumina Infinium HD Methylation protocol. Briefly, a whole genome amplification step was followed by enzymatic end-point fragmentation and hybridization to HumanMethylation EPIC BeadChips at 48°C for 17 h, followed by single nucleotide extension. The incorporated nucleotides were labelled with biotin (ddCTP and ddGTP) and 2,4-dinitrophenol (DNP) (ddATP and ddTTP). After the extension step and staining, the BeadChip was washed and scanned using the Illumina HiScan SQ scanner. The intensities of the images were extracted using the GenomeStudio (v.2011.1) Methylation module (1.9.0) software, which normalizes within-sample data using different internal controls that are present on the HumanMethylation EPIC BeadChip and internal background probes. The methylation score for each CpG was represented as a β-value according to the fluorescent intensity ratio representing any value between 0 (unmethylated) and 1 (completely methylated).

DNA methylation (DNAm) data were pre-processed and normalized using in-house software written for the R statistical computing environment, including background and color bias correction, quantile normalization, and Beta MIxture Quantile dilation (BMIQ) procedure to remove type I/type II probes bias, as described elsewhere [10]. DNAm levels were expressed as the ratio of the intensities of methylated cytosines over the total intensities (β values). Cross-reactive and polymorphic probes - with minor allele frequency greater than 0.01 in Europeans [11] - were excluded. Methylation measures were set to missing if the detection p-value was greater than 0.01. Samples with the bisulfite conversion control fluorescence intensity lower than 10,000 for both type I and type II probes and those with total call rate lower than 95% were excluded. Finally, samples were excluded if the predicted sex (based on chromosome X methylation) did not match that self-reported.

*Genotyping, imputation and quality control*

Genotyping was performed on the Illumina Infinium HumanCoreExome-12v1-1 BeadChip and quality control filters including call rate (>=97%), heterozygosity rate (<=3SD from the mean) were applied on the samples. Duplicated and second-degree relatives were further excluded and 14,062 samples of European ancestry based on principle component analysis remained. Markers were removed for high missing rate (>2%), deviation from Hardy-Weinberg equilibrium (P<1E-5) or minor allele frequency below 1%, resulting in 254,027 high-quality and common markers. Imputation was performed using the Haplotype Reference Consortium (HRC) panel (version r1.1 2016).

*Data availability*

All AIRWAVE data may be accessed upon application to Dementias Platform UK Data portal (https://portal.dementiasplatform.uk/Apply/ApplicationProcess). Additionally, the DNA methylation data is available for download from the Gene Expression Omnibus (GEO) repository at https://www.ncbi.nlm.nih.gov/geo/query/acc.cgi?acc=GSE147740.

*Acknowledgements*

OR was supported by an UK Research and Innovation Future Leaders Fellowship (MR/S03532X/1). This study was partly supported by the European Commission grant to the LIFEPATH project (Horizon 2020 grant number 633666). The Airwave Health Monitoring Study is funded by the Home Office (grant number 780- TETRA) with additional support from the National Institute for Health Research (NIHR) Biomedical Research Centre. The Airwave Study uses the computing resources of the UK MEDical BIOinformatics partnership (UK MED-BIO supported by the Medical Research Council (MR/L01632X/1). We thank all Airwave participants for their contributions.

**Atherosclerosis Risk in Communities (ARIC) Study**

*Cohort description*

The ARIC study is a prospective longitudinal investigation of the development of atherosclerosis and its clinical sequelae in which 15,792 individuals aged 45 to 64 years were enrolled at baseline (1987-1989). Details about the ARIC study have been previously reported [12]. Briefly, participants were selected by probability sampling from four communities in the United States: Forsyth County, North Carolina; Jackson, Mississippi (African-Americans only); suburban Minneapolis, Minnesota; and Washington County, Maryland. Six examinations have been completed (exam 1, 1987-1989; exam 2, 1990-1992; exam 3, 1993-1995; exam 4, 1996-1998; exam 5, 2011-2013; exam 6, 2016-2018) and a seventh is underway. Participants were contacted annually to update their medical histories.

*Ethics*

Written informed consent was provided by all study participants, and the study design and methods were approved by institutional review boards at each of the collaborating medical institutions: University of Mississippi Medical Center Institutional Review Board (Jackson Field Center); Wake Forest University Health Sciences Institutional Review Board (Forsyth County Field Center); University of Minnesota Institutional Review Board (Minnesota Field Center); and Johns Hopkins University School of Public Health Institutional Review Board (Washington County Field Center).

*DNA Methylation*

DNA methylation from whole blood was assessed using the Illumina HM450 array. To date, the arrays have been run on two batches of the ARIC data: the African-American cohort and a subset of the cohort of white participants with brain MRI data at exam 3. In QC step, a sample was removed if (1) having conflicting predicted sex, or (2) more the 5% of its CpG sites failed detection (using the pvalue cutoff of 0.01). A CpG site was removed if (1) more than 5% of samples with a beadcount <3, or (2) more than 5% of samples having fail detection (using the pvalue cutoff of 0.01). The raw probe signal was processed with Minfi’s preprocesses Noob() function [13], and were further normalized using BMIQ[14], before submission to the online DNA methylation age calculator [15].

*Genotyping, imputation and quality control*

Genotyping was performed using the Affymetrix® Genome-Wide Human SNP Array 6.0. QC procedures have been previously reported. Briefly, samples were removed in data cleaning procedures if they had an insufficient call rate, sex mismatch, discordance with previously-genotyped markers, first-degree relative of an included individual, and genetic outlier based on allele sharing and principal components analyses. Imputation was performed on the QCed data in two steps: (1) Pre-phasing with ShapeIt (v1.r532 ) (2) Imputation with IMPUTE2 using the 1000 Genomes Phase I v3 reference panel.

*Data availability*

The ARIC data are available from dbGaP under accession number phs000280 and from the ARIC study coordinating center at <https://sites.cscc.unc.edu/aric/distribution-agreements>.

*Acknowledgements*

The Atherosclerosis Risk in Communities study has been funded in whole or in part with Federal funds from the National Heart, Lung, and Blood Institute, National Institutes of Health, Department of Health and Human Services (contract numbers HHSN268201700001I, HHSN268201700002I, HHSN268201700003I, HHSN268201700004I and HHSN268201700005I), R01HL087641, R01HL086694; National Human Genome Research Institute contract U01HG004402; and National Institutes of Health contract HHSN268200625226C. Funding was also supported by 5RC2HL102419 and R01NS087541. Infrastructure was partly supported by Grant Number UL1RR025005, a component of the National Institutes of Health and NIH Roadmap for Medical Research. The authors thank the staff and participants of the ARIC study for their important contributions.

**Baltimore Longitudinal Study of Aging (BLSA)**

*Cohort description*

The Baltimore longitudinal study on Aging (BLSA) study is a population-based study aimed to evaluate contributors of healthy aging in the older population residing predominantly in the Baltimore-Washington DC area [16, 17]. Starting in 1958, participants are examined every one to four years depending on their age. Currently there are approximately 1,100 active participants enrolled in the study. Blood samples were collected for DNA extraction, and genome-wide genotyping was completed for 848 subjects of European descent was completed using Illumina 550K. Overnight fasted blood samples were used for genomic DNA extraction, and measurement of genetic variation and DNA methylation.

*Ethics*

The study protocol for BLSA were reviewed and approved by the Internal Review Board of the National Institute for Environmental Health Sciences (NIEHS) and all participants provided written informed consent.

*DNA Methylation*

DNA methylation was assayed in DNA samples collected at visits between November 1993 and March 2010 from participants who also had genome-wide SNP array or concurrent neuroimaging data available, where only array data was available for a participant the most recent DNA sample was selected. Analyses for BLSA included data from 507 participants with both DNA methylation and genotyping information. Genomic DNA was bisulfite converted using Zymo EZ-96 DNA Methylation Kit (Zymo Research Corp., Irvine, CA) per the manufacturer's protocol. CpG methylation status of 485,577 CpG sites was determined using the Illumina Infinium HumanMethylation450 BeadChip (Illumina Inc., San Diego, CA) per the manufacturer's protocol. Prior to selection of the study sample for the present analyses quality filtering and normalization were performed across all BLSA DNA methylation samples. Samples identified as outliers by multi-dimensional scaling (> 3 standard deviations from the mean) or with high rates of single nucleotide polymorphism (SNP) mismatch (≥5 SNPs) were excluded. NOOB background correction and BMIQ normalization were performed using the wateRmelon package [16]. Background and quantile normalization were applied to the filtered dataset. The data that passed quality control were uploaded to Horvath’s online calculator as described in the main text, to generate the various clock estimates.

*Genotyping, imputation and quality control*

Genome-wide genotyping was completed using Illumina Infinium HumanHap 550K SNP arrays were used for genotyping [19]. Genotyping was completed for 1210 subjects with a sample call rate >98.5%, and correct sex specification. 501,704 autosomal SNPs that passed quality control (MAF>1%, completeness >99%, HWE > 10-4) were used for imputation. The QC’d SNP data were imputed to the HRC reference panel (v1.1) using the Michigan imputation server [20]. GWAS was run using PLINK 1.9 [6].

*Data availability*

The BLSA data are available upon request. Applications should be made through the website: https://www.blsa.nih.gov/

*Acknowledgements*

This work was supported in part by the Intramural Research Program of the National Institute on Aging, National Institutes of Health, Baltimore, Maryland. This work utilized the computational resources of the NIH HPC Biowulf cluster. (http://hpc.nih.gov)

**Bogalusa Heart Study (BHS)**

*Cohort description*

The BHS is a long-term epidemiology study of the natural history of cardiovascular disease from childhood to adulthood in the community of Bogalusa (65% white, 35% African American), Louisiana, the United States (Berenson GS. Bogalusa Heart Study: a long-term community study of a rural biracial (Black/White) population. [21]. In the current study, 821 participants (56.6% women) were included who had data on DNA methylation from the Infinium HumanMethylation450K BeadChip (Illumina, San Diego, CA) and genotype data from the Illumina MetaboChip (Illumina, San Diego, CA).

*Ethics*

All participants in BHS provided written informed consents. Study protocols of BHS were approved by the Institutional Review Boards of Tulane University Health Sciences Center.

*DNA Methylation*

Genomic DNA was isolated from whole blood samples in the BHS using the FlexiGene DNA kit (Qiagen). The Infinium HumanMethylation450K BeadChip (Illumina, San Diego, CA) was used for whole-genome DNAm analysis. Samples were processed at the Microarray Core Facility Lab, University of Texas Southwestern Medical Center, Dallas, TX, USA. For each subject, 750ng genomic DNA was bisulfite converted using the 96 well EZ DNAm kit (Zymo Research, Irvine, CA) according to manufacturer’s instructions. The efficiency of the bisulfite conversion was confirmed by in-built controls on the 450K array. The methylation profile of each participant was measured by processing 4μl of bisulfite-converted DNA, at a concentration of 50ng/μl, on a 450K array. The bisulfite converted DNA was amplified, fragmented and hybridized to the array following the protocol. We scanned the arrays by using an Illumina iScan scanner, and then the raw methylation data was extracted using Illumina’s Genome Studio Methylation Module. The BHS used the data-driven separate normalization (dasen) from the wateRmelon R [16]. The probe exclusion criteria for filtering samples and probes included: 1) samples having 1% of CpG sites with a detection p-value greater than 0.05; 2) probes having 5% of samples with a detection p-value greater than 0.05; and 3) probes with beadcount less than 3 in 5% of the samples.

*Genotyping, imputation and quality control*

Individuals were genotyped on the Illumina MetaboChip according to the manufacturer’s instructions and protocol. Samples were processed at the Microarray Core Facility Lab, University of Texas Southwestern Medical Center, Dallas, TX, USA. The online tool of Michigan Imputation Server was used for genotype imputation with 1000G Phase3 v5 as the reference panel, with the following criteria for SNPs filtering: 1) r2≥0.3; 2) minor allele frequency≥0.05; 3) sample call rate≥0.95;4) probe call rate≥0.95; and 5) Hardy-Weinberg Equilibrium Test p>1.0E-6.

*Data availability*

Individual level genotype, epigenome and phenotype data from BHS are not permitted to be shared or deposited. The summary data that support the findings of the present study are available from the Bogalusa Heart Study Steering Committee on reasonable request. Application should be made to through online request (https://www.clersite.org/contact/).

*Acknowledgements*

BHS was supported by grants R01AG016592 and R03AG060619 from National Institute of Aging. We thank the participants and investigators and staff members of the BHS for their outstanding commitment and cooperation.

**Born in Bradford (BiB)**

*Cohort description*

BiB is a population-based prospective birth cohort including 12,453 women across 13,776 pregnancies, who were booked for antenatal care at the Bradford Royal Infirmary (the only maternity hospital in the city of Bradford). Full details of the study methodology have been reported previously [22]. In brief, most women were recruited at their oral glucose tolerance test (OGTT) at approximately 26-28 weeks gestation, which is offered to all women booked for delivery at Bradford Royal Infirmary. Eligible women had to have an expected delivery between March 2007 and December 2010. Bradford is a deprived multi-ethnic city in the North of the UK with ~5,000 deliveries per year. Approximately half of the births are to mothers of South Asian origin, with most of the remainder being of White European origin [23]. DNA was extracted from whole blood taken from women during pregnancy (mothers) and whole cord blood taken after delivery of the infant (offspring).

*Ethics*

Ethical approval for the study was granted by the Bradford National Health Service Research Ethics Committee (ref 06/Q1202/48), and all participants gave written informed consent.

*DNA Methylation*

DNA methylation from whole blood was assessed using the Illumina EPIC array. The arrays were run on 1000 mother offspring pairs. A total of 500 ng high molecular weight DNA was bisulfite-converted using the EZ-96 DNA methylation kit (Zymo Research, Orange, CA, USA). DNAm was quantified using Illumina HumanMethylation EPIC Arrays (Illumina, San Diego, CA, USA). During the data generation process, a wide range of batch variables were recorded in a purpose-built laboratory information management system (LIMS). Sample QC and normalization was performed using with *meffil* [4]. Samples failing QC (average probe p > 0.01) were excluded from further analysis. As an additional QC step genotype probes were compared with genotype data from the same individual to identify and remove any sample mismatches. Furthermore, samples failed on control probes (bisulfite 1 and bisulfite 2) were also excluded from the analysis. Finally, 864 samples passed QC. Samples were normalized using functional normalization [5] using *meffil* [4]. Data was normalised using 7 control probe PCs derived from the technical probes informed by *meffil* scree plots.

*Genotyping, imputation and quality control*

BiB maternal and offspring genotype data was obtained from an Illumina HumanCoreExome array or in a second batch an Infinium Global Screening array (GSA). Genotype data were imputed against HRC r1.1 using Minimac4, after quality control (MAF >1% and HWE>1×10-6). Association tests were adjusted for 20 genetic PCs using SNPTEST v2.5.2. We used genotype dosages and applied additive models.

*Data availability*

For access to BiB data the ‘Expression of Interest’ form is available at https://borninbradford.nhs.uk/wp-content/uploads/Expression-of-interest-proforma-v3_23.01.20.doc and should be submitted to.

*Acknowledgements*

We are grateful to everyone involved in the Born in Bradford study. This includes the families who kindly participated as well as the practitioners and researchers all of whom have made Born in Bradford happen. Sample processing, DNA extraction and running of Illumina assays were carried out at the Bristol Bioresource Laboratory at the University of Bristol.

*Funding*

BiB receives core funding from Wellcome (WT101597MA) a joint grant from the UK Medical Research Council (MRC) and UK Economic and Social Science Research Council (ESRC) (MR/N024397/1) and the National Institute for Health Research (NIHR) under its Collaboration for Applied Health Research and Care (CLAHRC) for Yorkshire and Humber. The research presented in this paper is supported by the British Heart Foundation (CS/16/4/32482, FS/17/60/33474, and AA/18/7/34219), US National Institute of Health (R01 DK10324), the European Research Council under the European Union's Seventh Framework Programme (FP/2007-2013) / ERC Grant Agreement (Grant number 669545; DevelopObese) and from the European Union’s Horizon 2020 research and innovation programme under grant agreement No 733206 (LifeCycle), and the NIHR Biomedical Centre at the University Hospitals Bristol NHS Foundation Trust and the University of Bristol. J.W. is funded by a UK Medical Research Council (MRC) Population Health Scientist Postdoctoral Award (MR/K021656/1). K.T., D.L.S.F. and D.A.L. work in a unit that receives UK MRC funding (MC_UU_00011/6) and D.A.L. is an NIHR senior investigator (NF-SI-0616-10102). The funders had no role in the design of the study, the collection, analysis, or interpretation of the data; the writing of the manuscript, or the decision to submit the manuscript for publication.

**The Danish Twin Registry (DTR)**

*Cohort description*

The Danish Twin Registry (DTR) sample included 375 individuals collected as part of the study of Middle-Aged Danish Twins (MADT, N=244) and the Longitudinal Study of Aging Danish Twins (LSADT, N=131) [24]. MADT was initiated in 1998 and includes 4,314 twins randomly chosen from the birth years 1931-1952. Surviving participants were revisited from 2008 to 2011, where the blood samples and survey data used in the present study were collected. LSADT was initiated in 1995 and includes twins aged 70 years and older. Follow-up assessments were conducted every second year through 2005. The individuals included here all participated in the 1997 assessment, where blood samples and survey data were collected from same sex twin pairs.

Methylation and genotype data were primarily available for complete monozygotic and dizygotic twin pairs, so to ensure a sample of unrelated individuals, one twin from each twin pair was randomly chosen to be included in the study.

*Ethics*

Written informed consents were obtained from all participants. Collection and use of biological material and survey information were approved by the Regional Scientific Ethical Committees for Southern Denmark (VF 20040241), and the study was approved by the Danish Data Protection Agency.

*DNA Methylation*

Genome-wide DNA methylation from whole blood was quantified using the Infinium HumanMethylation450 BeadChip (Illumina, San Diego, CA, USA). Quality control of the DNA methylation data was performed as described in [25]. Relevant DNAm estimates and age acceleration residuals were calculated as specified in the analysis plan using the provided scripts.

*Genotyping, imputation and quality control*

Samples were genotyped using the Illumina Infinium PsychArray (Illumina, San Diego, CA, USA). Genotyping was conducted by the SNP&SEQ Technology Platform, Science for Life Laboratory, Uppsala, Sweden (http://snpseq.medsci.uu.se/genotyping/snp-services/). Pre-imputation quality control included filtering SNPs on genotype call rate <98%, HWE *P*<10^-6^, and MAF=0, and individuals on sample call rate <99%, relatedness and gender mismatch. Pre-phasing and imputation to the 1000 Genomes phase 3 reference panel was performed using IMPUTE2 version 2.3.2 [26] and a chunk-size of 1 MB. The genome-wide association studies of the 6 age acceleration measures were run as specified in the analysis plan using the frequentist additive model and the expected method in SNPTEST v. 2.5.2 [27] and including sex as a covariate. Principal components were not included as all study sample individuals are of the same ancestry.

*Data availability*

According to Danish legislation, transfer and sharing of individual-level data require prior approval from the Danish Data Protection Agency and data sharing requests are dealt with on a case-by-case basis. For these reasons, the raw data cannot be deposited in a public database. However, we welcome any enquiries regarding collaboration and individual requests for data sharing.

*Acknowledgements*

DTR is supported by grants from The National Program for Research Infrastructure 2007 from the Danish Agency for Science, Technology and Innovation (09-063256) and the US National Institutes of Health (P01 AG08761). Genotyping was supported by NIH R01 AG037985 (Pedersen), while the DNA methylation analysis was supported by the European Union’s Seventh Framework Programme (FP7/2007-2011) under grant agreement n° 259679.

**Estonian Genome Center, University of Tartu (EGCUT)**

*Cohort description*

Estonian Genome Center at the University of Tartu (EGCUT) hosts the Estonian biobank [28] - a population-based biobank which comprises health, genealogical, and ‘omics’ data of close to 200,000 individuals ≥18 years of age, closely reflecting the age distribution of the adult Estonian population. All participants have completed a computer assisted interview, including personal data (place of birth, place(s) of living, nationality etc.), genealogical data (family history, three generations), educational and occupational history and lifestyle data (physical activity, dietary habits, smoking, alcohol consumption, quality of life). Anthropometric and physiological measurements have also been recorded. The samples used in this study were selected from the EGCUT Center for Translational Genomics (CTG) cohort of individuals who have been re-contacted for a second time-point sample. The collection of blood samples and data generation was conducted according to the Human Genes Research Act of Estonia.

*Ethics*

The study was approved by the Ethics Review Committee of Human Research of the University of Tartu, Estonia (permission no 206/T-4, date of issue 25.08.2011) and it was carried out in compliance with the Helsinki Declaration. All of the participants have signed a broad informed consent. All methods were carried out in accordance with approved guidelines.

*DNA Methylation*

DNA methylation was quantified from whole blood using the Illumina Infinium HumanMethylation450 array. Data pre-processing and quality control analyses were performed in R with the Bioconductor package *minfi* [29], using the original IDAT files extracted from the HiScanSQ scanner. Raw pre-processing was used to convert the intensities from the red and the green channels into methylated and unmethylated signals. Beta values were computed using Illumina’s formula [beta = M/(M + U + 100)]. Detection p-values were obtained for every CpG probe in each sample. Samples with detection p-value > 1E-16 in more than 5% of the CpG sites, outliers based on X and Y methylation patterns, and possibly contaminated or mixed up samples (determined using the 65 SNPs present on the methylation beadchip) were discarded. Beta-values of a subset of CpG sites were uploaded to the Horvath’s online epigenetic clock software to generate the various epigenetic clock estimates.

*Genotyping, imputation and quality control*

DNA from the samples were genotyped using HumanOmniExpress BeadChips (Illumina) at the Estonian Genome Center, according to the manufacturer’s instructions. Samples were excluded based on sample call-rate < 0.95 as well as sample heterozygosity test, check for population outliers and samples with wrong/unidentifiable sex. SNPs with marker call-rate < 0.95 and HWE p < 1E-06 were excluded. The dataset was imputed using the 1000 Genomes project phase 3 reference with IMPUTE v2 [26]. The GWAS was performed using SNPTEST v2.5.4 [27].

*Data availability*

Data of the subjects from the Estonian biobank are handled in accordance with the regulations of the Human Genes Research Act. Data can be accessed upon ethical approval by submitting a data release request to the Estonian Genome Center, University of Tartu (http://www.geenivaramu.ee/en/access-biopank/data-access).

*Acknowledgements*

EGCUT analyses were funded by the Estonian Research Council Grant PRG184, and the European Union through the European Regional Development Fund Project No. 2014-2020.4.01.15-0012 GENTRANSMED and 2014-2020.4.01.16-0125. The work was carried out in the High Performance Computing Center of the University of Tartu.

**Finnish Twin Cohort	(FTC)**

The FTC includes three large cohort studies 1) the older twin cohort of twins born before 1958, 2) Finntwin16, born in 1975-1979, and 3) Finntwin12, born in 1983-1987 [30, 31, 32]. All cohorts have been studied in an intensive longitudinal manner. All twins who had taken part in several clinical in-person studies with sampling for whole blood DNA and subsequent genotype and DNA methylation analyses were included.

*Ethics*

The FTC data collection has been approved by the ethics committees of the University of Helsinki (113/E3/01 and 346/E0/05) and Helsinki University Central Hospital (270/13/03/01/2008 and 154/13/03/00/2011). Written informed consent was provided by the participants before the sample collection.

*DNA Methylation*

Genome-wide DNAm was measured using Illumina’s Infinium HumanMethylation450 and the MethylationEPIC BeadChip, according to the manufacturer’s instructions (Illumina, San Diego, CA, USA). DNAm data were preprocessed using R package *minfi*. Detection p values comparing total signal for each probe to the background signal level, were calculated to evaluate quality of the data [33]. Probes of poor quality (mean detection p > 0.01) were excluded from further analysis. Beta values representing CpG methylation levels were calculated as ratio of methylated intensities (M) to the overall intensities (Beta value=M/(M+U+100), where U is unmethylated probe intensity). Resulting methylation data were uploaded to Horvath’s online calculator, to generate the various clock estimates. Samples were removed if predicted sex or tissue did not match with the reported sex or tissue. Outliers were filtered based on cut-off criteria > or < 5 standard deviations.

*Genotyping, imputation and quality control*

Genotyping was done using Illumina Human610-Quad v1.0 B and Human670-QuadCustom v1.0 arrays at the Wellcome Trust Sanger Institute (Cambridge UK), Illumina HumanCoreExome- (12 v1.0 A, 12 v1.1 A, 24 v1.0 A, 24 v1.1 A, 24 v1.2 A) arrays at the Broad Institute of MIT and Harvard (MA, USA), Wellcome Trust Sanger Institute (Cambridge, UK), University of Chicago Genomics Facility (Chicago IL, USA) and Institute for Molecular Medicine Finland (Helsinki, Finland) and with Affymetrix FinnGen Axiom array at Thermo Fisher Scientific (Santa Clara CA, USA). The algorithm for genotype calling were Illumina’s GenCall for all HumanCoreExome chip genotypes, Illuminus for 610k & 670k chip genotypes and AxiomGT1 for Affymetrix chip genotypes. On Illumina arrays where genotypes were called to Illumina’s TOP strand, strand were flipped to forward strand using strand files generated by Will Rayner (<https://www.well.ox.ac.uk/~wrayner/strand/>). The genome build of all genotypes were set to GRCh37/hg19. In case where genotypes were called to NCBI36/hg18 or GRCh38/hg38, genome positions were lifted to GRCh37/hg19 using University of California Santa Cruz LiftOver program [34] with appropriate chain file. Genotype quality control was done in three batches (batch1: 610k+670k, batch2: HumanCoreExome and batch3: Affymetrix chip genotypes). Variants with call rate below 97.5% (batch1 and batch3) or 95% (batch2), samples with call rate below 98% (batch1) or 95% (batch2 and batch3), and variants with minor allele frequency below 1% with Hardy-Weinberg Equilibrium p-value lower than 1e-06 were removed. In addition, samples from all batches with heterozygosity test method-of-moments F coefficient estimate value below -0.03 or higher than 0.05 (batch1 and batch2) or ±4SD from the mean (batch3) were removed along with the samples which failed sex check or were among the multi-dimensional scaling principal component analysis outliers. Total amount of genotyped autosomal variants after quality control were 475,526 (batch1), 239,894 (batch2) and 388,673 (batch3) with following number of samples remaining for imputation: 2,617 (batch1), 5,328 (batch2) and 8,218 (batch3). Pre-phasing was performed using Eagle v2.3 [35] and imputation with Minimac3 v2.0.1 using University of Michigan Imputation Server [20]. Genotypes of all batches were imputed to Haplotype Reference Consortium release 1.1 reference panel [8]. The final study sample were extracted from the data where all batches of imputed data were merged.

*Data availability*

FTC data are available at the Biobank of the National Institute for Health and Welfare, Finland. All the biobanked data are publicly available for use by qualified researchers following a standardized application procedure.

*Acknowledgements*

This work was supported by the Academy of Finland [213506, 265240, 263278, 312073 to JK, and 297908 to MO], EC FP5 GenomEUtwin (JK), NIH NIH/NHLBI (grant HL104125), EC MC ITN Project EPITRAIN (JK & MO) project and the University of Helsinki Research Funds to MO, Sigrid Juselius Foundation to JK and MO, Yrjö Jahnsson foundation (6868) and Juho Vainio foundation to ES. The authors thank all FTC study participants and research team members. The Gerontology Research Center is a joint effort between the University of Jyväskylä and the University of Tampere.

**The Framingham Heart Study (FHS)**

*Cohort description*

The Framingham Heart Study (FHS) [36] is a large-scale longitudinal study started in 1948, initially investigating the common factors of characteristics that contribute to cardiovascular disease (CVD), https://www.framinghamheartstudy.org/index.php. The study at first enrolled participants living in the town of Framingham, Massachusetts, who were free of overt symptoms of CVD, heart attack or stroke at enrolment. In 1971, the study started FHS Offspring Cohort to enrol a second generation of the original participants’ adult children and their spouses (n= 5,124) for conducting similar examinations [37]. Participants from the FHS Offspring Cohort were eligible for our GWAS study if they attended the eighth examination cycle (2005-2008) and consented to having their DNA to be used for genetic research. There are a total of 2,329 FHS participants with European genetic ancestry available for both DNA methylation and SNP array data. Multidimensional scaling (MDS) in conduction with principal components analysis was performed to identify genetic ancestry outliers, leaving 2,322 participants remained in analysis.

*Ethics*

All participants provided written informed consent at the time of each examination visit. The study protocol was approved by the Institutional Review Board at Boston University Medical Center (Boston, MA).

*DNA Methylation*

Peripheral blood samples were collected at the 8th examination. Genomic DNA was extracted from buffy coat using the Gentra Puregene DNA extraction kit (Qiagen) and bisulfite converted using EZ DNA Methylation kit (Zymo Research Corporation). DNA methylation quantification was conducted in two laboratory batches using the Illumina Infinium HumanMethylation450 array (Illumina). Methylation beta values were generated using the Bioconductor minfi package with Noob background correction [29].

*Genotyping, imputation and quality control*

FHS genotyping was carried out using Affymetrix 500k and MiPs 50K gene-centered chip.

Imputation was based on Minimac3 with HRC reference release 1. Prior imputation, quality control removed variants that satisfied any of the following criteria: Hardy-Weinberg equilibrium P < 1.0E-06, call rate less than 96.9%, minor allele frequency less than 0.01, number of Mendelian errors greater or equal to 1000, and variants at locations that did not map to GRCh37. Of the 546,344 genotyped SNPs, 412,049 variants remained after quality control. The genotyped and imputed markers datasets were downloaded from dbGAP (accession number: phs000724.v2.p9).

We performed GWAS in FHS offspring cohort (N=2,322) on both genotyped and imputed markers. A relaxed QC was applied to the imputed markers including the thresholds of MaCH Rsq at 0.2 and MAF at 0.005. A small number of monomorphic SNPs detected by Hardy-Weinberg analysis in PLINK, based on founders, were removed from GWAS analysis. GWAS was performed in fast MLM-based tool under GCTA (--fastGWA-lmm-exact), https://www.biorxiv.org/content/10.1101/598110v1. The random covariance matrix in the linear mixed analysis was estimated based on kinship coefficients of the pedigree structure. The kinship coefficients based genetic related covariance matrix was prepared using the R code provided by the GCTA tool. The fastGWA option adapts a sparse genetic relationship matrix that accounts up to the third relationship; the genetic variance component can be estimated from closely related individuals that captures both genetic variance and the amount of phenotypic variance attributable to shared environmental effect. The linear mixed analysis was also adjusted for sex and three principal components as fixed effects. GWAS of granulocytes was additionally adjusted for age as a fixed effect.

*Data availability*

The FHS data are available at dbGaP under the accession numbers phs000342 and phs000724.

*Acknowledgements*

The FHS data were accessed from dbGaP under the accession numbers phs000342 and phs000724 (https://www.ncbi.nlm.nih.gov/projects/gap/cgi-bin/study.cgi?study_id=phs000007.v31.p12). The Framingham Heart Study is conducted and supported by the National Heart, Lung, and Blood Institute (NHLBI) in collaboration with Boston University (Contract No. N01-HC-25195, HHSN268201500001I and 75N92019D00031). This manuscript was not prepared in collaboration with investigators of the Framingham Heart Study and does not necessarily reflect the opinions or views of the Framingham Heart Study, Boston University, or NHLBI. Funding for SHARe Affymetrix genotyping was provided by NHLBI Contract N02-HL64278. SHARe Illumina genotyping was provided under an agreement between Illumina and Boston University. DNA methylation data was funded from DIR research budget. National Institutes of Health, Bethesda, MD, USA, Daniel Levy PI. Horvath and Lu were supported by 1U01AG060908 – 01.

**Generation Scotland: Scottish Family Health Study (GS)**

*Cohort description*

Generation Scotland is a family-structured, population-based cohort study of over 24,000 people from across Scotland, aged between 18 and 99 years at the study baseline (2006-2011). A broad set of phenotype data were collected at baseline and data linkage to electronic health records has enabled follow-up to collect information on incident disease outcomes. Full details have been reported previously [36, 39, 40]. DNA was obtained from whole blood at the study baseline.

*Ethics*

All components of Generation Scotland received ethical approval from the NHS Tayside Committee on Medical Research Ethics (REC Reference Number: 05/S1401/89). All participants provided broad and enduring written informed consent for biomedical research. Generation Scotland has also been granted Research Tissue Bank status by the East of Scotland Research Ethics Service (REC Reference Number: 20/ES/0021), providing generic ethical approval for a wide range of uses within medical research. This study was performed in accordance with the Helsinki declaration.

*DNA Methylation*

DNA methylation from whole blood was assessed using the Illumina EPIC array. To date, the arrays have been run on two tranches of the GS data (Set 1 – n=5,100; Set 2 – n = 4,450). Full details on quality control have been reported previously. Briefly, probes were filtered based on three criteria: outliers based on visual inspection of the log median intensity of the methylated versus unmethylated signal per array; a beadcount <3 in more than 5% of samples; and more than 5% of samples have a detection p-value >0.05. Samples were removed if predicted sex did not match reported sex and if >1% of CpGs had a detection p-value >0.05. These data were then uploaded to Horvath’s online calculator, as described in the main text, to generate the various clock estimates.

*Genotyping, imputation and quality control*

GS genotyping was carried out over two batches with 9,863 samples genotyped using the Illumina HumanOmniExpressExome-8 v1.0 Bead Chip with the remainder genotyped using the Illumina HumanOmniExpressExome-8 v1.2 Bead Chip, with Infinium chemistry for both. Genotype quality control details have been described previously [41] and included the removal of individuals or SNPs with a low call rate (<98%), SNPs with a Hardy-Weinberg P-value<1x10^-6^. Ancestry outliers, defined as >6 SDs from the mean in a principal components analysis of GS merged with 1,092 individuals from the 1000 Genomes Project [42]. The QC’d SNP data were imputed to the HRC reference panel (v1.1). Further QC was carried out, as described in Nagy et al. [41], leaving an imputed dataset of over 24 million variants for analysis. The GWAS was run using RegScan v0.2 [43]. The effect size, standard error and p-values were corrected to account for relatedness using the GRAMMAR-Gamma factors from the polygenic function in GenABEL [44].

*Data availability*

According to the terms of consent for GS participants, access to individual-level data (omics and phenotypes) must be reviewed by the GS Access Committee. Applications should be made to [](https://genomemedicine.biomedcentral.com/articles/10.1186/s13073-019-0693-z). Guidance on the Generation Scotland Access Process and Policy can be found here: https://www.ed.ac.uk/generation-scotland/using-resources/access-to-resources

*Acknowledgements*

GS received core support from the Chief Scientist Office of the Scottish Government Health Directorates (CZD/16/6) and the Scottish Funding Council (HR03006). Genotyping and DNA methylation profiling of the GS samples was carried out by the Genetics Core Laboratory at the Clinical Research Facility, University of Edinburgh, Edinburgh, Scotland and was funded by the Medical Research Council UK and the Wellcome Trust (Wellcome Trust Strategic Award “STratifying Resilience and Depression Longitudinally” ([STRADL; Reference 104036/Z/14/Z]). CH is supported by an MRC University Unit Programme Grant MC_UU_00007/10 (QTL in Health and Disease).

**The Genetics of DNA Methylation Consortium (GoDMC) Meta-GWAS**

GoDMC was established with the view of bringing together researchers with an interest in studying the genetic basis of DNA methylation variation, to consolidate as many resources and expertise as possible and thereby expedite this field of research. The initial release of their findings consists of mQTL associations based on a sample size of 27,750 individuals. Summary statistics of these meta analyses are available on request from http://www.godmc.org.uk/projects.html.

*Genotype data*

Genotype data of all autosomes and chromosome X (if available) was imputed to 1000G and above using hg19/build37. Genotype data was filtered on an info score of 0.8 and a minor allele frequency (MAF) of 0.01. Genotype data was converted to bestguess data without a probability cut-off.

DNA methylation data: DNA methylation was measured in whole blood or cord blood using Illumina 450k or EPIC Beadchips in at least 100 European individuals. Normalized beta values were used, preferable normalized with the R package meffil [4]. Most analysts used meffil to quality control and normalize the DNA methylation data using functional normalization. Protocols can be found here: https://github.com/perishky/meffil/wiki.

A github pipeline was implemented to run the analyses locally. For the genotype data, several standard sample QC steps were performed including a sex check, removal of samples with >5% missingness, and the identification and exclusion of ethnic outliers. In datasets of ostensibly unrelated individuals, those that were found to be related (identity by state > 0.125) were excluded.

The pipeline then residualised the normalized methylation betas by replacing outliers that were 10 standard deviations from the mean (3 iterations) with the probe mean, rank transforming the normalized beta values and regressing out age, sex, predicted cell counts, predicted smoking, genetic principal components and non-genetic methylation principal components. In family-based cohorts, genetic relatedness matrices were constructed and relatedness adjusted for using the GRAMMAR approach. Genomic lambdas were checked by performing a GWAS of cg07959070. These residualised methylation measurements were used in all analyses.

*Association analysis*

First, every study performed a full analysis of all candidate mQTL associations, returning only associations at a threshold of p<1e-5. All candidate mQTL associations at p<1e-5 were combined to create a unique ‘candidate list’ of mQTL associations. In total, 102,965,711 candidate mQTL associations in cis (p<1e-5, SNP located within 1Mb of the methylation site) and 710,638,230 candidate mQTL associations in trans were identified in at least one dataset. To avoid computational burden, we included cis associations found in at least one dataset and trans associations in at least two datasets. The candidate list (n=120,212,413) was then sent back to all cohorts and the association estimates obtained for every mQTL association on the candidate list.

*Meta analyses*

Meta analyses were run using a modified version of METAL using 962 chunks. Candidate mQTL associations were meta-analysed using fixed effects, additive random effects and multiplicative random effects models. 36 datasets from European origin were included in the meta-analyses.

**GENOA African Americans**

*Cohort description*

The Genetic Epidemiology Network of Arteriopathy (GENOA) study, a part of the Family Blood Pressure Program [45], consists of hypertensive sibships that were recruited for linkage and association studies in order to identify genes that influence blood pressure and its target organ damage [46]. In the initial phase of the GENOA study (Phase I: 1996-2001), all members of sibships containing ≥ 2 individuals with essential hypertension clinically diagnosed before age 60 were invited to participate, including both hypertensive and normotensive siblings. In the second phase of the GENOA study (Phase II: 2000-2004), 1,239 non-Hispanic white and 1,482 African American participants were successfully re-recruited to measure potential target organ damage due to hypertension.

*Ethics*

The GENOA study was approved by Institutional Review Boards at the University of Michigan and the University of Mississippi Medical Center.

*DNA methylation*

A total of 1,106 samples at Phase I and 304 samples at Phase II were assessed using the Illumina HumanMethylationEPIC BeadChip. First, raw IDAT files were imported using Minfi R package [29]. We used the shinyMethyl R package [47] to visualize the raw intensity data and identify sex mismatches and outliers, which were removed. We also obtained detection p-value for each sample at each probe, and individual probes with detection p-value <10^-16^ were considered to be detected successfully [48]. Samples and probes with detection rate <10% were removed. Samples with incomplete bisulfite conversion identified using the QCinfo() function in the ENmix R package were removed [49]. We also checked sample identity using the 59 SNP probes implemented in the EPIC chip and removed mismatched samples. Next, Noob was used for individual background and dye-bias normalization [50]. Since two types of probes are present on the EPIC BeadChip (Infinium I and Infinium II), we used the Regression on Correlated Probes (RCP) method to adjust for probe-type bias [51]. ComBat, an empirical Bayes batch-correction method was used sequentially to remove those batch effects [52]. We performed principal variance component analysis (PVCA) to quantify the variance explained by the known batch variables before and after batch adjustment to ensure that no single batch factors explained more than 3% of the variance. After exclusions, a total of 857,121 probes in 1,100 samples at Phase I and 294 samples at Phase II were available for analysis.

A subset of CpG sites from EPIC data was used with the online Horvath epigenetic clock software found at <https://dnamage.genetics.ucla.edu/new> to calculate the DNAm estimates of the epigenetic measures. The measures extracted and used for analysis included Hannum age-acceleration (AgeAccelerationResidualHannum), Intrinsic Horvath age-acceleration (IEAA), PhenoAge-acceleration (AgeAccelPheno), GrimAge-acceleration (AgeAccelGrim), DNAmPAI1 adjusted for age (DNAmPAI1adjAge), and Granulocyte count (variable Gran). After removing outliers beyond 5 standard deviations from the mean, the final sample with non-missing clock and genotype data included N=962 participants for three of the measures and N=961 for the other three.

*Genotyping and quality control*

All GENOA participants were genotyped on the Affymetrix Genome-Wide Human SNP Array 6.0 or the Illumina Human 1M-Duo BeadChip. Samples were removed if they had a missing call rate ≥0.05 or were an outlier ≥6 standard deviations from the mean of the first 10 genome-wide principal components from genotype data. SNPs with a missing call rate ≥0.05 were removed. Imputation was done on the Michigan Imputation server (<https://imputationserver.sph.umich.edu/index.html#!pages/home>) using SHAPEIT and minimac3. Genotypes were imputed to the Haplotype Reference Consortium (HRC version r1.1) reference panel.

*Data availability*

According to the terms of consent for GENOA participants, access to epigenetic data will be made available by reasonable request and with a corresponding Data Use Agreement. Please contact Sharon L.R. Kardia and Jennifer A. Smith.

*Acknowledgements*

Support for the Genetic Epidemiology Network of Arteriopathy (GENOA) was provided by the National Heart, Lung and Blood Institute (HL054457, HL100185, HL119443, and HL133221). We would also like to thank the families that participated in the GENOA study.

**GOLDN**

The Genetics of Lipid Lowering Drugs and Diet Network Study (GOLDN) study determined genetic predictors of participant lipid and inflammatory response to diet and fenofibrate treatment interventions. Participants were identified through 3-generation families previously screened in the National Heart, Lung and Blood Institute Family Heart Study (FHS) Minnesota and Utah centers. DNA collected before the two interventions in a total of 993 GOLDN participants was assayed using Illumina’s Infinium HumanMethylation450 Beadchip as previously described by Irvin et al. Briefly, CD4+ T cells were harvested from stored buffy coats using antibody-linked Invitrogen Dynabeads. Cells were lysed and DNA was extracted using DNeasy kits (Qiagen, Venlo, Netherlands). After following standard Illumina protocols, the resulting intensity files were analyzed with Illumina’s GenomeStudio. After QC, methylation data were available for 461,281 5′-cytosine-phosphate-guanine-3′ genomic site (CpGs). The non-normalized data were uploaded into the DNA methylation age calculator using the advanced analysis in blood option. For the GWAS analysis of methylation age we used Whole Genome Sequencing data from GOLDN study. In summary, a total of 24,369,178 single nucleotide polymorphisms (SNPs) were genotyped on 965 individuals. After quality control exclusions, 7,770,568 SNPs remained

*Data availability*

The GOLDN study methylation data is available on dbGaP Study Accession: phs000741.v1.p1 or through direct data request with the PI.

*Acknowledgements*

GOLDN biospecimens, baseline phenotype data, and intervention phenotype data were collected with funding from National Heart, Lung and Blood Institute (NHLBI) grant U01 HL072524. Whole-genome sequencing in GOLDN was funded by NHLBI grant R01 HL104135-04S1.

**Grady Trauma Project (GTP)**

*Cohort description*

Grady Trauma Project (GTP) investigates the influence of genetic and environmental factors on response to stressful events in a predominantly African-American, urban population of low socioeconomic status. A broad set of phenotype data were collected from GTP research participants who were approached in the waiting rooms of the primary care clinic or obstetrical-gynecological clinic of a large, urban, public hospital in Atlanta, GA while either waiting for their medical appointments or while waiting with others who were scheduled for medical appointments [54]. Demographic variables including age, sex and race were assessed through self-report.

*Ethics*

All participants provided broad and enduring written informed consent for biomedical research. The Institutional Review Boards of Emory University School of Medicine and the Research Oversight Committee of Grady Memorial Hospital approved this study.

*DNA Methylation*

DNA was extracted from whole blood and interrogated using the MethylationEPIC BeadChip (Illumina) for 737 samples and HumanMethylation450 BeadChip (Illumina) for 91 samples according to manufacturer’s instructions. Raw methylation beta values were determined via GenomeStudio. Samples with probe detection call rates <90% and those with an average intensity value of either <50% of the experiment-wide sample mean or <2,000 arbitrary units (AU) were removed using R package CpGassoc [55]. Probes with detection p-values >0.01 were set to missing. CpG sites that cross-hybridize between autosomes and sex chromosomes were removed [56]. These data were then uploaded to Horvath’s online calculator, as described in the main text, to generate the various clock estimates. DNA methylation data for both arrays are publicly available on GEO database (GSE132203 for MethylationEPIC BeadChip and GSE72680 for HumanMethylation450 BeadChip).

*Genotyping, imputation and quality control*

Genotyping was performed using Illumina Omni-Quad 1M, and genotypes were called in Illumina’s GenomeStudio. Genotype quality control and imputation details have been described previously [57]. Briefly, samples with low call rates (<98%, using SNPs with call rates >95%), deviating from expected inbreeding coefficient (*f*_het_ < −0.2 or >0.2), or with a sex discrepancy between reported and estimated sex based on inbreeding coefficients calculated from SNPs on X chromosomes were removed. Similarly, monomorphic SNPs as well as SNPs with low call rates (<98%), Hardy-Weinberg P-value<1x10^-6^, or a > 2% difference in missing genotypes between cases and controls were excluded. Within-study relatedness was estimated using the IBS function in PLINK 1.9 [58]. From each pair with relatedness $\hat{\pi}$ > 0.2, one individual was removed from further analysis, retaining cases where possible. Imputation was based on the 1000 Genomes phase 3 data (1KGP phase 3) [8], using default settings in SHAPEIT2 v2.r837 [7] for pre-phasing and IMPUTE2 v2.2.2 [26] for imputation of haplotypes. To calculate principal components (PCs), SNPs that had a MAF <5%, HWE P > 0.05, were ambiguous (A/T, G/C), or located in the MHC region (chr. 6, 25–35 MB) or chromosome 8 inversion (chr. 8, 7–13 MB) were excluded. SNPs were pairwise LD-pruned (r^2^ > 0.2) and PCs for each subject were calculated using PLINK 1.9 [6]. The number of significant principal components was identified by a Tracy-Widom test. The GWAS was run using PLINK 1.9 [6], adjusting for first 6 PCs, sex and methylation array type.

*Data availability*

GTP DNA methylation data are available through the NCBI Gene Expression Omnibus (GEO) under accession number GSE132203 for MethylationEPIC BeadChip and GSE72680 for HumanMethylation450 BeadChip. Individual level genotype and phenotype data may be requested through the PGC Data Access Portal at https://www.med.unc.edu/pgc/shared-methods/ data-access-portal/ or from the Grady Trauma Project (http://gradytraumaproject.com/), upon review by GTP Access Committee. Applications should be made to Associate Program Director Rebecca Hinrichs at.

*Acknowledgements*

This study was supported by the National Institutes of Health, MH071537 and MH096764.

**INVECCHIARE NEL CHIANTI (InCHIANTI study)**

*Cohort description*

The InCHIANTI study is a population-based epidemiological study aimed at evaluating the factors that influence mobility in the older population living in the Chianti region in Tuscany, Italy. The details of the study have been previously reported [59]. Briefly, 1616 residents were selected from the population registry of Greve in Chianti (a rural area: 11,709 residents with 19.3% of the population greater than 65 years of age), and Bagno a Ripoli (Antella village near Florence; 4,704 inhabitants, with 20.3% greater than 65 years of age). The participation rate was 90% (n=1453), and the subjects ranged between 21-102 years of age. Overnight fasted blood samples were for genomic DNA extraction, and measurement of genetic variation and DNA methylation.

*Ethics*

The InCHIANTI study protocol was approved by the Italian National Institute of Research and Care of Aging Institutional Review and Internal Review Board of the National Institute for Environmental Health Sciences (NIEHS) and all participants provided written informed consent.

*DNA Methylation*

Genome-wide DNA methylation (DNAm) was assayed on DNA samples collected at the baseline visit with genetic data. Genomic DNA was bisulfite converted using Zymo EZ-96 DNA Methylation Kit (Zymo Research Corp., Irvine, CA) per the manufacturer's protocol. CpG methylation status of 485,577 CpG sites was determined using the Illumina Infinium HumanMethylation450 BeadChip (Illumina Inc., San Diego, CA) per the manufacturer's protocol [60]. Quality control procedures for this data has been previously described [61]. Briefly, probes were filtered if a beadcount <3 in more than 5% of samples; and more than 5% of samples have a detection p-value >0.05. Subjects were filtered call rate was <95%, ≥5% of CpGs had a detection p-value >0.01, and predicted sex did not match the reported sex. Background and quantile normalization were applied to the filtered dataset. The data that passed quality control were uploaded to Horvath’s online calculator as described in the main text, to generate the various clock estimates.

*Genotyping, imputation and quality control*

Genome-wide genotyping was completed using Illumina Infinium HumanHap 550K SNP arrays were used for genotyping [62]. Genotyping was completed for 1210 subjects with a sample call rate >97%, heterozygosity rates > 0.3 and correct sex specification. 495,343 autosomal SNPs that passed quality control (MAF>1%, completeness >99%, HWE > 10-4) were used for imputation. The QC’d SNP data were imputed to the HRC reference panel (v1.1) using the Michigan imputation server [20]. GWAS was run using PLINK 1.9 [6].

*Data availability*

The InCHIANTI data are available upon request. Applications should be made through the website: http://inchiantistudy.net/wp/

*Acknowledgements*

The InCHIANTI study baseline (1998-2000) was supported as a "targeted project" (ICS110.1/RF97.71) by the Italian Ministry of Health and in part by the U.S. National Institute on Aging (Contracts: 263 MD 9164 and 263 MD 821336). This work was supported in part by the Intramural Research Program of the National Institute on Aging, National Institutes of Health, Baltimore, Maryland. This work utilized the computational resources of the NIH HPC Biowulf cluster. (http://hpc.nih.gov).

**Jackson Heart Study (JHS)**

*Cohort description*

Between 2000 and 2004, JHS recruited 5,306 African American participants from the Jackson, Mississippi, metropolitan tri-county area (Hinds, Madison, and Rankin). JHS is a prospective, community-based cohort designed to investigate risk factors for cardiovascular disease among African Americans. A range of measures, including traditional and putative CVD risk factors, health behaviors, detailed demographic, socioeconomic and sociocultural factors, medication use, anthropometry, blood pressure, assessments of kidney function and diabetes, and biochemical analytes, were obtained at the baseline JHS examination and in two subsequent clinic visits (2005-2008 and 2009-2013). [63, 64, 65]

*Ethics*

All participants included in this analysis provided written, informed consent for use of genetic data, and all study protocols conform to the 1975 Declaration of Helsinki guidelines. The study was approved by the Institutional Review Boards of the participating institutions (University of Mississippi Medical Center, Jackson State University and Tougaloo College).

*DNA Methylation*

Illumina Methylation EPIC array data (containing over 850,000 CpG methylation sites) was generated using whole blood samples collected at the JHS baseline exam. Methylation β values (the ratio of intensities between methylated and un-methylated alleles) were normalized with respect to background color intensity using the normal-exponential out-of-band (NOOB) preprocessing method in the R package minfi [50]. Data on non-duplicate samples passing quality control were then uploaded to Horvath’s online calculator, as described in the main text, to generate the various clock estimates. We excluded values outside of 5 SD from the mean for the methylation age measures before performing GWAS analyses.

*Genotyping, imputation and quality control*

Affymetrix 6.0 GWAS genotyping was performed in n=3,029 JHS participants with consent for genetic analyses through NHLBI’s Candidate Gene Association Resource (CARe) consortium. [66] Genotyping and quality control have been previously described [67]. Imputation to 1000 Genomes Project (1000G) Phase 3 version 5 reference panel was completed using Minimac3 on the Michigan Imputation Server [20]. Prior to imputation, SNPs were filtered for minor allele frequency ≥1%, call rate ≥ 90%, HWE p-value > 10-6, as well as exclusion of sites with invalid or mismatched alleles for the reference panel. Samples with sex or pedigree mismatches and principal components outliers were excluded. Analysis was conducted using the EMMAX test as implemented in EPACTS 3.2.6.

*Data availability*

JHS data is available on dbGaP at phs000286, as well as by manuscript proposal request at https://www.jacksonheartstudy.org/.

*Acknowledgements*

The Jackson Heart Study (JHS) is supported and conducted in collaboration with Jackson State University (HHSN268201800013I), Tougaloo College (HHSN268201800014I), the Mississippi State Department of Health (HHSN268201800015I) and the University of Mississippi Medical Center (HHSN268201800010I, HHSN268201800011I and HHSN268201800012I) contracts from the National Heart, Lung, and Blood Institute (NHLBI) and the National Institute for Minority Health and Health Disparities (NIMHD). The authors also wish to thank the staffs and participants of the JHS.

The views expressed in this manuscript are those of the authors and do not necessarily represent the views of the National Heart, Lung, and Blood Institute; the National Institutes of Health; or the U.S. Department of Health and Human Services. The funders had no role in the design and conduct of the study, in the collection, analysis, and interpretation of the data, and in the preparation, review, or approval of the manuscript.

LMR was funded by T32 HL129982. JWG and APR were funded by R01 HL116446

**Cooperative Health Research in the Region of Augsburg (KORA)**

*Cohort description*

The KORA (Kooperative Gesundheitsforschung in der Region Augsburg) research platform has been collecting clinical and genetic data from individuals of German nationality from the general population living in the region of Augsburg in southern Germany for over 20 years. F4 (2006-2008) and FF4 (2013-2014) cohorts are follow-up studies from the KORA S4 (n=4,261) survey carried out 1999-2000. In the baseline examinations all inhabitants of German nationality between the ages of 25 and 74 years were enrolled. Participants completed a lifestyle questionnaire, including details on health status and medication use, underwent standardized examinations with blood samples taken; study design, sampling method and data collection have been described in detail elsewhere [68]. The present study is based on a subsample of 1,727 participants of KORA F4 and with methylation and genotyping data available.

*Ethics*

The KORA cohort ethical approval was granted by the ethics committee of the Bavarian Medical Association (REC reference numbers: F4: #06068) and all were carried out in accordance with the principles of the Declaration of Helsinki. All research participants have signed informed consent prior to taking part in any research activities. The KORA data protection procedures were approved by the responsible data protection officer of the Helmholtz Zentrum München.

*DNA Methylation*

Genome-wide DNA methylation measurement at 485,577 genomic sites for KORA F4 was performed using the Infinium HumanMethylation450K BeadChip® (Illumina, Inc., CA, USA) [69] in 1802 KORA F4 samples. Sample preparation and measurement have been described previously [70]. Methylated and unmethylated signal intensities were obtained for each CpG site, which were then converted to β-values, the ratio of the methylated signal intensity divided by the overall signal intensity [69, 71]. DNA methylation data were preprocessed following the CPACOR pipeline of Lehne et al. [48]. First, 65 probes that represent SNPs were excluded. Second, background correction was performed using the R package minfi, version 1.6.0 [29]. Third, detection p-values were defined as the probability of a signal being detected above the background signal level, as estimated from negative control probes. Consequently, signals with detection p-values ≥ 0.01 were removed, as they indicate putatively unreliable signals. Similarly, signals summarized from less than three functional beads on the chip were characterized as potentially unreliable and removed from the data set. Observations with less than 95% of CpG sites providing reliable signals (75 in KORA F4) were excluded, resulting in 1727 samples overall.

Data were normalized using quantile normalization on the raw signal intensities. Beta-mixture quantile normalization was performed on a stratification of the probe categories into 6 types, based on probe type and color channel, using the R package limma, version 3.16.5 [72]. These data were then uploaded to Horvath’s online calculator, as described in the main text, to generate the various clock estimates.

*Genotyping, imputation and quality control*

Genotyping and calling were performed using the Affymetrix Axiom platform and software. Samples and SNPs were subject to a 97% and 98% call rate threshold respectively. SNPs with minor allele frequency < 1% or with Hardy-Weinberg Equilibrium P > 5x10-6 were also removed. Imputation was performed using IMPUTE v2.3.0, reference panel 1000g phase1 integrated haplotypes (produced using SHAPEIT2) and GWASs were conducted using SNP Test v.2.5 [27]. No evidence of population stratification was found for multiple published analyses. [73, 74]

*Data Availability*

The informed consents given by KORA study participants do not cover data posting in public databases. However, data are available upon request from KORA Project Application Self-Service Tool (https://epi.helmholtz-muenchen.de/) Data requests can be submitted online and are subject to approval by the KORA Board.

*Acknowledgements*

The KORA study was initiated and financed by the Helmholtz Zentrum München – German Research Center for Environmental Health, which is funded by the German Federal Ministry of Education and Research (BMBF) and by the State of Bavaria. Furthermore, KORA research has been supported within the Munich Center of Health Sciences (MC-Health), Ludwig-Maximilians-Universität, as part of LMUinnovativ.

**Lothian Birth Cohorts of 1921 and 1936 (LBC1921 and LBC1936)**

*Cohort description*

The Lothian Birth Cohorts of 1921 and 1936 are two longitudinal studies of ageing. They include individuals born in 1921 and 1936, respectively, who completed an IQ test at age 11 as part of the Scottish Mental Surveys of 1932 and 1947. Survivors living in the Edinburgh area of Scotland were contacted in later life (from mean age 79 in LBC1921 and 70 in LBC1936) to join these prospective cohort studies. There have been multiple waves of data collection, approximately every 3 years, with a vast array of cognitive, personality, health, lifestyle and omics data generated. Full details have been reported previously [75, 76]. DNA for the current study was obtained from whole blood at the study baseline.

*Ethics*

Following informed consent, venesected whole blood was collected for DNA extraction in both LBC1921 and LBC1936. Ethics permission for the LBC1921 was obtained from the Lothian Research Ethics Committee (Wave 1: LREC/1998/4/183). Ethics permission for the LBC1936 was obtained from the Multi-Centre Research Ethics Committee for Scotland (Wave 1: MREC/01/0/56), the Lothian Research Ethics Committee (Wave 1: LREC/2003/2/29).

*DNA Methylation*

DNA methylation from whole blood was assessed using the Illumina HumanMethylation450BeadChips array. IDAT files (for individuals who passed previous QC thresholds as reported in Zhang et al. [77]) were processed using the minfi package in R, with Noob being used to normalise the data – preprocessesNoob() function in minfi – via background subtraction and dye-bias normalisation [29]. Normalised betas were obtained via the getBeta() function in minfi.

*Genotyping, imputation and quality control*

LBC DNA was genotyped on the Illumina 610-Quadv1 array by the Edinburgh Clinical Research Facility. Quality control steps have been reported [78]. Filtering steps included: sex discrepancies, removal of one person per related pair, individual call rate <95%, non-European ancestry, SNP call rate <98%, and Hardy-Weinberg equilibrium test P<0.001. Imputation was carried out to the 1000G reference panel (phase 3) [42] and GWASs were conducted in mach2qtl [79], adjusting for four genetic principal components.

*Data availability*

LBC data are available on request from the Lothian Birth Cohort Study, University of Edinburgh (Ian Deary,). LBC data are not publicly available due to them containing information that could compromise participant consent and confidentiality.

*Acknowledgements*

The authors thank all LBC study participants and research team members who have contributed, and continue to contribute, to ongoing LBC studies. The LBC1936 is supported by Age UK (Disconnected Mind program) and the Medical Research Council (MR/M01311/1).  The Lothian Birth Cohort 1921 was supported by the UK’s Biotechnology and Biological Sciences Research Council (BBSRC), The Royal Society and The Chief Scientist Office of the Scottish Government. Methylation typing was supported by Centre for Cognitive Ageing and Cognitive Epidemiology (Pilot Fund award), Age UK, The Wellcome Trust Institutional Strategic Support Fund, The University of Edinburgh, and The University of Queensland. IJD and SEH are supported by Age UK (Disconnected Mind programme). IJD is supported by the UKRI Medical Research Council grant, MR/R0245065/1, and by National Institute of Health R01 grant, 1R01AG054628-01A1. GD is supported by the University of Edinburgh School of Philosophy, Psychology and Language Sciences.

**The Melbourne Collaborative Cohort Study (MCCS)**

*Cohort description*

The MCCS is a prospective cohort study of 41,513 residents (17,044 men) of Melbourne, Australia. The majority (99%) were aged 40 to 69 years and 41% were men. Southern European migrants were oversampled to extend the range of lifestyle-related exposures [79]. Participants were recruited via the electoral rolls (registration to vote is compulsory for adults in Australia), advertisements, and community announcements in local media (e.g., television, radio, and newspapers). Participants were contacted at both baseline (between 1990 and 1994) and follow-up (between 2003 and 2007). Blood samples were taken at baseline and follow-up from 99% and 64% of participants, respectively. Baseline samples were stored as dried blood spots on Guthrie cards for the majority (73%), as mononuclear cell samples for 25% and as buffy coat samples for 2% of the participants. Follow-up samples were stored as buffy coat samples and dried blood spots on Guthrie cards.

*Ethics*

All participants provided written informed consent and the study protocols were approved by the Cancer Council Victoria’s Human Research Ethics Committee.

*DNA Methylation*

The present study sample comprised MCCS participants selected for inclusion in one of seven previously conducted nested-case control studies of DNA methylation of colorectal, gastric, kidney, lung, and prostate cancer, B-cell lymphoma, and urothelial cell carcinoma (UCC) [80, 81, 82, 83, 84, 85]. All participants were cancer-free at blood draw. Controls were matched to incident cases of prostate, colorectal, gastric, lung or kidney cancer, UCC or mature B-cell neoplasms on sex, year of birth, country of birth, baseline sample type and smoking status (the latter for the lung cancer study only). In the UCC case-control study, 303 participants had their blood sample collected at follow-up (2004–2007); we excluded these participants from our cross-sectional analyses because their questionnaire data and storage time were different from those with blood collected at baseline. Only MCCS participants selected as controls were included in the present study. Methylation data for baseline blood samples (baseline study) were available from a total of 2,511 controls after quality control and exclusions. The Illumina Infinium HumanMethylation450K BeadChip (HM450K) array (Illumina, Inc.; San Diego, CA, USA) was used to measure DNA methylation as per the Illumina protocol [8]. To minimize the potential batch effects by nested case-control study cohort heterogeneity, the Illumina IDAT files were processed separately by study using R package “minfi”, and both Illumina’s background correction and control normalization were applied based on internal control probes. The result β values, which correspond to the percentage of methylation, and the corresponding detection P values were obtained via functions of package “minfi”, getBeta() and detectionP() [29]. For each sample, each CpG with a detection P value greater than 0.01 was assigned as missing. Samples with missing values for more than 5% of probes were excluded. CpG sites were excluded if they were missing for more than 20% of samples. These data were then reformatted and uploaded to the online calculator following the protocols established for the present study.

*Genotyping, imputation and quality control*

Genome-wide genotyping was been conducted on germline DNA samples from a subset of MCCS participants using the Illumina Infinium OncoArray-500K BeadChip [79]. These included prevalent and incident cases of breast, prostate and colorectal cancer, and incident cases of urothelial cell carcinoma, as well as controls matched to a subset of these using incidence density sampling. A total of 13,827 MCCS participants in these studies had available genetic data and 2,044 of those also were included in the previously mentioned methylation dataset.

Missing autosomal variants were imputed using the Michigan imputation server with the 1000 genome reference panel (phase 3) [20, 42] following standardised protocols [86]. Principal components were calculated based on the genotype data, which was first pruned based on pairwise linkage disequilibrium, using PLINK (Ver. 1.9).

*GWAS*

The GWAS were performed using the software R (version 3.4.4 on Ubuntu 14.04.6 LTS) with adjustments of top 10 principal components (continuous) and sex.

*Data availability*

The MCCS governance committee has determined that MCCS data are not to be made publicly available, but requests for access can be made to Cancer Council Victoria (https://www.cancervic.org.au/research/epidemiology/pedigree).

*Acknowledgements*

This work was supported by the Australian National Health and Medical Research Council (NHMRC) [grant 1088405]. MCCS cohort recruitment was funded by VicHealth and Cancer Council Victoria. The MCCS was further supported by Australian NHMRC grants 209057, 251553 and 504711 and by infrastructure provided by Cancer Council Victoria. Cases were ascertained through the Victorian Cancer Registry (VCR) and the Australian Cancer Database (Australian Institute of Health and Welfare). The nested case-control methylation studies were supported by the NHMRC grants 1011618, 1026892, 1027505, 1050198, 1043616 and 1074383.

**The Multi-Ethnic Study of Atherosclerosis (MESA) Study**

*Cohort description*

The Multi-Ethnic Study of Atherosclerosis (MESA) is a study of the characteristics of subclinical cardiovascular disease and the risk factors that predict progression to clinically overt cardiovascular disease or progression of the subclinical disease. MESA consisted of a diverse, population-based sample of an initial 6,814 asymptomatic men and women aged 45-84. 38 percent of the recruited participants were white, 28 percent African American, 22 percent Hispanic, and 12 percent Asian, predominantly of Chinese descent. Participants were recruited from six field centers across the United States: Wake Forest University, Columbia University, Johns Hopkins University, University of Minnesota, Northwestern University and University of California - Los Angeles. This study was approved by the IRB of each study site, and written informed consent was obtained from all participants. Each participant received an extensive physical exam and determination of coronary calcification, ventricular mass and function, flow-mediated endothelial vasodilation, carotid intimal-medial wall thickness and presence of echogenic lucencies in the carotid artery, lower extremity vascular insufficiency, arterial wave forms, electrocardiographic (ECG) measures, standard coronary risk factors, sociodemographic factors, lifestyle factors, and psychosocial factors. Selected repetition of subclinical disease measures and risk factors at follow-up visits allowed study of the progression of disease. Participants are being followed for identification and characterization of cardiovascular disease events, including acute myocardial infarction and other forms of coronary heart disease (CHD), stroke, and congestive heart failure; for cardiovascular disease interventions; and for mortality. The first examination took place over two years, from July 2000 - July 2002. It was followed by four examination periods that were 17-20 months in length, and a sixth exam is currently taking place.

*DNA Methylation*

Microarray technology was used to assay methylation at CpG sites in genomic DNA samples. The assays were performed using the Illumina EPIC array, which targets over 850,000 CpG sites. DNA samples were treated with bisulfite to measure 5-methylcytosine (5-mC) and 5-hydroxymethylcytosine (5-hmC): ~200 of these matched samples were also treated with an oxidizing reagent to distinguish between 5-mC and 5-hmC. Beta values will be corrected for background and normalized by standard procedures (e.g. BMIQ, SWAN, DASEN). Quality control procedures will include removing sites with detection p-values P>0.05 in more than 5-20% of samples and removing samples with detection p-values P>0.05 in > 5% of sites.

*Genotyping, imputation and quality control*

MESA European Americans, Hispanics, and Chinese were typed on the Affymetrix 6.0 SNP array at Affymetrix Research Services Lab. An additional 1738 African American samples were genotyped at Broad as part of the CARe project. Affymetrix performed plate-based genotype calling using Birdseed v2. Sample QC was based on call rates and contrast QC (cQC) statistics. Broad performed similar QC for CARe samples. Additional sample and SNP QC was carried out locally, including sample call rate, sample cQC, and sample heterozygosity by ethnicity at the sample level as well as outlier plate checking by call rate, median cQC or heterozygosity at plate level. Cryptic sample duplicates based on IBD/IBS were dropped. We excluded monomorphic SNPs across all samples; SNPs with missing rate > 5% or observed heterozygosity > 53% were also excluded. SNPs that were not directly genotyped were imputed to the 1000 Genomes Phase I integrated variant set (BCBI build 37 / hg19) based on data freeze from November 23, 2010 and May 21, 2011, using the IMPUTE version 2.2.2 program.

*Data availability*

Individual level genotype, phenotype, and methylation data are available on dbGaP (https://www.ncbi.nlm.nih.gov/projects/gap/cgi-bin/study.cgi?study_id=phs000209.v13.p3).

*Acknowledgements*

MESA and the MESA SHARe project are conducted and supported by the National Heart, Lung, and Blood Institute (NHLBI) in collaboration with MESA investigators. Support for MESA is provided by contracts 75N92020D00001, HHSN268201500003I, N01-HC-95159, 75N92020D00005, N01-HC-95160, 75N92020D00002, N01-HC-95161, 75N92020D00003, N01-HC-95162, 75N92020D00006, N01-HC-95163, 75N92020D00004, N01-HC-95164, 75N92020D00007, N01-HC-95165, N01-HC-95166, N01-HC-95167, N01-HC-95168, N01-HC-95169, UL1-TR-000040, UL1-TR-001079, UL1-TR-001420, UL1-TR-001881, and DK063491. MESA and the MESA SHARe project are conducted and supported by the National Heart, Lung, and Blood Institute (NHLBI) in collaboration with MESA investigators. Support for MESA is provided by contracts 75N92020D00001, HHSN268201500003I, N01-HC-95159, 75N92020D00005, N01-HC-95160, 75N92020D00002, N01-HC-95161, 75N92020D00003, N01-HC-95162, 75N92020D00006, N01-HC-95163, 75N92020D00004, N01-HC-95164, 75N92020D00007, N01-HC-95165, N01-HC-95166, N01-HC-95167, N01-HC-95168, N01-HC-95169, UL1-TR-000040, UL1-TR-001079, UL1-TR-001420, UL1-TR-001881, and DK063491

**National Institute on Alcohol Abuse and Alcoholism (NIAAA) cohort**

*Cohort description*

The sample consisted of 119 European American (EA) and 82 African American (AA) healthy controls that were recruited at the National Institute on Alcohol Abuse and Alcoholism (NIAAA) at the National Institutes of Health (NIH), USA. All participants completed the Structured Clinical Interview for Diagnostic and Statistical Manual of Mental Disorders (DSM)-IV-TR (SCID-IV) and were included if they did not meet criteria for alcohol dependence or other major psychiatric disorders and were considered healthy controls. The participants provided a blood sample which was used for genome-wide DNA methylation analysis and clinical marker collection. Full details have been published previously [82].

*Ethics*

All participants provided written informed consent in accordance with the Declaration of Helsinki and were compensated for their time. The study was approved by the Institutional Review Board of the NIAAA.

*DNA Methylation*

DNA methylation levels from whole blood samples were assessed using Illumina Infinium Methylation EPIC BeadChip microarray (Illumina Inc., San Diego, California) according to the manufacturer’s protocol. The wateRmelon package in R was used to process the raw data. After cross-reactive probes and probes that failed quality assessment were removed, a scale-based correction was applied for Illumina type I relative to type II probes. We used a quantile normalization approach to make methylated and unmethylated intensity values identical and then quantified the *β*-value using the ratio of intensities between methylated and unmethylated alleles [88]. The dataset consisted of *β*-values for 835,928 CpG sites. Full details on quality control have been reported previously [82].

*Genotyping, imputation and quality control*

All EA and AA participants were genotyped with the Illumina OmniExpress and Illumina OmniExpressExome BeadChips (Illumina, San Diego, CA). We performed population stratification analysis with the NIAAA participants and 2,504 individuals from the 1000 Genomes Project Phase 3 as a reference sample and identified and confirmed the ancestry information of each participant using EIGENSTRAT [89]. Based on the genetically confirmed group of EA and AA participants, we conducted a series of quality control (QC) procedures within each race group including sex discrepancies, Hardy-Weinberg equilibrium (HWE) test in EA and AA participants separately (P > 1.0E-5), SNP call rate > 99%, individual missing rate < 3%, and minor allele frequency (MAF) over 1%. The final sample size for the GWAS analyses after QC procedure was 103 EA and 75 AA individuals. The total number of SNPs tested in the GWAS was 613,209 for EA and 666,531 for AA. Furthermore, Imputation with Minimac3 available Michigan Imputation Server was performed using the 1000 Genomes reference panel (phase 3 v.5) in EA and AA separately [20]. The total number of imputed SNPs were 11,947,494 for EA and 20,178,596 for AA participants.

*Data availability*

The methylation data is not yet publicly available, but requests will be considered and can be submitted to Falk W. Lohoff.

*Acknowledgements*

This work was supported by the National Institutes of Health (NIH) intramural funding [ZIA-AA000242 to F.W.L]; Division of Intramural Clinical and Biological Research of the National Institute on Alcohol Abuse and Alcoholism (NIAAA).

**Netherlands Twin Register (NTR)**

*Cohort description*

The Netherlands Twin Register is a population-based cohort of over 200,000 people from across the Netherlands. It consists of twin-families, i.e. twins, their parents, spouses and siblings aged between 0 and 99 years at recruitment, and started around 1,987 with new–born twins and adolescent and adult twins. Full details have been reported previously [90]. DNA was collected from buccal cells and whole blood as part of multiple projects. For the current paper, we analysed DNA methylation measured in whole blood collected in the NTR-Biobank study [91, 92, 93]. Good quality whole blood DNA methylation data and genome-wide SNP data were available for 2,920 individuals, including monozygotic and dizygotic twins, parents of twins, siblings of twins and spouses of twins.

*Ethics*

Informed consent was obtained from all participants. The study was approved by the Central Ethics Committee on Research Involving Human Subjects of the VU University Medical Centre, Amsterdam, an Institutional Review Board certified by the U.S. Office of Human Research Protections (IRB number IRB00002991 under Federal-wide Assurance- FWA00017598; IRB/institute codes, NTR 03-180).

*DNA Methylation*

DNA methylation was assessed with the Infinium HumanMethylation450 BeadChip Kit (Illumina, San Diego, CA, USA) by the Human Genotyping facility (HugeF) of ErasmusMC, the Netherlands (http://www.glimdna.org/) as part of the Biobank-based Integrative Omics Study (BIOS) consortium [94]. DNA methylation measurements have been described previously [93, 94]. Genomic DNA (500ng) from whole blood was bisulfite treated using the Zymo EZ DNA Methylation kit (Zymo Research Corp, Irvine, CA, USA), and 4 µl of bisulfite-converted DNA was measured on the Illumina 450k array following the manufacturer’s protocol. A number of sample- and probe-level quality checks and sample identity checks were performed, as described in detail previously [93]. In short, sample-level QC was performed using MethylAid [95]. Probes were set to missing in a sample if they had an intensity value of exactly zero, or a detection p>.01, or a bead count of<3. After these steps, probes that failed based on the above criteria in >5% of the samples were excluded from all samples (only probes with a success rate≥ 0.95 were retained). The methylation data were normalized with functional normalization [5].

*Genotyping, imputation and quality control*

Genotyping in NTR was carried out in multiple batches on the Affymetrix 6.0, Affymetrix Perlegen, Illumina Human660, Illumina Omni 1M platforms. Initially samples were removed that (1) had abnormal heterozygosity (inbreeding F value <-0.075, or >0.075), or (2) had a Mendelian error rate > 5 standard deviations from the mean. SNPs were removed based on (1) minor allele frequency (<0.005), Hardy-Weinberg Equilibrium (HWE, *p* < 1*10^-12^), and call rate (<95%). Datasets from all platforms were merged and imputed to the Genome of the Netherlands (GONL, [96]) using Mac-admix [97, 98]. The resulting dataset were converted to best-guess genotypes using Plink 1.90, and filtered based on (1) a significant association with genotyping platform (*p* < 1*10^-5^), (2) a large mismatch between allele frequency and allele frequency in the GONL reference set (>10%), (3) HWE (*p* < 1*10^-5^), (4) a Mendelian error rate over 5 SD from the mean (*N*_errors_ > 40), (5) imputation quality (*R^2^* < 0.90). After this second phase the sample was imputed with the Haplotype Reference Consortium (HRC 1.1). The GWAS was performed with a mixed linear model association analysis in GCTA [99], using sex, genotyping platform and the first 10 principal components as fixed effect covariates and two genetic relatedness matrices (GRMs, [100]), which capture close and distant genetic relationships between individuals, respectively, as random effects.

*Data availability*

The HumanMethylation450 BeadChip data from the NTR are available as part of the Biobank-based Integrative Omics Studies (BIOS) Consortium in the European Genome-phenome Archive (EGA), under the accession code EGAD00010000887.

*Acknowledgements*

NTR warmly thanks all participants. We acknowledge funding from the Netherlands Organization for Scientific Research (NWO): Biobanking and Biomolecular Research Infrastructure (BBMRI–NL, 184.021.007; 184.033.111); epigenetic data were generated at HUMAN GENOMICS FACILITY (HUGE-F) at EUR.

Genotyping was made possible by grants from NWO/SPI 56-464-14192, Genetic Association Information Network (GAIN) of the Foundation for the National Institutes of Health, Rutgers University Cell and DNA Repository (NIMH U24 MH068457-06), the Avera Institute, Sioux Falls (USA) and the National Institutes of Health (NIH R01 HD042157-01A1, MH081802, Grand Opportunity grants 1RC2 MH089951 and 1RC2 MH089995) and European Research Council (ERC-230374). JvD is supported by NWO Large Scale infrastructures, X-Omics (184.034.019). DIB acknowledges the Royal Netherlands Academy of Science Professor Award (PAH/6635). MvdZ is supported by a grant from the Avera Institute for Human Genetics.

**Rotterdam Study 2 & Rotterdam Study 3 (RS2 &RS3)**

*Cohort description*

The Rotterdam Study is a large prospective, population-based cohort study aimed at assessing the occurrence of and risk factors for chronic diseases I the elderly. The study comprises 14,926 subjects in total, living in the well-defined Ommoord district in the city of Rotterdam in the Netherlands. In 1989, the first cohort, Rotterdam Study-I (RS-I) was comprised of 7983 subjects of age 55 years and above. IN 2000, the second cohort, Rotterdam Study-II (RS-II) was included with 3,011 subject who had reached an age of 55 or over in 2000. In 2006, the third cohort, Rotterdam Study-III (RS-III) was further included with 3932 subjects with age 45 years and over. Each participant gave written informed consent and the study was approved by the medical ethics committee of the Erasmus University Medical Center, Rotterdam.

*Methylation*

At the genetic Laboratory (department of Internal Medicine, ErasmusMC, Rotterdam, the Netherlands), the DNA methylation set was generated for a subset of 747 individuals of RS-III at baseline visit, as well as another subset of in total 864 individuals comprising of individuals at their fifth, third and second visit of RS-I, RS-II and RS-III respectively, who visited the research center between 2009 and 2013. Genomic DNA was extracted from whole peripheral blood by standard salting out methods. This was followed by bisulphite conversion using the Zymo EZ-96 DNA-methylation kit (Zymo Research, Irving, CA, USA). For each sample, whole genome amplification, fragmentation and hybridisation to the Infinium Illumina Human Methylation 450K arrays was performed according to the manufacturer’s protocol. Quality control of the samples was performed using MethylAid [95]. Probes with either a high detection p value (> 0.01), low bead count (< 3 beads), or low success rate (missing in > 5% of the samples) were set to missing. Samples were excluded from the analysis if they contained an excess of missing probes (> 5%). These data were then uploaded to Horvath’s online calculator, as described in the main text, to generate the various clock estimates.

*Genotyping, imputation and quality control*

Genotyping of the samples in RS-I and RS-II was carried out with the Illumina HumanHap 550v3 Genotyping BeadChip, for RS-III the Illumina HumanHap610-Quad V1 was used. The Beadstudio GenCall algorithm was used for genotype calling and quality control procedures. The following quality control inclusion filters were applied: call rate ≥97.5%, MAF≥1%, P for Hardy-Weinberg equilibrium <1x10-6. The total number of genotyped SNPs that passed these filters was; RS-I=512,349, RS-II=466,389 and RS-III=517,658. Imputation was done against the 1000G V.3 reference panel using the maximum likelihood method implemented in MACH (http://www.sph.umich.edu/csg/abecasis/MACH/download/1000G.2012-03-14.html). Analysis of imputed genotype data accounted for uncertainty in each genotype prediction by using the dosage information from MACH. GWAS was performed using rvtest software in RS-II and RS-III only as there were not enough samples with both genotype and methylation data in RS-I (n = 41).

*Data availability*

The informed consents given by Rotterdam study participants do not cover data posting in public databases. However, requests for the data accession may be sent to: Frank van Rooij.

*Acknowledgements*

The methylation data was funded by the Genetic Laboratory of the Department of Internal Medicine, Erasmus MC, and by a Rainbow Project (RP3; BIOS) of the Biobanking and Biomolecular Research Infrastructure Netherlands (BBMRI-NL), financed by the Netherlands Organisation forscientific research (NWO project 184.021.007). We thank Mr. Michael Verbiest, Ms. Mila Jhamai, Ms. Sarah Higgins, Mr. Marijn Verkerk, and Lisette Stolk PhD for their help in creating the methylation database.

The Rotterdam Study is funded by Erasmus Medical Center and Erasmus University, Rotterdam, Netherlands Organization for the Health Research and Development (ZonMw), the Research Institute for Diseases in the Elderly (RIDE), the Ministry of Education, Culture and Science, the Ministry for Health, Welfare and Sports, the European Commission (DG XII), and the Municipality of Rotterdam. The authors are grateful to the study participants, the staff from the Rotterdam Study and the participating general practitioners and pharmacists.

**The Southall And Brent REvisited Study (SABRE)**

*Cohort description*

The Southall And Brent REvisited Study (SABRE) is a population based cohort including 1710 first generation South Asian migrants, 801 first generation African Caribbean migrants, and 2346 people of European origin aged 40 to 69 living in West London, UK [101]. Baseline investigations were performed between 1988 and 1991. Peripheral blood samples were collected at baseline visits from participants recruited in the Southall district.

*Ethics*

The study was approved by St Mary's Hospital Research Ethics Committee (07/H0712/109) and all participants provided written informed consent. The study adheres to the principles of the Declaration of Helsinki and Title 45, US Code of Federal Regulations, Part 46, Protection of Human Subjects, Revised November 13, 2001, effective December 13, 2001.

*DNA Methylation*

Following DNA extraction, genomic DNA (500 ng) was bisulfite modified using an EZ DNA methylation kit (Zymo Research, Orange, CA, USA). The protocol was as described by the manufacturer, utilising the alternative incubation conditions recommended when using Illumina Infinium Methylation Arrays. Genome-wide DNAm was measured using the Illumina HumanMethylation450 BeadChip (Illumina, San Diego, CA, USA) following the manufacturer’s protocol with no modifications. The arrays were scanned using an Illumina iScan with software version 3.3.28.

Sample QC and normalisation was performed using *meffil* [4] in R version 3.1.1. Briefly, DNAm quality was evaluated by: sex detection outliers (N=13), the median intensity methylated vs unmethylated signal (N=10), dyebias (N=21), DNAm detection pvalue (N=588), and low bead numbers (N=167). Finally, 1993 samples passed QC. SABRE was normalized using 15 control probe PCs derived from the technical probes informed by *meffil* scree plots.

After filtering for European ancestry, those with DNAm measured at baseline, and those with genetic data available there were 731 individuals and 484,781 DNAm probes available for analysis.

*Genotyping, imputation and quality control*

2980 SABRE participants from the Southall centres were genotyped using a new UCL druggable target array. The array has a GWAS backbone comprising the Illumina Human Core BeadChip (~240k genome wide markers) and an additional custom set of 200k markers on genes encoding proteins involved in drug handling, drug action, and druggable targets. This was developed in collaboration with LSHTM and EBI.

Individuals were excluded due to incorrect sex assignment, high missingness (>5%), abnormal heterozygosity (het > mean (het) + 3*sd (het) or het < mean (het) - 3*sd (het)), cryptic relatedness (pihat>=0.9999 identified 20 cases of sample mislabelling (excluded), Pihat = 0.5 → 1^st^ degree relative. Pihat = 0.25 → 2^nd^ degree relative. – these were known relatives and remain in the dataset), and non-European ancestry (detected via PCA). Phasing was performed using Eagle and ethnicity-specific imputation was performed using Minimac3 [102] against the HRC reference panel (<http://www.haplotype-reference-consortium.org/>) [8]. Genotypes were filtered to have a MAF >0.005, imputation info score >0.3 and HWE p<0.00001. After data cleaning, QC, filtering for European ancestry, and sample matching to DNAm data availability, 731 individuals and 6,912,559 SNPs remained. Association tests were adjusted for 20 genetic PCs using SNPTEST v2.5.2. We used genotype dosages and applied additive models.

*Data availability*

Individual level genotype and DNA methylation data from SABRE are not permitted to be shared or deposited due to the original consent given at the time of data collection. However, genotype, DNA methylation, and phenotype data can be accessed through application to the LHA Data sharing committee

*Acknowledgements*

450K DNAm array data was generated in the Bristol Bioresource Laboratory Illumina Facility, University of Bristol.

*Funding*

The SABRE study was funded at baseline by the Medical Research Council, Diabetes UK, and the British Heart Foundation. DNAm analysis in the SABRE cohort was supported by a Wellcome Trust Enhancement grant 082464/Z/07/C. Genotyping analysis in the SABRE cohort was supported by the British Heart Foundation (CS/13/1/30327).

**The Swiss Cohort Study on Air Pollution and Lung and Heart Diseases in Adults (SAPALDIA)**

*Cohort description*

SAPALDIA is an on-going population based study initiated in 1991 to study air pollution impact on respiratory health. The design and protocol of SAPALDIA have been described in detail previously [103, 104]. At baseline (SAPALDIA1), 9,651 participants of age 18-60 years were recruited from eight communities representing various meteorological and geographical characteristics as well as different levels of urbanization. Participants underwent health examination and interviews. 8,047 and 6,088 out of the 9,651 participants were followed up in the second (SAPALDIA2 in 2001-3) and the third (SAPALDIA3 in 2010-11) survey, respectively. Peripheral blood samples were collected, processed, and archived at -80℃ in biobank at SAPALDIA2 and SAPALDIA3.

*Ethics*

SAPALDIA is in agreement with the declaration of Helsinki. All participants provided written informed consent and ethical approval was obtained from the Swiss Academy of Medical Sciences and the regional committees for each study center and for each survey.

*DNA Methylation*

DNA methylation was measured using Illumina Infinium 450k BeadChip array from 1971 peripheral blood samples collected at SAPALDIA2 and SAPALDIA3. Preprocessing and quality control have been described previously [105, 106]. In brief, raw fluorescence intensities were retrieved and processed using R package “minfi” [29], where Noob background correction and dye-bias correction were applied. Intensities with detection p-value > 10^-16^ were set missing. Probes with call rate < 0.95 or on sex chromosomes were excluded. Samples with call rate < 0.95 or sex mismatch were excluded. Beta-mixture quantile normalization (BMIQ) procedure was applied to correct for the Illumina probe design bias [14]. The epigenetic clock variables were then computed from Horvath’s online calculator. The epigenetic variables from 930 participants with DNA methylation as well as genotype data available were used in this study. In case of participants DNA methylation available for both time points (n=923), the epigenetic variables were averaged.

*Genotyping, imputation and quality control*

Genotyping was conducted on archived EDTA blood samples from two non-overlapping subsets of SAPALDIA2 participants using different arrays: 1,612 samples using Human610-Quad BeadChip (Illumina, San Diego, CA, USA) and 3,015 samples using Infinium Human OmniExpressExome-8 (Illumina, San Diego, CA, USA). Samples with call rate < 0.97 or population outliers were excluded. SNPs with call rate < 0.95, minor allele frequency < 0.05, or Hardy-Weinberg p < 10^-6^ were excluded. Each of the two genotype datasets was phased using ShapeIT (v2.r790) [7] and imputed using MiniMac2 (version 2014) [107] to 1000 Genome phase 1 reference panel. The two imputed datasets were finally merged to yield 38 million markers.

*Data availability*

Raw methylation and genetic data used in this study are not publically available because the informed consents provided by the study participants do not allow it. However, access can be granted upon reasonable request.

*Acknowledgements*

SAPALDIA was supported by the Swiss National Science Foundation, SNF-SAPALDIA (grants numbers 33CS30-148470/1&2, 33CSCO-134276/1, 33CSCO-108796, 324730_135673, 3247BO-104283, 3247BO-104288, 3247BO-104284, 3247-065896, 3100-059302, 3200-052720, 3200-042532, 4026-028099, PMPDP3_129021/1, PMPDP3_141671/1); the Swiss Federal Office for the Environment, the Swiss Federal Office of Public Health, the Swiss Federal Office of Roads and Transport, the canton's government of Aargau, Basel-Stadt, Basel-Land, Geneva, Luzern, Ticino, Valais, and Zürich, the Swiss Lung League, the canton's Lung League of Basel Stadt/ Basel Landschaft, Geneva, Ticino, Valais, Graubünden and Zurich, Stiftung ehemals Bündner Heilstätten, SUVA, Freiwillige Akademische Gesellschaft, UBS Wealth Foundation, Talecris Biotherapeutics GmbH, Abbott Diagnostics, European Commission 018996 (GABRIEL), Wellcome Trust WT 084703MA, EC FP7 grant (EXPOsOMICS; grant number 308610), EC Horizon 2020 research and innovation programme (ALEC study; grant number 633212).

The study could not have been done without the help of the study participants, technical and administrative support and the medical teams and field workers at the local study sites.

Study directorate: NM Probst-Hensch (PI; e/g); T Rochat (p), C Schindler (s), N Künzli (e/exp), JM Gaspoz (c)

Scientific team: JC Barthélémy (c), W Berger (g), R Bettschart (p), A Bircher (a), C Brombach (n), PO Bridevaux (p), L Burdet (p), Felber Dietrich D (e), M Frey (p), U Frey (pd), MW Gerbase (p), D Gold (e), E de Groot (c), W Karrer (p), F Kronenberg (g), B Martin (pa), A Mehta (e), D Miedinger (o), M Pons (p), F Roche (c), T Rothe (p), P Schmid-Grendelmeyer (a), D Stolz (p), A Schmidt-Trucksäss (pa), J Schwartz (e), A Turk (p), A von Eckardstein (cc), E Zemp Stutz (e).

Scientific team at coordinating centers: M Adam (e), I Aguilera (exp), S Brunner (s), D Carballo (c), S Caviezel (pa), I Curjuric (e), A Di Pascale (s), J Dratva (e), R Ducret (s), E Dupuis Lozeron (s), M Eeftens (exp), I Eze (e), E Fischer (g), M Foraster (e), M Germond (s), L Grize (s), S Hansen (e), A Hensel (s), M Imboden (g), A Ineichen (exp), A Jeong (g), D Keidel (s), A Kumar (g), N Maire (s), A Mehta (e), R Meier (exp), E Schaffner (s), T Schikowski (e), M Tsai (exp)

(a) allergology, (c) cardiology, (cc) clinical chemistry, (e) epidemiology, (exp) exposure, (g) genetic and molecular biology, (m) meteorology, (n) nutrition, (o) occupational health, (p) pneumology, (pa) physical activity, (pd) pediatrics, (s) statistics

Local fieldworkers SAPALDIA2: Aarau: M Broglie, M Bünter, D Gashi. Basel: R Armbruster, T Damm, U Egermann, M Gut, L Maier, A Vögelin, L Walter, Davos: D Jud, N Lutz. Geneva: M Ares, M Bennour, B Galobardes, E Namer. Lugano: B Baumberger, S Boccia Soldati, E Gehrig-Van Essen, S Ronchetto, Montana: C Bonvin, C Burrus. Payerne: S Blanc, AV Ebinger, ML Fragnière, J Jordan. Wald: R Gimmi, N Kourkoulos, U Schafroth.

Local fieldworkers SAPALDIA3: Aarau: S Brun, G Giger, M Sperisen, M Stahel. Basel: C Bürli, C Dahler, N Oertli, I Harreh, F Karrer, G Novicic, N Wyttenbacher. Davos: A Saner, P Senn, R Winzeler, Geneva: F Bonfils, B Blicharz, C Landolt, J Rochat. Lugano: S Boccia, E Gehrig, MT Mandia, G Solari, B Viscardi. Montana: AP Bieri, C Darioly, M Maire., Payerne: F Ding, P Danieli A Vonnez. Wald: D Bodmer, E Hochstrasser, R Kunz, C Meier, J Rakic, U Schafroth, A Walder.

Administrative staff: C Gabriel, R Gutknecht, N Bauer Ott

**The Swedish Adoption/Twin Study of Aging (SATSA)**

*Cohort description*

The SATSA is a twin cohort that aims to understand individual differences in aging. It is a part of the Swedish Twin Registry [108], which is a population-based national register including twins born between 1886 and 2000. SATSA includes 2,018 same sex twin individuals measured by nine questionnaire and ten in-person testing (IPT) waves from 1984 to 2014. The data collection and sampling procedures have been previously described [109, 110]. DNA material was obtained from whole blood longitudinally but only baseline measures were used in the analysis. For each twin pair, one twin was randomly selected so that participants were independent in the analysis. After selection, the age of participants ranged from 48 to 94 with a mean of 69 years.

*Ethics*

All participants in SATSA have provided written informed consents. This study was approved by the ethics committee at Karolinska Institutet with Dnr 2015/1729-31/5.

*DNA Methylation*

DNA methylation data used in the analysis were measured from whole blood using the Illumina 450k array for 203 participants and the Illumina EPIC array for 77 participants. Full details on quality control have been reported previously [111]. Briefly, probes were filtered based on three criteria: overlap a SNP in probe sequence; have a detection p-value >0.05 in any sample and locate on sex chromosomes. Samples were removed if predicted sex did not match reported sex, or if the correlation between sample mix-up probes and measured genotypes was lower than 0.7. These data were then uploaded to Horvath’s online calculator, as described in the main text, to generate the various clock estimates.

*Genotyping, imputation and quality control*

Genotyping of SATSA participants was carried out using the Illumina PsychChip. Genotype quality control details have been described previously [111], which included the removal of samples with >1% missing genotypes, estimated inbreeding coefficient >3SD from sample mean, wrong relatedness and wrong predicted sex; SNPs not mapped to a chromosome, with a low call rate (<98%), with a Hardy-Weinberg P-value<1x10^-6^ and with no observed minor alleles. The genotype data after QC were imputed to the 1000 Genomes Project phase 1 [42]. Further QC was carried out to remove SNPs of low imputation quality Info<0.6 and MAF<0.05, leaving an imputed dataset of over 6.5 million variants for analysis. The GWAS was run using PLINK v1.90 [6].

*Data availability*

The DNA methylation data of the SATSA study are archived in ArrayExpress at EMBL-EBI under accession number E-MTAB-7309. Please contact for further request.

*Acknowledgements*

Funding

The SATSA study was supported by NIH grants R01 (AG04563, AG10175, AG028555), the MacArthur Foundation Research Network on Successful Aging, the Swedish Council for Working Life and Social Research (FAS/FORTE) (97:0147:1B, 2009-0795, 2013-2292), and the Swedish Research Council (825-2007-7460, 825-2009-6141, 521-2013-8689, 2015-03255, 2015-06796), the Loo & Hans Osterman Foundation, the Foundation for Geriatric Diseases, the Magnus Bergwall Foundation, the Karolinska Institutet delfinansiering (KID) grant for doctoral students, the Strategic Research Program in Epidemiology at Karolinska Institutet, King Gustaf V:s and Queen Victorias Freemason Foundation, and Erik Rönnbergs donation for scientific studies in aging and age-related diseases.

**Sister Study (SS)**

*Cohort description*

The Sister Study is a prospective cohort study of 50,884 women recruited from the United States and Puerto Rico between 2003 and 2009 [112]. To be eligible, women had to be age 35-75 years, could not have had breast cancer themselves, but must have had a biological sister with breast cancer. Participants provided extensive information at baseline interview. Women are re-contacted annually for information on breast cancer and other basic health information and every 2-3 years for more detailed follow-up, with ~95% response rates for annual updates. The methylation case–cohort study [113] included a random sample of 1,336 non-Hispanic white women (74 of whom were later diagnosed with breast cancer) drawn from the full cohort, and an additional 1,542 non-Hispanic white women diagnosed with incident breast cancer during the time between enrolment and July 2014 (Data Release 7.0). DNA was obtained from whole blood at the study baseline.

*Ethics*

Written informed consent were obtained during a home visit. The study was conducted in accordance with recognized ethical guidelines and approved by the institutional review boards of the National Institute of Environmental Health Sciences (NIEHS), National Institutes of Health (NIH), and the Copernicus Group.

*DNA Methylation*

DNA methylation profile in blood was assessed using Illumina Human450 Methylation Arrays. We excluded samples due to data quality issues: average bisulfite intensity < 4000 or had more than 5% of probes with low-quality methylation values (detection p >0.000001, number of beads <3, or values outside of 3IQR); samples that were outliers for their methylation beta value distributions. Methylation data preprocessing and quality control was completed using the *ENmix* R software package [49], including the following steps: ENmix background correction [49], RELIC dye bias correction [114], quantile normalization and RCP [51] probe design bias adjustment. These data were then uploaded to Horvath’s online calculator, as described in the main text, to generate the various clock estimates.

*Genotyping and quality control*

Sister Study genotyping was carried out with 3,030 samples genotyped using the Illumina OncoArray Bead Chip [115], which has probes for genotyping of 533,000 SNPs. Approximately 50% of the markers were selected as a GWAS backbone to tag the majority of known common variants. The remaining markers were selected for various cancer types. Samples and SNPs with low quality data were excluded as following: we first exclude samples with call rate <80%; then excluded SNPs with call rate <80%; and we then excluded samples with call rate <95%; finally, we excluded SNPs with call rate <95%. Hardy-Weinberg equilibrium were checked and SNPs were excluded if P<10^-7^.

*Data availability*

Data from the Sister Study can be requested via https://sisterstudy.niehs.nih.gov/English/coll-data.htm.

*Acknowledgements*

This work was supported by Intramural Research Program of the NIH, National Institute of Environmental Health Sciences (Z01 ES049033, Z01 ES049032, Z01 ES044005).

**TwinsUK**

*Cohort description*

The UK Adult Twin Registry (TwinsUK) registry was established in 1992 to recruit same-sex monozygotic and dizygotic twins from the UK. Over 80% of participants are female and age range is from 16 to 98 years old). The cohort includes more than 14,000 twin participants from all regions across the United Kingdom. Research twin volunteers have contributed longitudinal questionnaire data and attended clinical visits where many twins have had multiple visits over the years. The TwinsUK cohort also includes collection of different biological samples, as well as biological data profiles. The TwinsUK cohort has been used in many epidemiological studies and is representative of the general UK population for a wide range of diseases and traits [116].

*Ethics*

Ethical approval was granted by the National Research Ethics Service London-Westminster, the St Thomas’ Hospital Research Ethics Committee (EC04/015 and 07/H0802/84). All twins provided written informed consent prior to taking part in research activities.

*DNA Methylation*

DNA methylation profiles generated using the Illumina 450K array in whole blood samples in TwinsUK have previously been described [117, 118, 119]. Briefly, DNA samples were extracted from whole blood using DNeasy kit (Qiagen, Inc) and bisulfite converted using EZ DNA methylation kit (Zymo Research Corporation). DNA methylation levels were detected using the Illumina Infinium HumanMethylation450 array (Illumina) and methylation betas were generated using the R package minfi with background correction. Raw beta levels were first applied the beta mixture quantile dilation (BMIQ) method. Probe exclusion criteria including probes mapped to multiple locations to the reference sequence, and probes where more than 1% of subjects had detection p-value > 0.05. Individuals with over 5% missing probes, with mismatched sex, and with mismatched genotypes were excluded. This study was carried out in a subset of 496 unrelated subjects.

*Genotyping, imputation and quality control*

The larger TwinsUK dataset was genotyped using a combination of Illumina HumanHap300, HumanHap610Q, HumanHap1M Duo, and HumanHap1.2M Duo 1M arrays. Separate imputations were performed for the datasets and then merged using GTOOL. Genotype data were pre-phased using IMPUTE2 without reference panels. The resulting haplotypes underwent fast imputation using the 1000 genome PHASE1 data set. The reference set using 1000 Genomes Phase I (interim) was based on sequence data freeze from November 23, 2010, and the phased haplotype data release in June 2011. After imputation, SNPs were filtered at a MAF>5%.

*Data availability*

The majority of TwinsUK DNA methylation data are available through the NCBI Gene Expression Omnibus (GEO) under accession numbers GSE62992 and GSE121633. Individual level genotype, DNA methylation, and phenotype data can be accessed through application to the TwinsUK data access committee (https://twinsuk.ac.uk/resources-for-researchers/access-our-data/).

*Acknowledgments*

The TwinsUK study was funded by the Wellcome Trust and European Community’s Seventh Framework Programme (FP7/2007-2013). The TwinsUK study also receives support from the National Institute for Health Research (NIHR)-funded BioResource, Clinical Research Facility and Biomedical Research Centre based at Guy’s and St Thomas’ NHS Foundation Trust in partnership with King’s College London. This research study was also supported by a joint UK Economic and Social Research Council (ESRC) and Biotechnology and Biological Sciences Research Council (BBSRC) grant (ES/N000404/1 to J.T.B.), and funding support from the JPI ERA-HDHL DIMENSION project (BBSRC BB/S020845/1 to J.T.B).

**The Cardiovascular Risk in Young Finns Study (YFS)**

*Cohort description*

YFS is one of the largest existing follow-up studies into cardiovascular health from childhood to adulthood, running in a longitudinal prospective setup with regular follow-ups from 1980 onwards [120, 121]. The study began in 1980 with 3596 children and adolescents aged 3 to 18 years randomly selected from the areas of five university hospitals in Finland (Turku, Tampere, Helsinki, Kuopio and Oulu). The participants have been followed up for over 40 years. The present study is based on subsample of 1402 participants from the 2011 follow-up. DNA was obtained from whole blood samples.

*Ethics*

The study followed the guidelines of the Declaration of Helsinki and was approved by the ethical review committee of the Hospital district of Southwest Finland and the Regional Ethics Committee of the Expert Responsibility area of Tampere University Hospital. All participants submitted informed consent to participate.

*DNA Methylation*

Leukocyte DNA of the YFS cohort was obtained from EDTA-blood samples using a Wizard® Genomic DNA Purification Kit (Promega Corporation, Madison, WI, USA) according to the manufacturer’s instructions. Genome-wide DNA methylation levels were obtained using Infinium MethylationEPIC array according to the protocol by Illumina. All the analysed samples have sum of detection P-values across all the probes less than 0.01. Logged (log2) median of methylated and unmethylated intensities of the analysed samples clustered visually well. Further, samples for which real sex did not match the predicted sex were excluded. Background subtraction and dye-bias normalization was performed via noob method [13] followed by stratified quantile normalization. Probes with detection p-value more than 0.01 in 99% of the samples were filtered out. These data were then uploaded to Horvath’s online calculator, as described in the main text, to generate the various clock estimates.

*Genotyping, imputation and quality control*

Genomic DNA was extracted from peripheral blood leukocytes using a commercially available kit and Qiagen BioRobot M48 Workstation according to the manufacturer’s instructions (Qiagen, Hilden, Germany). Genotyping was done for 2556 samples using custom build Illumina Human 670 k BeadChip atWelcome Trust Sanger Institute. Genotypes were called using Illuminus clustering algorithm. 56 samples failed Sanger genotyping pipeline QC criteria (i.e., duplicated samples, heterozygosity, low call rate, or Sequenom fingerprint discrepancy). From the remaining 2500 samples one sample failed gender check, three were removed due to low genotyping call rate (< 0.95) and 54 samples for possible relatedness (pi-hat > 0.2). 11766 Short Nucleotide Polymorphisms (SNPs) were excluded based on Hardy-Weinberg equilibrium test (p ≤ 1e-06), 7746 SNPs failed missingness test (call rate < 0.95) and 34596 SNPs failed frequency test (minor allele frequency < 0.01). After quality control there were 2442 samples and 546677 genotyped SNPs available for further analysis. Genotype imputation was performed using Minimac3 [107] and 1000G phase3 reference set on Michigan Imputation Server. Autosomes and sex chromosomes were phased using Eagle [3] and SHAPEIT [4] respectively.

*Data availability*

The YFS dataset comprises health related participant data and their use is therefore restricted under the regulations on professional secrecy (Act on the Openness of Government Activities, 612/1999) and on sensitive personal data (Personal Data Act, 523/1999, implementing the EU data protection directive 95/46/EC). Due to these legal restrictions, the Ethics Committee of the Hospital District of Southwest Finland has in 2016 stated that individual level data cannot be stored in public repositories or otherwise made publicly available. Data sharing outside the group is done in collaboration with YFS group and requires a data-sharing agreement with the understanding that collaborators will protect the data and not share it with any other parties. The list of all investigators that collaborate with the YFS group is displayed at the website of the YFS (http://youngfinnsstudy.utu.fi/). Investigators can submit an expression of interest to the chairman of the data sharing and publication committee (Prof Mika Kähönen, Tampere University, Finland).

*Acknowledgements*

The Young Finns Study has been financially supported by the Academy of Finland: grants 322098, 286284, 134309 (Eye), 126925, 121584, 124282, 129378 (Salve), 117787 (Gendi), and 41071 (Skidi); the Social Insurance Institution of Finland; Competitive State Research Financing of the Expert Responsibility area of Kuopio, Tampere and Turku University Hospitals (grant X51001); Juho Vainio Foundation; Paavo Nurmi Foundation; Finnish Foundation for Cardiovascular Research ; Finnish Cultural Foundation; The Sigrid Juselius Foundation; Tampere Tuberculosis Foundation; Emil Aaltonen Foundation; Yrjö Jahnsson Foundation; Signe and Ane Gyllenberg Foundation; Diabetes Research Foundation of Finnish Diabetes Association; EU Horizon 2020 (grant 755320 for TAXINOMISIS; grant 848146 for To_Aition); European Research Council (grant 742927 for MULTIEPIGEN project); and Tampere University Hospital Supporting Foundation.

**NHLBI Trans-Omics for Precision Medicine (TOPMed) Consortium**

The Genetics of Lipid Lowering Drugs and Diet Network Study (GOLDN) and The Multi-Ethnic Study of Atherosclerosis (MESA) cohorts received Omics support from the TOPMed consortium.

| TOPMed Omics Support Table | | | | | | |
| --- | --- | --- | --- | --- | --- | --- |
| **TOPMed Accession #** | **TOPMed Project** | **Parent Study** | **TOPMed Phase** | **Omics Center** | **Omics Support** | **Omics Type** |
| **phs001359** | GOLDN | GOLDN | 2 | NWGC | 3R01HL104135-04S1 | WGS |
| **phs001416** | MESA | MESA | pilot | Keck MGC | HHSN268201600034I | Methylomics |
