## Appendix 2: Heterogeneity Analyses for "Genome-wide association studies identify 137 loci for DNA methylation biomarkers of ageing"

**Appendix 2: Heterogeneity Analysis**

We found little evidence of systematic between-study heterogeneity. In the European-ancestries meta-analysis, the Sister study was identified as an influential M-Statistic outlier (P_Bonferroni_<0.05) for the GWAS of PhenoAge acceleration. Re-running the respective meta-analyses without this cohort made negligible differences to the meta-analysis output for the genome-wide significant SNPs (**Extended Data Figures 1-3**). Meta-regression analysis of the M-Statistics on mean and SD of cohort age, proportion of females, sample size, genomic inflation, SNP imputation reference panel (HRC or 1000G), cohort region (Europe, USA, Australia), and N_SNPs_ contributed to the meta-analysis was carried out for each trait, with no significant (P <0.05) associations between study characteristics and cohort-level heterogeneity.

The GENOA study was identified as an influential M-statistic outlier in the African-ancestries analysis of Granulocyte proportions. As was the case with the Sister study and PhenoAge acceleration, removal of GENOA from the meta-analysis made little difference to the output for the genome-wide significant SNPs (**Extended Data Figures 4-12**). Meta-regression analysis was performed as above. There was a significant relationship between the proportion of female samples and cohort-level heterogeneity in the analysis of IEAA (Estimate=0.24; P = 0.035)

There was relatively low evidence of heterogeneity for the top findings in either the European-ancestries or African-ancestries populations, based on the heterogeneity I^2^ statistic. However, it is more pronounced in the African-ancestries populations with 30% of loci with P<5e-8 (across all traits) having I^2^ >40% compared to 16% in European-ancestries (**Extended Data Figure 13)**. This is likely to be due to the small number of African-ancestries studies (N=7) compared to European-ancestries studies (N=28). Contour plots of I2 versus P-value are presented in **Extended Data Figures 14-25**.

We filtered the African-ancestries SNPs to those with I^2^ <40% and recalculated lambda finding little difference to lambda calculated on the full set of SNPs (**Extended Data Table 1)**. The results suggest heterogeneity is not contributing to the observed inflation. Furthermore, inflation was observed across all allele frequency groupings in the African-ancestries analysis, and was slightly more pronounced in low-frequency variants for all traits with the exception of IEAA (**Extended Data Figure 26**). There was greater variability in effect sizes across African-ancestries analyses compared to effect sizes in the European-ancestries analyses (**Extended Data Figure 27**).

**Extended Data Table 1:**

|  | IEAA | Hannum | GrimAge | Gran | PhenoAge | PAI1 |
| --- | --- | --- | --- | --- | --- | --- |
| **ALL** | 1.15 | 1.14 | 1.21 | 1.11 | 1.19 | 1.17 |
| **I^2^ < 40** | 1.15 | 1.14 | 1.2 | 1.11 | 1.19 | 1.17 |

**Extended Data Figure 1:** Cohort-level and meta-analysis effect sizes for PhenoAge acceleration at marker 17:3378876 with (Meta) and without the Sister study (Meta_PostHet).

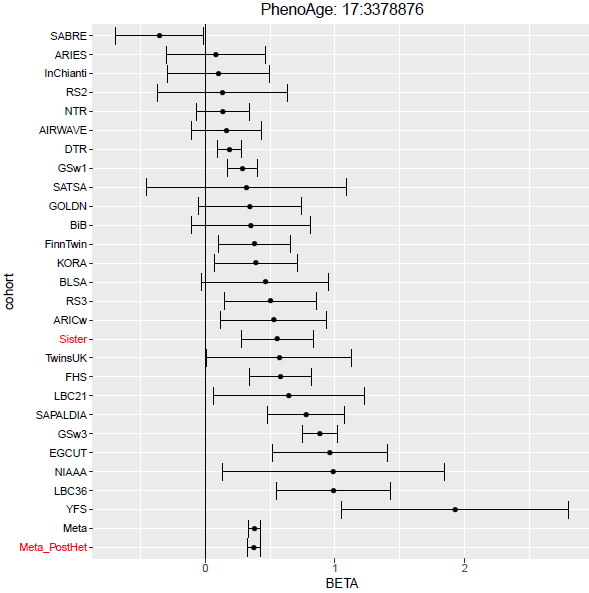

**Extended Data Figure 2:** Cohort-level and meta-analysis effect sizes for PhenoAge acceleration at marker 6:18106076 with (Meta) and without the Sister study (Meta_PostHet).

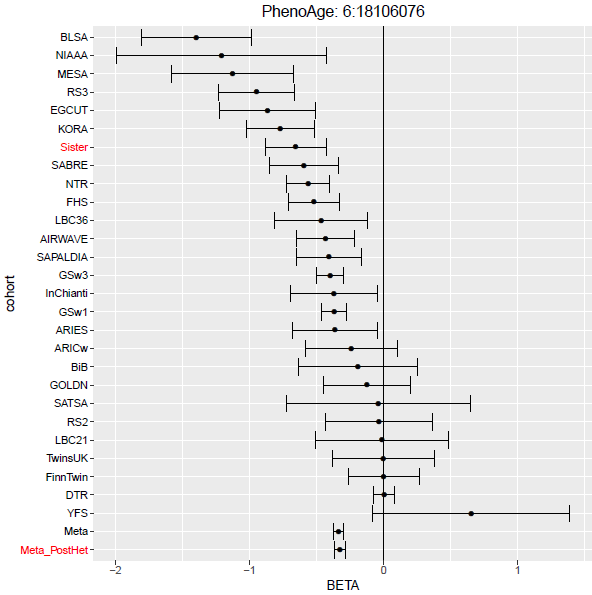

**Extended Data Figure 3:** Cohort-level and meta-analysis effect sizes for PhenoAge acceleration at marker 7:44925896 with (Meta) and without the Siste r study (Meta_PostHet).

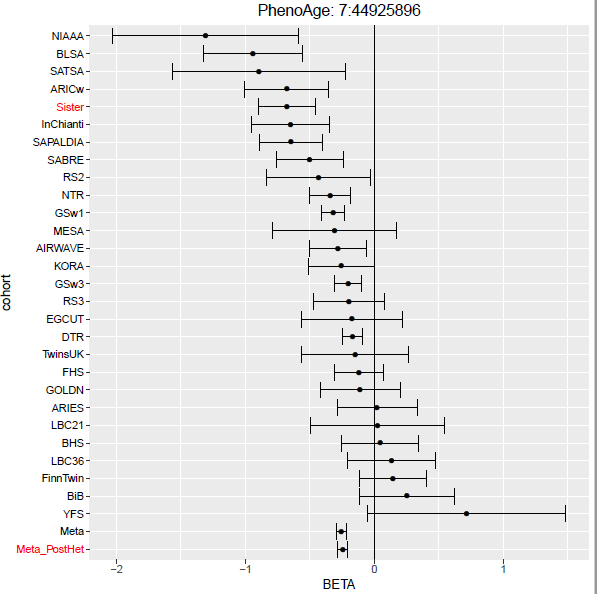

**Extended Data Figure 4:** Cohort-level and meta-analysis effect sizes for Granulocyte proportions at marker 1:159223787 with (Meta) and without the GENOA study (Meta_PostHet).

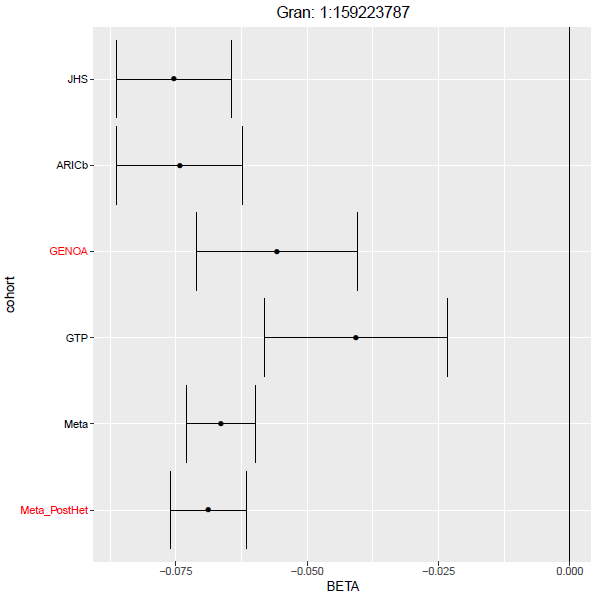

**Extended Data Figure 5:** Cohort-level and meta-analysis effect sizes for Granulocyte proportions at marker 1:159237950 with (Meta) and without the GENOA study (Meta_PostHet).

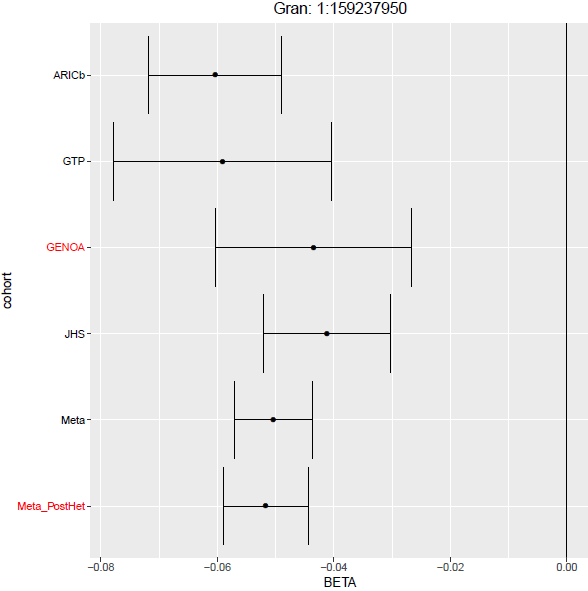

**Extended Data Figure 6:** Cohort-level and meta-analysis effect sizes for Granulocyte proportions at marker 1:159247238 with (Meta) and without the GENOA study (Meta_PostHet).

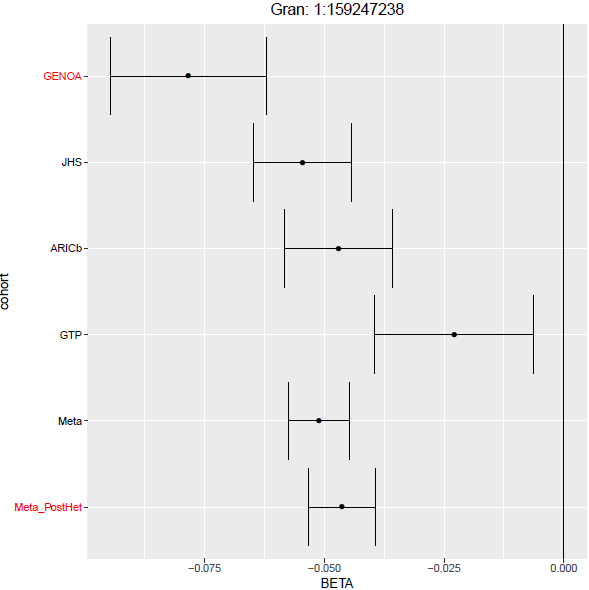

**Extended Data Figure 7:** Cohort-level and meta-analysis effect sizes for Granulocyte proportions at marker 1:159558258 with (Meta) and without the GENOA study (Meta_PostHet).

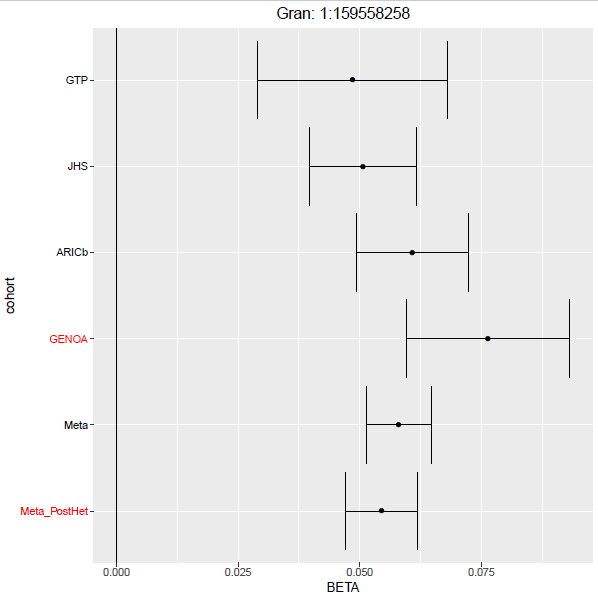

**Extended Data Figure 8:** Cohort-level and meta-analysis effect sizes for Granulocyte proportions at marker 1:159608855 with (Meta) and without the GENOA study (Meta_PostHet).

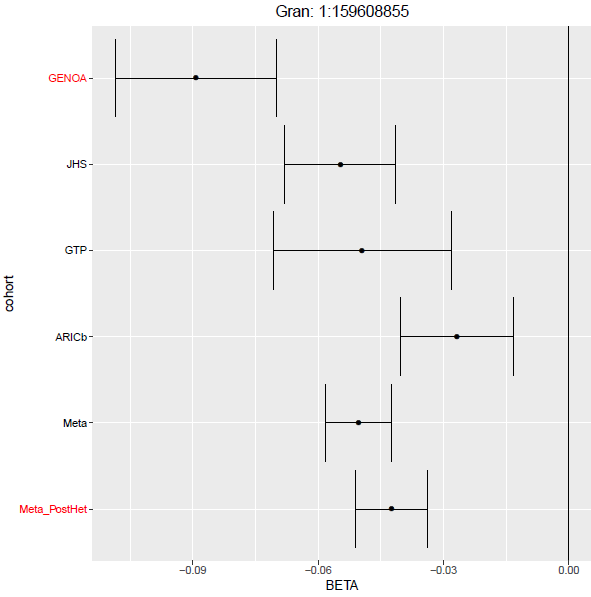

**Extended Data Figure 9:** Cohort-level and meta-analysis effect sizes for Granulocyte proportions at marker 1:160046447 with (Meta) and without the GENOA study (Meta_PostHet).

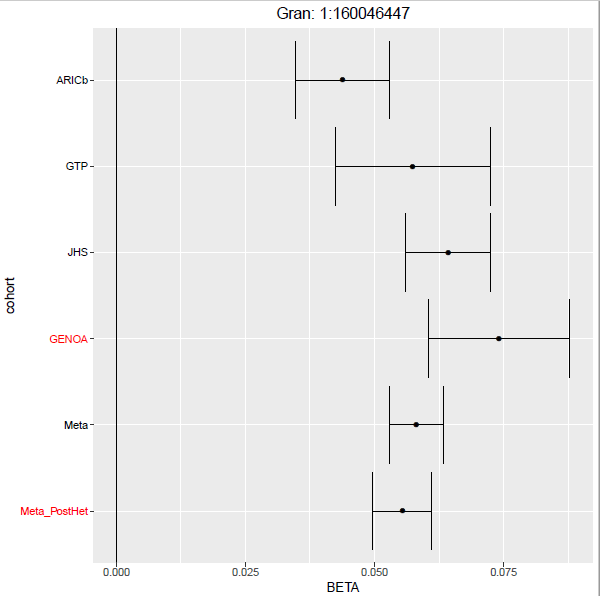

**Extended Data Figure 10:** Cohort-level and meta-analysis effect sizes for Granulocyte proportions at marker 1:160053342 with (Meta) and without the GENOA study (Meta_PostHet).

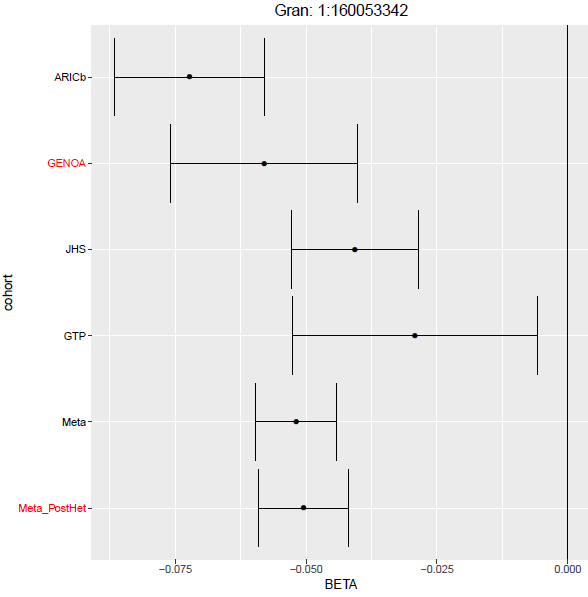

**Extended Data Figure 11:** Cohort-level and meta-analysis effect sizes for Granulocyte proportions at marker 1:160908841 with (Meta) and without the GENOA study (Meta_PostHet).

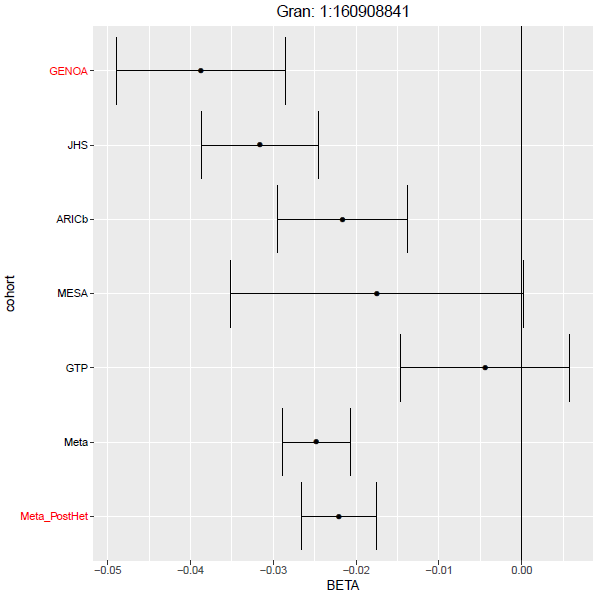

**Extended Data Figure 12:** Cohort-level and meta-analysis effect sizes for Granulocyte proportions at marker 1:163590534 with (Meta) and without the GENOA study (Meta_PostHet).

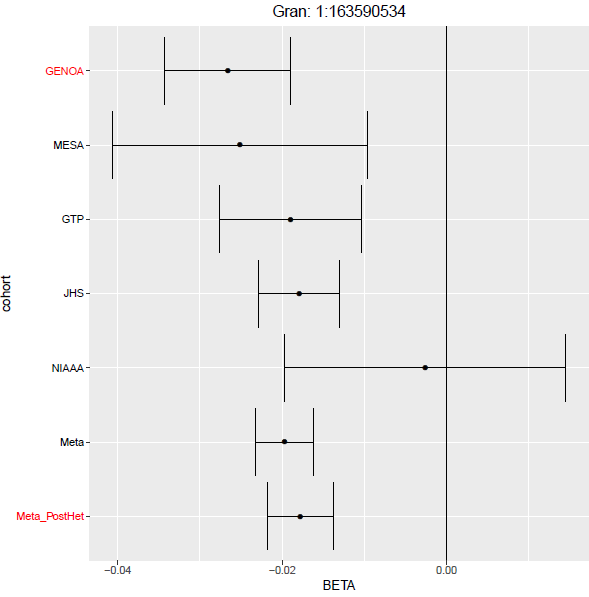

**Extended Data Figure 13:** I^2^ statisitc distribution for SNPs at P < 5 x 10^-8^ in European-ancestries (left) and African-ancestries analyses (right)

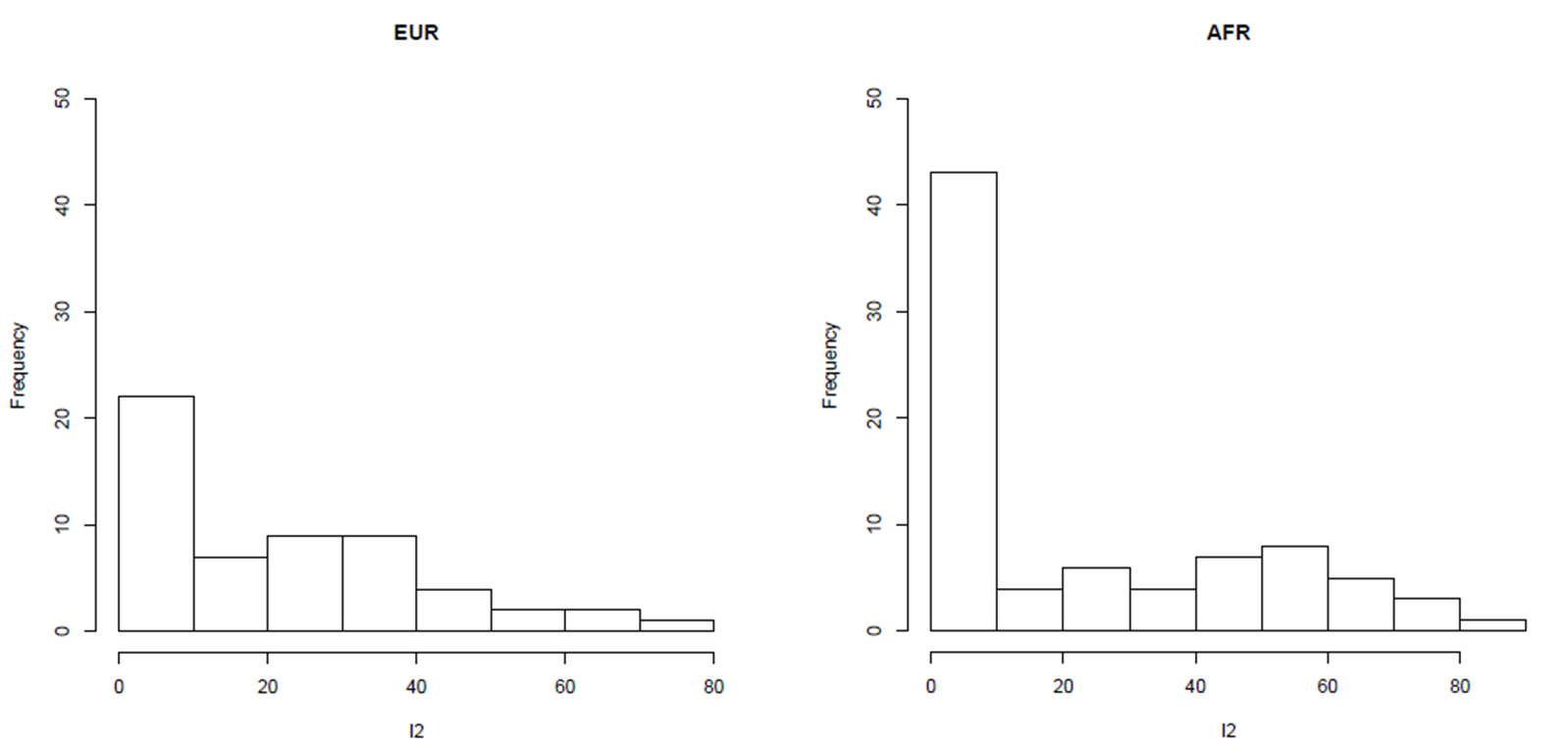

**Extended Data Figure 14:** I^2^ statistic vs –log10(P-value) for IEAA (European Ancestries analysis)

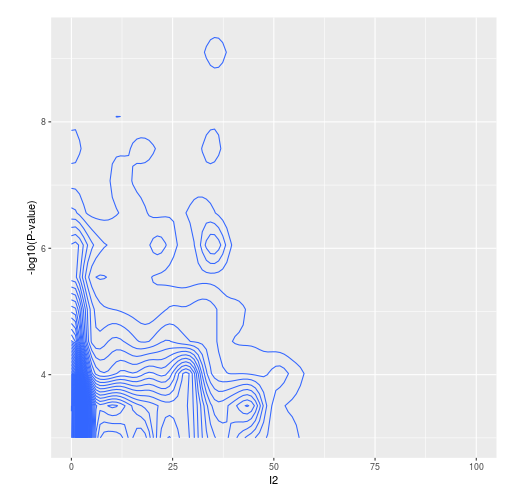

**Extended Data Figure 15:** I^2^ statistic vs –log10(P-value) for PhenoAge acceleration (European Ancestries analysis)

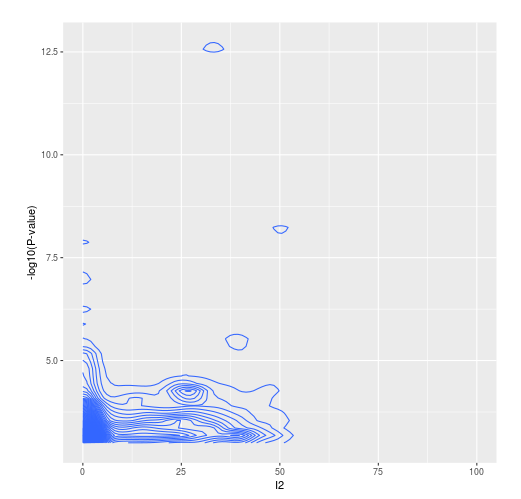

**Extended Data Figure 16:** I^2^ statistic vs –log10(P-value) for GrimAge acceleration (European Ancestries analysis)

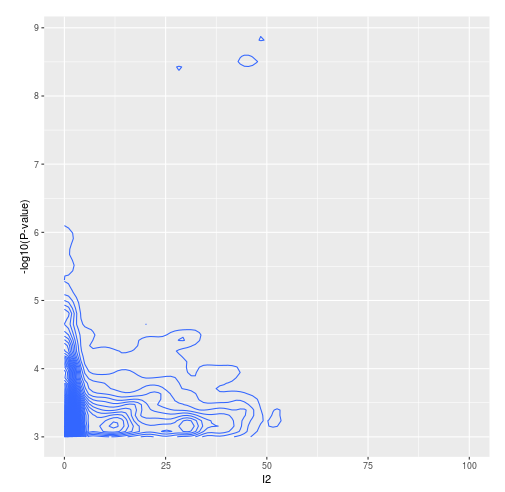

**Extended Data Figure 17:** I^2^ statistic vs –log10(P-value) for Granulocyte proportions (European Ancestries analysis)

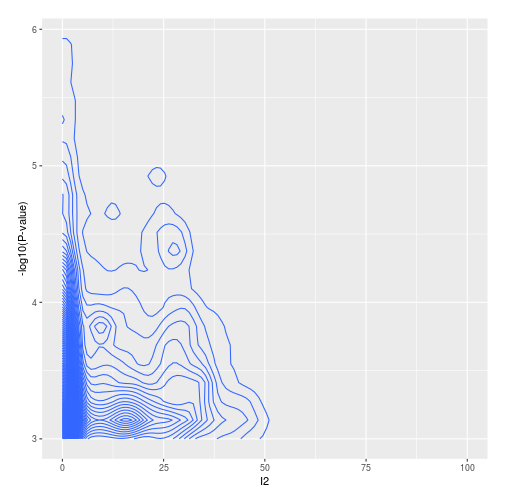

**Extended Data Figure 18:** I^2^ statistic vs –log10(P-value) for Hannum age acceleration (European Ancestries analysis)

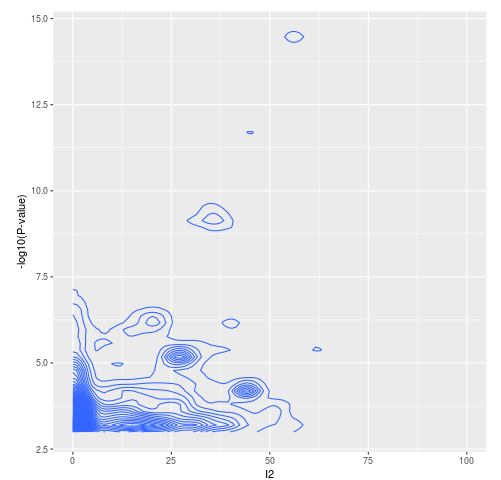

**Extended Data Figure 19:** I^2^ statistic vs –log10(P-value) for PAI1 levels (European Ancestries analysis)

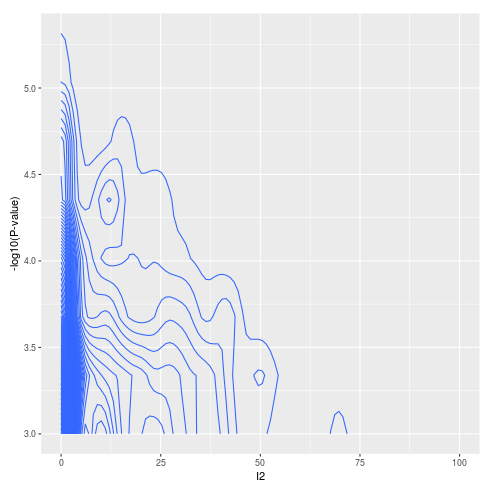

**Extended Data Figure 20:** I^2^ statistic vs –log10(P-value) for IEAA (African Ancestries analysis)

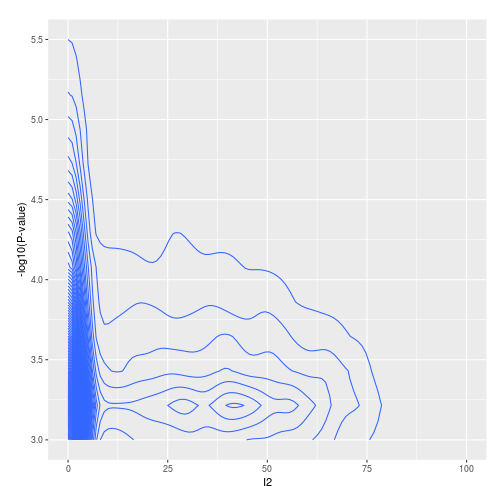

**Extended Data Figure 21:** I^2^ statistic vs –log10(P-value) for PhenoAge acceleration (African Ancestries analysis)

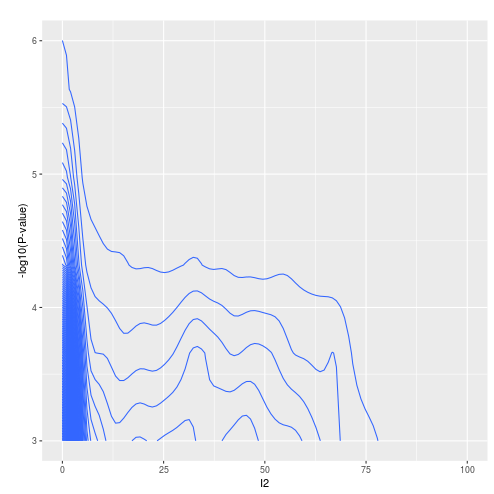

**Extended Data Figure 22:** I^2^ statistic vs –log10(P-value) for GrimAge acceleration (African Ancestries analysis)

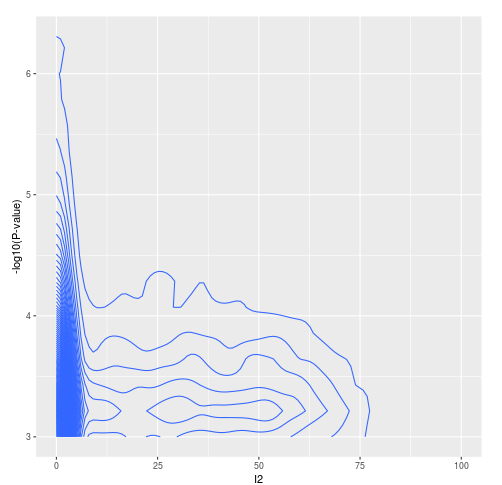

**Extended Data Figure 23:** I^2^ statistic vs –log10(P-value) for Granulocyte proportions (African Ancestries analysis)

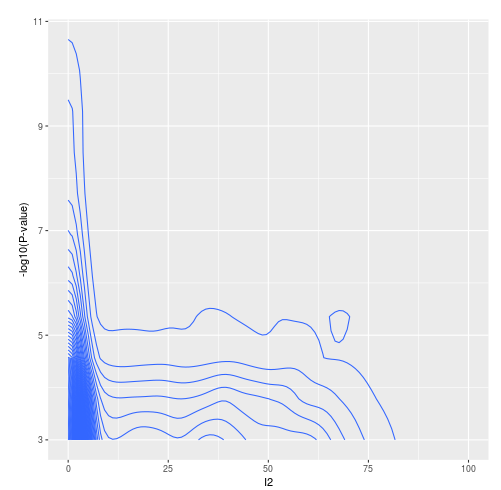

**Extended Data Figure 24:** I^2^ statistic vs –log10(P-value) for Hannum age acceleration (African Ancestries analysis)

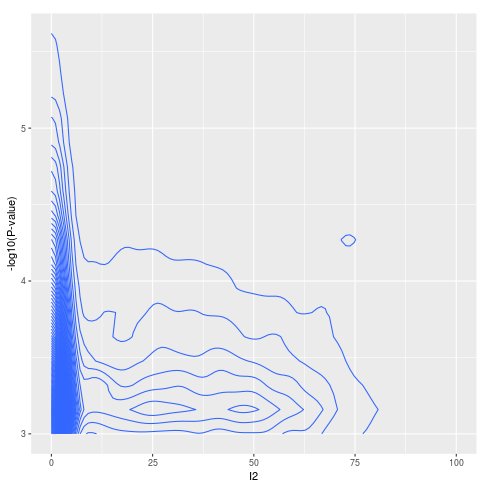

**Extended Data Figure 25:** I^2^ statistic vs –log10(P-value) for PAI1 levels (African Ancestries analysis)

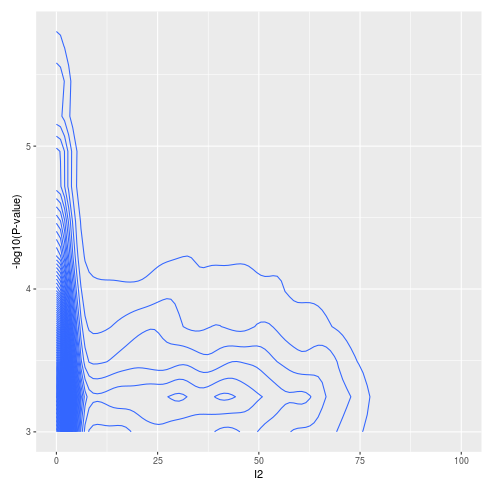

**Extended Data Figure 26:** Genomic inflation by minor allele frequency grouping in the African-ancestries meta-analyses. The red horizontal line indicates genomic inflation when including all variants.

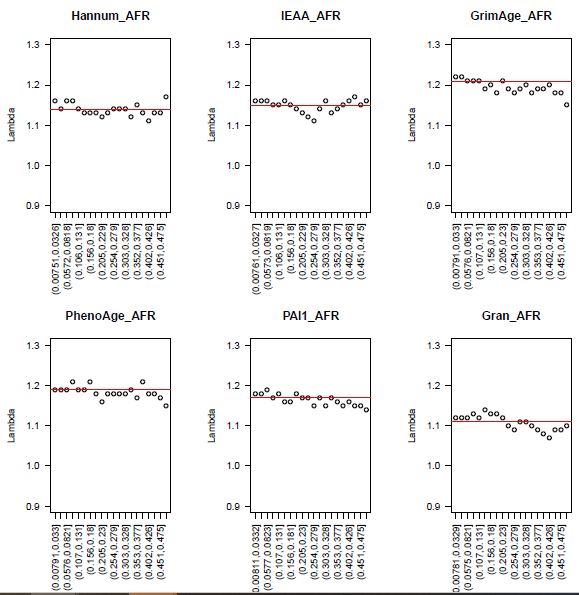

**Extended Data Figure 27:** Effect sizes for all SNPs present in both European-ancestries and African-ancestries meta-analyses.

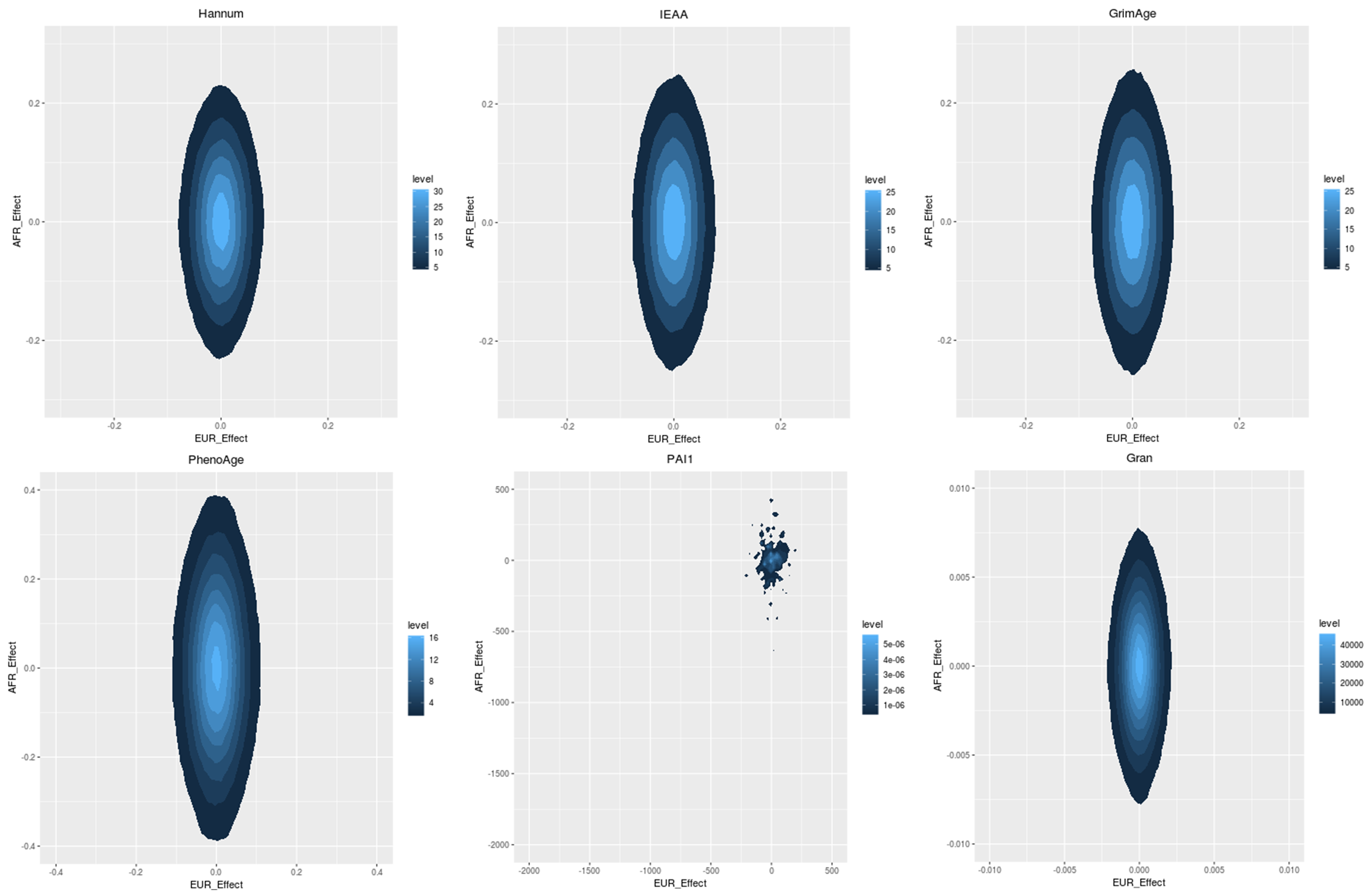
